# Topological Complementarity Predicts Synergistic Drug Pairs in Branched Metabolic Networks

**DOI:** 10.64898/2026.10.24.761489

**Authors:** Deeptanshu Pandey, Purbita Sikdar, Aniruddha Chattaraj, Sourav Chowdhury

**Affiliations:** Department of Biological Sciences, Birla Institute of Technology and Science-Pilani, Hyderabad, India; Max Plank Institute for Dynamics and Self Organization, Göttingen, Germany

**Author notes:** **Correspondence: Sourav Chowdhury**, Department of Biological Sciences, Birla Institute of Technology and Science, BITS-Pilani Hyderabad Campus, Hyderabad, Telangana, India, 500078, **Aniruddha Chattaraj** Max Plank Institute for Dynamics and Self Organization, Göttingen, Germany.

**Keywords:** Drug synergy, Metabolic network topology, Folate metabolism, Kinetic modeling, Antifolate combinations, Metabolic compensation

## Abstract

Drug discovery typically ranks targets by individual sensitivity. Branched metabolic networks, in contrast, have redundant flux routes that buffer single-enzyme inhibition possibly deviating the best target from the most sensitive node. We developed a topology-driven framework for identifying optimal dual-target pairs and applied it to the E. coli folate pathway. A kinetic model of the folate pathway and its extended one-carbon pool showed that single-node inhibition is compensated by flux redistribution. Pairwise synergy scoring, a metric to measure drug-pair potency as compared to single drug, identified DHPS (folP) and DHFR (folA) as a top-ranked complementary pair. Experimental validation in E. coli MG1655 using sulfamethoxazole and trimethoprim confirmed strong dose-dependent synergy at intermediate inhibition and attenuation at saturation. Our results support topological complementarity as a key predictor of combination efficacy and provide a rationale for drug-pair design in branched biosynthetic pathways.

## Introduction

The bacterial folate pathway has been an important antimicrobial target for more than half a century, yet therapeutic intervention has focused on only a small number of enzymes within a much larger metabolic network. Because the pathway is branched, it does not respond to inhibition in a simple, proportional manner. Consequently, the effect of inhibiting a single enzyme depends on both its kinetic properties and on flux redistribution through the surrounding network, and this buffering currently explains why single-target inhibition is often insufficient to predict optimal dual-target pairs. We develop a topology-driven approach for the E. coli folate pathway, use it to identify the optimal pair of enzymes for dual targeting, and validate the prediction experimentally.

Inhibiting a single enzyme in any metabolic network with redundant routes does not shut down the entire pathway. We therefore develop a framework, not limited to folate metabolism, that chooses drug targets in a branched pathway [1-3]. Metabolic control analysis showed that control is shared among several reactions rather than residing in one rate-limiting step [1,2]. Flux control coefficients sum to unity. No single enzyme dominates flux, and most enzymes exert only a small effect on pathway output [1]. The control matrix extends this to metabolite concentrations, showing that sensitivity is distributed across the network [2]. Control can be calculated from network connectivity and reaction kinetics without perturbing each enzyme individually [4]. Supply–demand analysis formalized how control is distributed between upstream supply blocks and downstream demand blocks [4]. When one block controls the flux, it loses influence over the linking metabolite maintaining which becomes the sole function of the other block [4]. Blocking one route therefore shifts control to the remaining routes, and the system compensates through remaining open routes. This motivates the search for combinations that block alternative routes as single-target inhibition tends to be insufficient. Network pharmacology formalizes this idea: the optimal drug pair is defined by topology, not by rank order of individual targets [3,5]. Yet, quantitative topology-driven frameworks for identifying such pairs in bacterial biosynthetic pathways remain scarce.

Bacterial folate metabolism can be used as a test case for such a framework. Folate drives one-carbon transfer reactions that sustain nucleotide, amino acid, and cofactor biosynthesis, and it remains a validated antimicrobial target in Escherichia coli and other pathogens [6]. The pathway is branched. DHNTP diverts into folate biosynthesis and a tetrahydromonapterin shunt that produces more pterin than the folate branch itself [7,8]. THF feeds three separate biosynthetic routes: thymidylate, purine, and methionine. Salvage and recycling reactions maintain intracellular folate levels through additional routes [9,10]. When one route is blocked, flux redistributes through the remaining branches.

The pathway is also physically organized. A map of 1225 protein-protein interactions across E. coli one-carbon metabolism found that folate biosynthesis enzymes interact strongly with each other and with purine biosynthesis enzymes [11]. FolA (encoded by folA), the TMP target, binds GlyA (encoded by glyA) and PurH (encoded by purH) through surfaces distinct from its active site, forming a physical enzyme cluster [11]. Within this cluster, the product of one enzyme passes directly to the next rather than diffusing through bulk cytoplasm [11]. When a drug inhibits one enzyme in the cluster, the remaining enzymes depend on the clustered partner that is now blocked. They cannot recruit free intermediates from solution to compensate. Sulfamethoxazole (SMX) and trimethoprim (TMP) are two well-characterized drugs used in combination that target this pathway, inhibiting DHPS (encoded by folP) and DHFR (encoded by folA) respectively [6,12,13]. SMX blocks production of the THF precursor dihydropteroate. TMP blocks conversion of dihydrofolate (DHF) to THF. SMX potentiates TMP by limiting de novo DHF production. This mono-potentiation mechanism is the current explanation for their synergistic action [13]. Mathematical modeling of this pathway explains how SMX potentiates TMP. It does not explain how TMP potentiates SMX [13]. No single enzyme inhibition produces a simple, proportional response in pathway flux.

We then evaluated pairwise synergy by combining two nodes across the network. Individual sensitivity indices of the two nodes were integrated with a topological weighting. This weighting penalizes pairs acting on the same linear segment and rewards pairs that close distinct compensatory routes. The weighting reflects a general principle. Enzymes that are functionally coupled but topologically separable can co-evolve as adaptive units. Comparative genomics and experimental evolution in E. coli identified one such unit: DHFR and thymidylate synthase (TYMS, encoded by thyA) [14]. This unit responds to antibiotic stress relatively independently from the rest of folate metabolism. Metabolomic profiling showed that TYMS activity must not exceed DHFR activity. If it does, reduced folates are depleted and dihydrofolate accumulates [14]. This creates a co-evolutionary dependency. Perturbing one enzyme shifts selection on its partner.

The same logic extends to DHPS and DHFR. DHPS gates entry to the committed de novo branch. DHFR gates THF regeneration downstream. Blocking both closes the de novo route and removes the salvage-dependent rebound that sustains flux under single-enzyme inhibition. DHPS and DHFR are also physically proximal within the clustered folate metabolon. Blocking both disrupts the branch point and the downstream regeneration step within the same enzyme assembly. No other folate-specific pair closes both routes simultaneously.

We tested this prediction experimentally in E. coli MG1655. We exposed the strain to SMX and TMP individually and in combination across a two-fold dilution series in minimal medium. We evaluated drug interactions using four complementary metrics [15-18]. Bliss independence tests whether the combined effect equals the product of the individual effects [15]. Chou– Talalay combination indices classify the interaction across the full dose-response curve, where a combined index below 1 indicates synergy, equal to 1 indicates additivity, and above 1 indicates antagonism [16]. Isobologram analysis maps combinations that produce equivalent effect at a fixed endpoint [17]. Comparison of combination AUC with a multiplicative null model compares total growth inhibition over time to what independent action would predict [18]. The network buffers single-node perturbation through salvage, competing, and recycling routes [9,10]. Simultaneous inhibition of DHPS and DHFR restricts several routes that sustain folate production and THF availability. Drug concentration determines the magnitude of interaction. Stronger-than-additive growth inhibition occurs when the network retains capacity for flux redistribution. The interaction weakens as inhibition approaches saturation. These findings explain the behaviour of the TMP-SMX combination.

## Results

(Central panels) THF interconverts reversibly with 5,10-methylene-THF, 5,10-methenyl-THF, and 10-formyl-THF (eco00670, one-carbon pool), and with 5-methyl-THF, which feeds the methyl cycle and regenerates methionine and THF. Boundary inputs (serine, glycine, dUMP, PABA) and outputs (dTMP, PABA, formate) are shown, along with the recycling reactions that close the folate cycle. The legend defines metabolite classes, enzyme notation, inhibition, and shunt pathways. (D) Full network representation of the model as a bipartite graph, showing 51 metabolite nodes and 39 reaction nodes. The graph collapses to 38 enzyme nodes for pairwise synergy scoring. Green nodes are metabolites; yellow nodes are reactions. The layout shows all connections of the folate, one-carbon, PreQ1, and MoCo modules, including boundary inputs and outputs.

### The folate network is scale-free, modular, and vulnerable at structurally distinct nodes

Folate biosynthesis in E. coli is one of three pterin pathways that share GTP as a precursor. The other two are PreQ1 biosynthesis and molybdenum cofactor (MoCo) biosynthesis (Figure 1A–C). The folate arm starts with folE which converts GTP to 7,8-dihydroneopterin triphosphate (DHNTP) [19]. DHNTP is then processed by nudB [7], phoA, folB, and folK to form HMDHP-PP [7,20]. The committed step is the condensation of HMDHP-PP with PABA to form DHP [21] by folP (DHPS), the target of sulfamethoxazole (SMX) [6]. DHP is converted by folC to DHF [22] which is reduced by folA (DHFR) to THF, the universal one-carbon carrier [23]. DHFR is the target of trimethoprim (TMP) [12]. THF feeds dTMP, purines, and methionine (Figure 1A). The pathway is not linear. A folX shunt diverts DHNTP to dihydromonapterin [8]. A salvage arm re-enters downstream of folE through folB and folK [9]. This layered topology determines how the pathway responds to inhibition. To study it we built the full bipartite network of the three GTP-consuming modules (90 nodes: 51 metabolites, 39 reactions; 96 edges).

**Figure 1:**
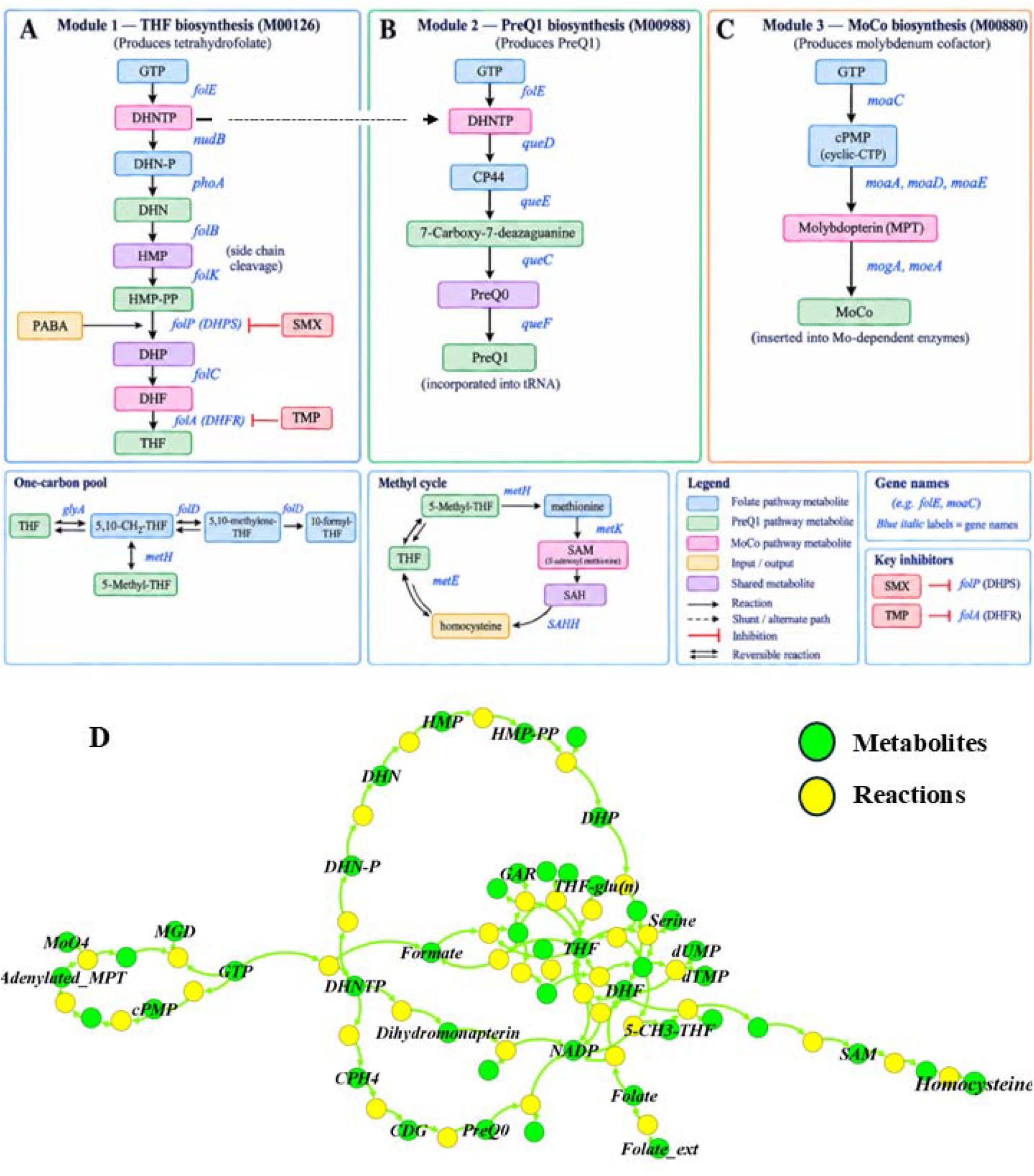
Architecture of the folate and one-carbon network in E. coli K-12 MG1655. (A– C) Three modules share GTP as a common precursor. (A) Module 1 (M00126, eco00790) builds tetrahydrofolate (THF) through nine enzymatic steps. The pathway forks at dihydroneopterin triphosphate (DHNTP): nudB directs flux to the folate branch, folX diverts it to the dead-end shunt dihydromonapterin. The folate branch proceeds through DHN-P, DHN, HMP, HMP-PP, dihydropteroate (DHP), and dihydrofolate (DHF) to THF. DHPS (folP) and DHFR (folA) are marked with red inhibitor arrows (TMP targets folA). (B) Module 2 (M00988) produces PreQ1 via CPH4, 7-carboxy-7-carbaguanine, and PreQ0. (C) Module 3 (M00880) produces the molybdenum cofactor via Cyclo-GTP, Precursor Z, and Molybdopterin.

Figure 1D shows this network as a bipartite graph: circles metabolites, squares reactions, and edges connecting each reaction to metabolites it consumes or produces. The folate arm runs along the left and centre of the graph. The PreQ□ and MoCo arms branch to the right. The folX shunt diverges near the top. The salvage arm re-enters from below. The one-carbon cycle and the methyl cycle appear at the bottom. Two nodes stand out visually. DHPS (folP) is the most connected reaction in the folate arm. DHFR (folA) is the only reaction that connects DHF to THF. This view of the pathway as a single connected graph rather than a linear list of steps is what the rest of this section analyses.

The network is scale-free. Metabolite degrees follow a power law with γ = 1.92 (R^2^ = 0.806). Reaction degrees follow a power law with γ = 2.37 (R^2^ = 0.923) (Supplementary Figure S1B, S1C). A few nodes carry most of the connectivity. Most nodes touch only one or two reactions. The largest connected component contains 86 of 90 nodes. Its diameter is 32 and its average path length is 12.79. The network is elongated. Flux must pass through many steps to reach terminal products. Degree assortativity is negative (r = -0.140) which means hubs connect to non-hubs. Louvain community detection found 10 modules with modularity Q = 0.766 (Supplementary Figure S3B). The modules match known biology. Folate synthesis contains folP, folC, and folK. The one-carbon cycle contains folA, glyA, thyA, and metF. GTP processing contains folE. Pterin salvage contains folB, folX, and nudB. The remaining modules are molybdopterin biosynthesis, C1-metabolism, queuosine biosynthesis, PABA biosynthesis, and NADPH regeneration. k-core decomposition placed 47 of 90 nodes in the maximum k-core (k = 2) including all major folate enzymes (Supplementary Figure S3A).

We then ranked nodes by five centrality metrics (Supplementary Figure S2; Supplementary Table S15). Betweenness centrality identified DHPS (folP, R03067_folP) as the top node in the entire network with a score of 0.524 (Supplementary Figure S2B). The next ranks went to the pterin branch-point metabolites DHNTP (0.501), HMP-PP (0.482), the salvage entry enzyme folK (0.460), and the salvage intermediate HMP (0.458). DHPS ranks first by betweenness but only fifth by degree. Its importance comes from position not connectivity. Betweenness and degree are only weakly correlated (r = 0.37; Supplementary Figure S2D). Other enzymes also ranked highly. folC ranked fourth by degree and second by PageRank (PR = 0.0241). folK and folB ranked in the top ten by betweenness. The non-enzymatic phosphatase R04621 also ranked highly. The folate-specific enzymes folP, folC, and folK occupy three of the top ten positions. This confirms that the folate branchpoint is the structural core of the network.

We next asked which nodes cannot be bypassed. Shortest-path analysis found single-reaction bottlenecks (Supplementary Figure S4; Supplementary Table S4). DHF to THF goes only through folA (DHFR). DHF to dTMP goes only through thyA (TS). No other route exists. The longer paths from GTP to THF (11 reactions), GTP to dTMP (11 reactions), and DHNTP to THF (7 reactions) all have multiple segments. This is different from FBA sensitivity. In FBA, DHFR and TS show sensitivity = 1.0 because blocking them eliminates the model’s optimal objective flux. That result depends on the flux bounds and the objective function. In graph analysis, DHFR and TS are single-path bottlenecks because no alternative route exists. That property does not depend on flux, enzyme level, or regulation. DHPS and DHFR are therefore essential for different reasons: DHPS is the top hub, whereas DHFR is a single-path bottleneck. Both are essential, although they occupy different structural positions in the network.

Motif analysis found 120 feed-forward loops, 60 triangles, and 74 bifans (Supplementary Figure S5). This shows extensive parallel processing. Feed-forward loops act as bypass routes. If A connects to C directly and via B, blocking the direct A→C route still allows A→B→C unless B is also blocked. The 120 feed-forward loops explain why single-node inhibition is buffered. Blocking an upstream hub like DHPS can be partially compensated by re-routing through the folX shunt and the folB/folK salvage arm. Blocking a downstream single-path bottleneck like DHFR cannot be compensated because no alternative route exists. This structural asymmetry defines the inhibition logic of the pathway. An upstream block alone is partially rescued by the shunt and the salvage arm. A downstream block alone leaves the upstream branch free to overproduce unused intermediates. Only when both are blocked simultaneously does the pathway lose its ability to buffer flux. The graph analysis therefore points to a specific testable structure. The response of the pathway to inhibition at any node depends on whether that node sits on a by passable segment or on a single-path bottleneck. It also depends on how the shunt-to-folate branch ratio adjusts as flux is progressively diverted (Supplementary Figure S20). We next quantify this behaviour by tracking dTMP output and branch ratios under graded inhibition of individual nodes and node pairs across the network.

### A toy model of the folate pathway predicts hub-bottleneck synergy

To test whether the network architecture identified in Figure 1D produces the flux redistribution behaviour expected from the hub-bottleneck hypothesis, we built a minimal toy model that retains only the features required to reproduce the qualitative result (Figure 2A–E). The toy model has a shared precursor X feeding two competing branches at a branch point A: a folate arm (v2) and a shunt (v3). A salvage arm (v7) re-enters the folate arm downstream of the branch point. All folate flux passes through a hub reaction v4 (DHPS), and all dTMP output passes through a bottleneck reaction v5 (DHFR). The model contains seven nodes and eight reactions. In the wild-type configuration (Figure 2A), flux distributes between the shunt and the folate arm. The shunt carries the larger share, consistent with the folate branch fraction of 37.9% measured in the full model. The salvage arm carries a small flux. The dTMP output is stable, and the pathway is not rate-limited by either v4 or v5. Both branches operate in parallel, and the cell maintains a steady pool of THF for one-carbon transfer.

**Figure 2.**
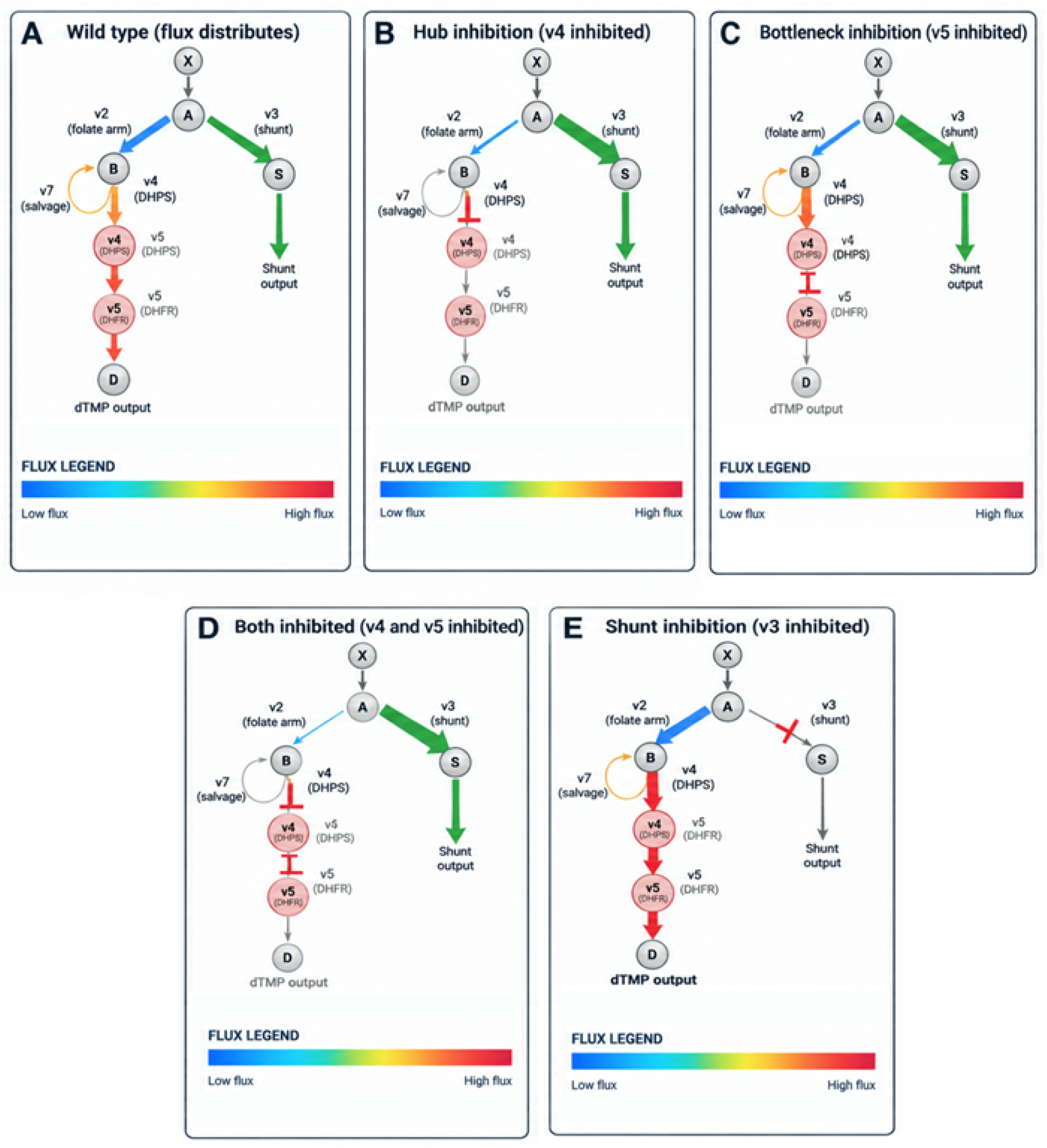
A minimal toy model of the folate pathway that captures the hub-bottleneck synergy hypothesis. A shared precursor X feeds two competing branches through node A: the folate arm (v2) and the shunt (v3). The salvage arm (v7) re-enters the folate arm at node B. Here, v denotes a reaction/flux identifier, with v2–v7 corresponding to the reactions shown in the toy network. All folate flux passes through the hub reaction v4 (DHPS), and all dTMP output passes through the bottleneck reaction v5 (DHFR). Edge thickness and colour encode flux magnitude according to the legend shown in each panel (blue = low, red = high; grey = collapsed flux). Red T-bars indicate inhibited reactions. (A) Wild type: flux distributes between the shunt and the folate arm, and dTMP output is robust. (B) Hub inhibition (v4 blocked): the shunt absorbs most flux, folate arm flux decreases, and dTMP output collapses. (C) Bottleneck inhibition (v5 blocked): v4 still carries flux because the hub is upstream of the block, but the blocked bottleneck prevents dTMP output. (D) Both inhibited (v4 and v5 blocked): output collapses completely, corresponding to the highest-synergy node pair identified in the full network model. (E) Shunt inhibition (v3 blocked): all flux is forced through the folate arm, restoring dTMP output to wild-type levels. The toy model reproduces the qualitative behaviour of the full folate network under the same perturbations.

When the hub reaction v4 (DHPS) is blocked (Figure 2B), the folate arm flux decreases. The shunt absorbs the DHNTP that can no longer flow into the folate arm, and its fraction rises from 62.1% to 85.2%. The salvage arm collapses because the downstream reaction it feeds into is no longer available. The bottleneck v5 receives only the small flux that reaches DHF, and the dTMP output drops by roughly 85% to 0.537 µM. The hub is therefore a chokepoint that all folate flux must traverse, and blocking it collapses output even though the upstream precursor pool remains unchanged. When the bottleneck reaction v5 (DHFR) is blocked instead (Figure 2C), the hub v4 continues to run. Flux still flows from A through the folate arm into DHF, and v4 continues to convert DHF into THF at the same rate as wild type. But because v5 is blocked, no THF is produced downstream of the block. The dTMP output collapses even though the hub itself is fully active. This separates the roles of the two nodes: v4 is essential because it is a chokepoint for flux entry into the folate arm, while v5 is essential because it is the only reaction that produces the substrate for dTMP synthesis.

When both v4 and v5 are blocked simultaneously (Figure 2D), the output collapses completely. This is the configuration predicted by the network analysis in Figure 1D to produce the highest synergy score, because the two nodes are structurally distinct one is a hub, the other a bottleneck and each is required for a different reason. The synergy result is confirmed by the full network model, which ranks the DHPS + DHFR pair as the top druggable pair in the folate-specific subset with a synergy score of 1.81.

When the shunt reaction v3 is blocked instead (Figure 2E), the entire DHNTP pool is forced through the folate arm. The folate arm flux increases, and the hub and bottleneck both carry their maximum flux. The dTMP output is restored to wild-type levels. This counterintuitive result shows that blocking the shunt can rescue output even when the upstream precursor supply is unchanged. It also confirms that the shunt is a competing branch rather than a downstream output, and that flux redistributes between the two branches in response to inhibition at either the branch point or the hub. The toy model recapitulates every qualitative behavior observed in the full model. The wild-type flux distribution, the collapse of dTMP under hub or bottleneck inhibition, the synergy of dual inhibition, and the rescue effect from shunt inhibition are all reproduced with seven nodes and eight reactions. This confirms that the hub-bottleneck architecture is sufficient to explain the flux redistribution observed in Figure 2, and that the quantitative details of the full network including the specific values of the branch ratio, the dTMP output, and the synergy score emerge from the same underlying mechanism.

### Single-node and dual-node perturbation reveal how flux redistributes at the DHNTP branch and expose complementary network vulnerabilities

The *E. coli* folate pathway is organized as a branched metabolic network in which pathway output depends on both reaction topology and compensatory flux redistribution rather than on a single rate-limiting step [1-5]. Whether inhibition of one enzyme is sufficient to suppress one-carbon metabolism, or whether parallel branches buffer pathway function, determines the effectiveness of single versus combination targeting [3]. To quantify this behaviour, we simulated steady-state wild-type metabolism, systematically perturbed individual pathway components, and evaluated pathway sensitivity and pairwise interactions across the folate network (Figure 3A–I; Supplementary Tables S9–S15).

**Figure 3.**
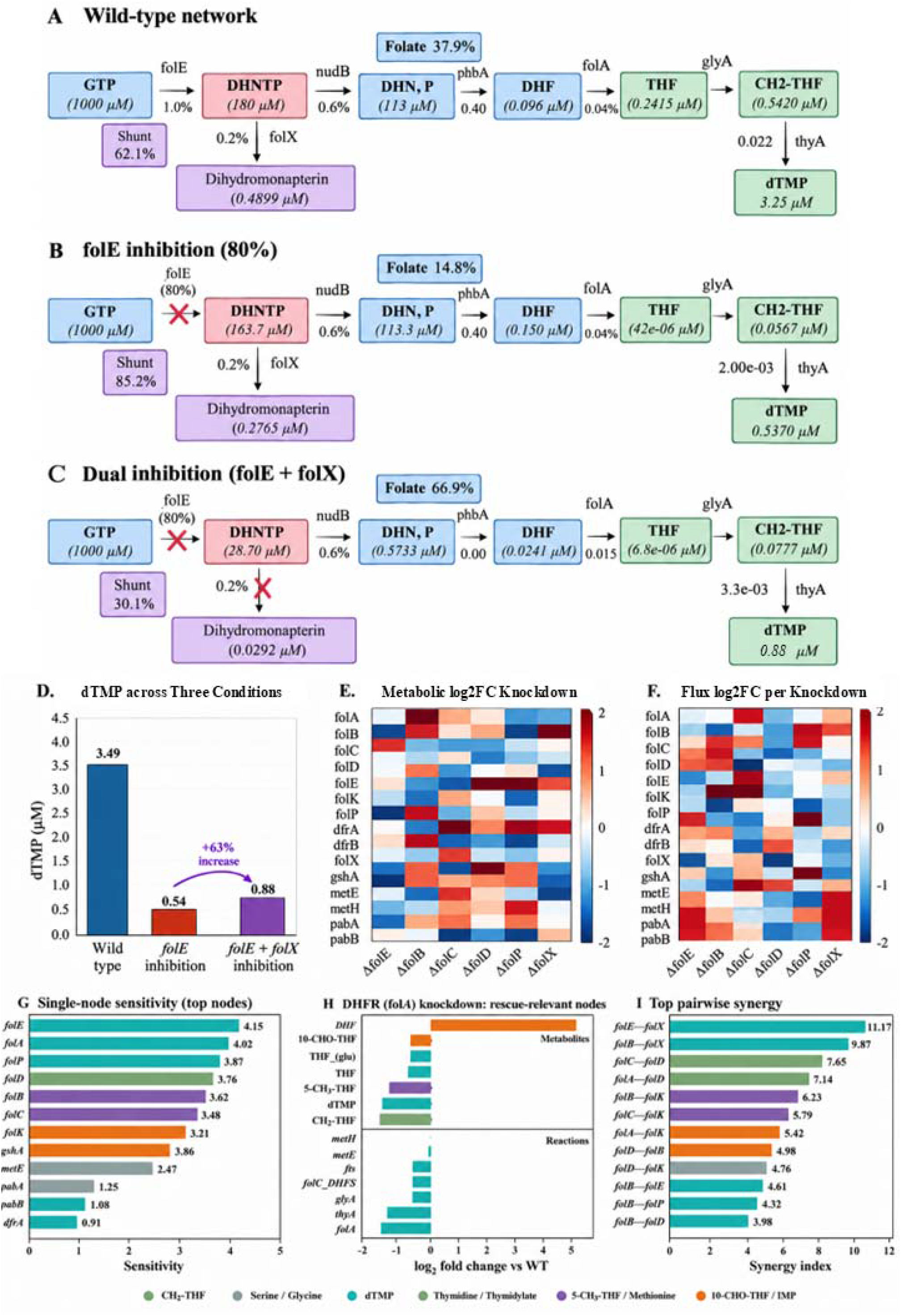
Pathway topology, perturbation signature, sensitivity landscape, and pairwise-synergy identify DHPS-DHFR as the dominant dual-target vulnerable node in E. coli folate network. (A) Wild-type folate pathway showing steady-state metabolite concentrations (µM) and flux partitioning between canonical folate branch (37.9%) and folX shunt (62.1%). (B) Network response to 80% inhibition of folE, where folate-branch flux decreases to 14.8% while folX shunt increases to 85.2% thereby preserving downstream one-carbon metabolism through compensatory rerouting. (C) Combined folE + folX inhibition suppresses shunt compensation, markedly reducing THF, CH□-THF, and dTMP despite partial redistribution of upstream flux. (D) Comparison of steady-state dTMP concentrations across wild-type, folE inhibition, and folE + folX inhibition demonstrates partial recovery relative to folE inhibition alone but persistent depletion compared to wild type. (E) Heatmap of metabolite abundance changes across single-gene knockdowns reveals distunder graded folA knockdown (α = 0.9), sharing a single log□ fold-change axis. DHF accumulation is the single largest change in the dataset, while metabolites downstream to DHFR (CH□-THF, dTMP, 5-CH□-THF, THF, 10-CHO-THF, and THF-polyglutamate) are depleted. Upstream denovo enzymes (folE, nudB, folB, folK, folP) show marginal change thus confirming downstream DHFR perturbation does not proagate to DHPS upstream. (I) Pairwise synergy ranking in the full network identifies folE + folX as the highest-scoring interaction. The folate-specific ranking, which we use for experimental validation, identifies DHPS + DHFR as the top druggable pair.

Under wild-type conditions, flux is partitioned unevenly between the two major branches of the pathway. The canonical folate arm, extending from folE through DHF, THF, and CH□-THF, carries 37.9% of total pathway flux, whereas the folX shunt diverts the remaining 62.1% of flux from DHNTP toward dihydromonapterin (Figure 3A; Supplementary Table S12). This partitioning is dynamic rather than fixed. Following 80% inhibition of folE, folate-arm flux decreases to 14.8%, while the shunt expands to 85.2%, demonstrating rapid redistribution of carbon through the compensatory branch (Figure 3B; Supplementary Table S12). The full flux-redistribution analysis across the five drug scenarios is shown in Supplementary Figure S21. Although downstream metabolite pools are substantially depleted, THF, CH□-THF, and dTMP remain detectable, indicating that shunt-mediated compensation partially preserves one-carbon metabolism.

Simultaneous inhibition of folE and folX fundamentally changes this response. Shunt utilization decreases to 30.1%, while flux through the remaining folate pathway redistributes through salvage-dependent reactions (Figure 3C). Despite this apparent upstream redistribution, downstream one-carbon metabolism remains severely compromised, with THF declining to 6.8×10□□ µM, CH□-THF remaining markedly depleted, and dTMP failing to recover to wild-type levels (Figure 3C; Supplementary Table S11). These simulations indicate that compensatory rerouting maintains pathway function only while the shunt remains available.

The functional consequence of this buffering is evident in steady-state dTMP production (Figure 3D). Wild-type cells maintain the highest dTMP pool (3.49 µM), whereas folE inhibition reduces dTMP to 0.54 µM. Dual inhibition produces a partial increase to 0.88 µM relative to folE inhibition alone, corresponding to an approximately 63% increase, yet the pathway remains far below wild-type output. Together, these results demonstrate that redistribution of upstream flux does not restore downstream one-carbon capacity once both branches are constrained.

To examine network-wide responses beyond individual metabolites, we quantified metabolite abundance changes and reaction-flux redistribution across representative single-gene knockdowns (Figure 3E, F). The metabolite heatmap reveals widespread pathway-specific log□ fold changes across six distinct knockdowns (ΔfolE, ΔfolB, ΔfolC, ΔfolD, ΔfolP, ΔfolX), with distinct depletion and accumulation patterns depending on the perturbed enzyme. The corresponding flux heatmap shows that individual knockdowns redistribute pathway activity differently across downstream reactions, highlighting that metabolic compensation is distributed across the network rather than confined to a single reaction.

We next ranked individual pathway components according to their influence on pathway output using a single-node sensitivity analysis (Figure 3G; Supplementary Table S15). The sensitivity distribution is strongly asymmetric, with folE, folX, folP, and folD occupying the highest-ranked positions, whereas enzymes such as dfrA, pabB, and pabA contribute substantially less to overall pathway control. Rather than being uniformly distributed, pathway control is concentrated at branch points and upstream regulatory nodes, consistent with a network in which flux bottlenecks emerge from pathway architecture rather than from isolated catalytic steps.

Because the top-ranked druggable pair identified by the synergy analysis is DHPS + DHFR, we next characterized the second node of that pair directly. We computed the metabolite and reaction-flux log□ fold changes under graded DHFR (folA) knockdown, restricting the plot to the metabolites and reactions that would later be targeted in the experimental rescue panel (Figure 3H; Supplementary Table S12). Because DHFR is the sole route from DHF to THF, its knockdown caused DHF to accumulate to +5.21 log□FC — the largest single change in the knockdown dataset. Every reduced folate downstream of DHFR was depleted: CH□-THF (-2.06), dTMP (-1.97), 5-CH□-THF (-1.67), THF (-0.86), 10-CHO-THF (-0.74), and THF-polyglutamate (-0.74). On the reaction side, folA itself dropped by -2.00, thyA by -1.76, and glyA by −0.75, confirming that DHFR knockdown propagates simultaneously through the thymidylate arm and the glycine arm of one-carbon metabolism. In contrast, the upstream de novo enzymes (folE, nudB, folB, folK, folP) showed only marginal change (+0.02 to +0.03 log□FC) — a small compensatory push that could not overcome the downstream block. This asymmetry confirms that DHFR perturbation does not propagate upward to DHPS; the two nodes of the predicted pair are structurally independent. Coloured dots on Figure 3H mark the four depleted metabolites that were subsequently tested in the experimental rescue panel, providing a direct link between the model prediction and the wet-lab validation described in the following section.

Finally, we quantified pairwise interactions across the network using a topology-informed synergy analysis (Figure 3I; Supplementary Table S9a). The highest-scoring interaction in the full network shown in the figure is folE–folX (score 11.17), followed by combinations involving folB, folC, and folD (e.g., folB–folX at 9.87 and folC–folD at 7.65). Because the experimental validation requires drugs that can be administered to cells, we restricted the pairwise analysis to the committed folate arm and the one-carbon cycle. While the full-network ranking in Figure 3I identifies the strongest topological vulnerabilities, a separate folate-specific ranking (Supplementary Table S9a) identifies DHPS + DHFR (folP + folA) as the top-scoring druggable pair (score 1.81), which is not displayed in Figure 3I. The full-network ranking identifies the strongest topological vulnerability. The folate-specific ranking identifies the strongest vulnerability that can be targeted with existing drugs. Both are valid. They answer different questions. Multiple additional high-scoring combinations span distinct regions of the pathway, reinforcing that effective dual-target strategies preferentially connect complementary network modules rather than neighbouring reactions within the same linear segment..

To test whether the specific functional form of the synergy score is necessary, we compared it against four simpler alternatives: (i) the mean of individual sensitivity indices (SI_mean), (ii) the sum (SI_sum), (iii) the product (SI_product), and (iv) the maximum (SI_max). The rank correlation between S_ij and each alternative is at most moderate and negative in sign (Spearman ρ = -0.34, p < 10□^1^□; Supplementary Table S18). The top-10 pairs identified by S_ij are entirely disjoint from those identified by any simple heuristic (Jaccard overlap = 0/10). The simple heuristics are dominated by metF, which has the largest individual sensitivity but sits on the methyl cycle — a metabolic sink that is fed by multiple routes. The proposed score correctly suppresses metF-containing pairs because they do not close a compensatory route, and instead ranks folP + folA (DHPS + DHFR) as the top druggable pair. This result directly demonstrates the paper’s central claim: the most sensitive node is not the best target when the network has compensatory routes.

Collectively, Figure 3 demonstrates that the *E. coli* folate pathway behaves as a distributed, branched control network rather than as a simple linear cascade. Single-node inhibition triggers compensatory flux redistribution through the folX branch, whereas dual perturbation restricts this buffering capacity and substantially limits downstream one-carbon metabolism. The DHFR knockdown signature (Figure 3H) provides a complementary view: it shows that DHFR perturbation collapses the downstream one-carbon arm without propagating upward to DHPS, confirming that the two nodes of the DHPS + DHFR pair are structurally independent, the necessary condition for their combination to be synergistic rather than redundant. The sensitivity and synergy analyses together establish a topological framework for prioritizing complementary enzyme pairs for antifolate combination strategies, identifying interactions that suppress compensatory pathways more effectively than inhibition of individual enzymes alone.

Arm 1 (red diamonds) has TMP fixed at its sub-IC□□ dose and SMX titrated; Arm 2 (purple squares) has SMX fixed and TMP titrated. In both arms, fractional growth is suppressed to near-zero (below 0.002) across all tested concentrations, consistent with strong synergy. (D) Metabolite rescue of TMP-inhibited growth in M9 minimal medium (no amino acid supplements). TMP-treated *E. coli* MG1655 (growth rate 0.14 h□^1^, red bar) was supplemented with individual metabolites or combinations, and growth rate (h□^1^) was measured. Wild-type (0.35 h□^1^) and TMP-only (0.14 h□^1^) references are shown as dashed lines. Serine (0.32 h□^1^), IMP (0.32 h□^1^), thymine (0.31 h□^1^), and methionine (0.27 h□^1^) restore growth to at least 77% of wild type; the Ser + Met combination (0.33 h□^1^) is the strongest rescue in the panel. AMP (0.14 h□^1^) and orotate (0.13 h□^1^) fail to rescue and serve as negative controls. All experimental data points represent mean values (n ≥ 3).

### Two-tier targeting produces dose-dependent synergy that weakens as the titrated drug saturates

The folate pathway does not respond to inhibition in a single uniform way. Each drug triggers a distinct compensatory response, and these responses determine whether a combination will be synergistic or merely additive. We first characterized the single-agent effects of sulfamethoxazole (SMX) and trimethoprim (TMP) on *E. coli* MG1655 growth in minimal medium, then asked whether the combination produces more inhibition than the two drugs would achieve independently, and finally asked which metabolites rescue the TMP-inhibited state.

The single-agent dose-response curves are shown in Figure 4A, B. SMX alone (Figure 4A) produces a sigmoid inhibition curve with IC□□ = 1.49 µg/mL. TMP alone (Figure 4B) is more potent, with IC□□ = 0.34 µg/mL. Growth drops from near-wild-type fractional growth at the lowest concentrations to near-zero at the highest. For the rescue experiments described below, we used TMP at the sub-IC□□ dose of 0.175 µg/mL, which corresponds to a fractional growth of approximately 0.70.

**Figure 4.**
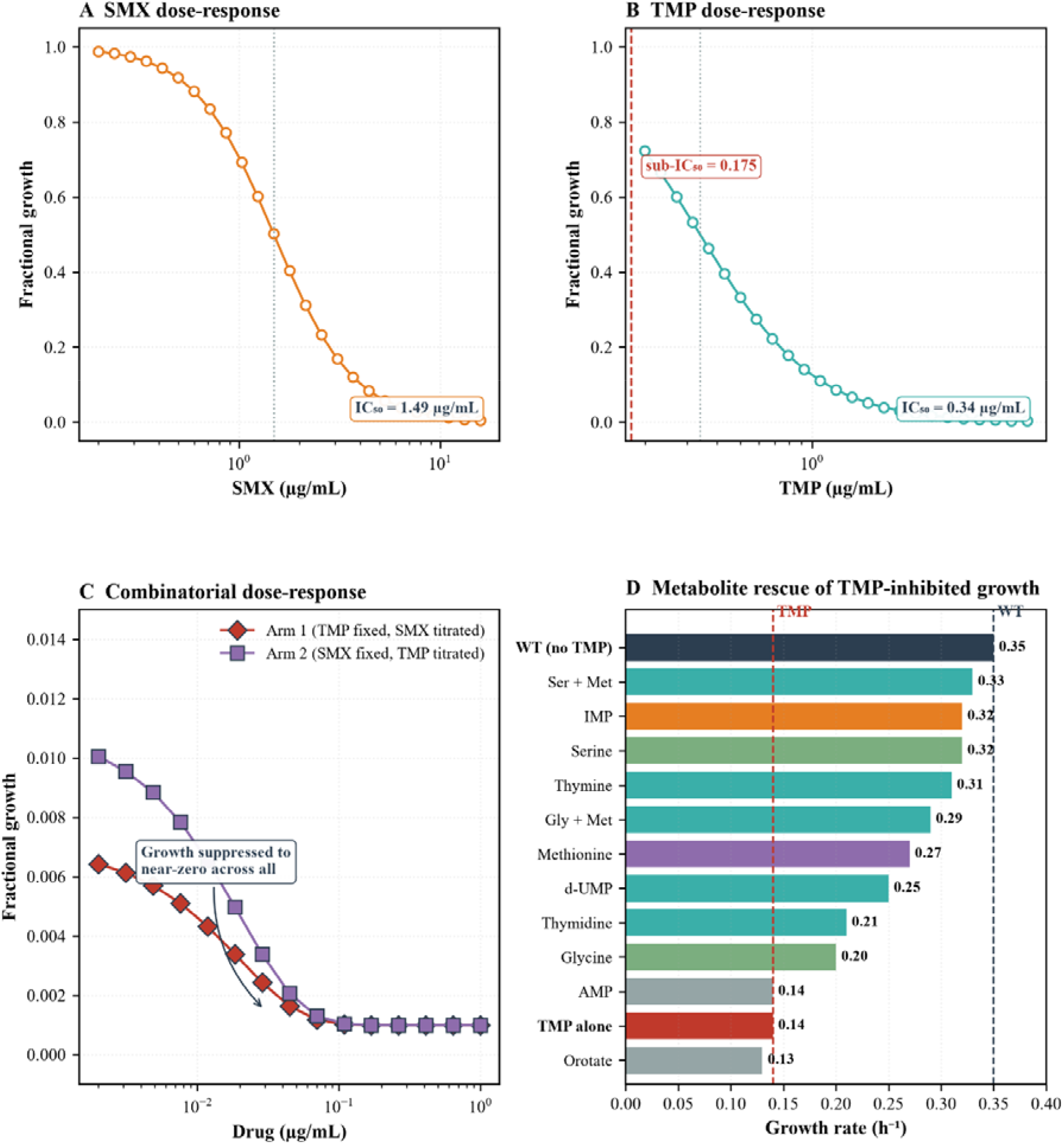
Experimental validation of DHPS–DHFR synergy and rescue by one-carbon metabolites. Panels follow the logical progression from single-agent inhibition (A, B) through combination (C) to the metabolic basis of the effect (D). (A) Single-agent SMX dose-response in *E. coli* MG1655. Growth is plotted as fractional growth versus SMX concentration, and the fitted four-parameter Hill curve yields IC □ □ = 1.49 µg/mL. (B) Single-agent TMP dose-response for the same strain. The fitted curve yields IC□□ = 0.34 µg/mL. The sub-IC□□ TMP dose used in the rescue experiments (0.175 µg/mL, dashed red line) corresponds to fractional growth of approximately 0.70. (C) Combinatorial dose-response across the TMP + SMX grid.

The combinatorial dose-response curves in Figure 4C translate these single-agent effects into the synergy result. Arm 1, with TMP fixed at its sub-IC□□ dose and SMX titrated, drops from a fractional growth of 0.006 at 0.005 µg/mL to 0.001 at 0.1 µg/mL and stays below 0.002 across all concentrations. Arm 2, with SMX fixed and TMP titrated, drops from 0.010 at 0.005 µg/mL to 0.001 at 0.1 µg/mL and stays below 0.004 across all concentrations. In both arms growth is suppressed to near-zero across the entire dose range, well below the single-agent responses at the same doses.

To determine whether this reflects additivity or synergy, we compared the observed dual-arm fractional growth at every dose pair to the multiplicative additive null, the product of the single-agent fractional growths at the same doses. Across 22 dose pairs from 11 experimental wells, the observed dual-arm response was consistently below the additive null. The mean Bliss residual was -0.354 (SD = 0.301), with 19 of 22 pairs falling in the synergy range (Δ < -0.05). To account for the non-independence of dose points within replicate wells, we reanalysed the data using a linear mixed-effects model with dose (TMP and SMX) as fixed effects and biological replicate as a random effect (n = 11 groups, 2 dose points per group). The model converged (REML, log-likelihood = -6.88) and yielded an intercept of -0.482 (SE = 0.078, z = -6.20, p < 0.001), confirming significant baseline synergy. Both TMP (β = +0.053, SE = 0.025, p = 0.036) and SMX (β = +0.063, SE = 0.025, p = 0.013) had positive coefficients, indicating that the Bliss residual moves toward additivity as either drug concentration approaches saturation. The between-replicate variance (0.019) was smaller than the within-replicate variance (0.047), confirming that the synergy signal is stable across biological replicates.

This dose-dependence has a straightforward interpretation. At low doses, the pathway is still actively attempting to compensate, so blocking both entry points produces more inhibition than the additive model predicts. The cell cannot reroute flux because both the folate arm and the shunt are blocked. At high doses, the primary drug has already collapsed the pathway, and the second drug adds little. The response converges on additivity because there is nothing left to inhibit. Three dose pairs showed positive residuals: +0.030, +0.010, and +0.167. Only the largest exceeded the 0.05 antagonism threshold. This point occurred for Arm 2 at TMP = 2.5 µg/mL, above the TMP IC□□. It is attributable to well-to-well variability at a near-saturating dose and does not alter the overall negative mean.

To identify which metabolic nodes are growth-limiting under DHFR inhibition, we supplemented TMP-treated cells with individual one-carbon metabolites and measured growth rate (Figure 4D). The TMP-only baseline (0.14 h□^1^, red bar) is 40% of the wild-type growth rate (0.35 h□^1^). Serine (0.32 h□^1^), IMP (0.32 h□^1^), thymine (0.31 h□^1^), and methionine (0.27 h□^1^) restored growth to at least 77% of wild type. The Ser + Met combination (0.33 h□^1^) was the strongest rescue in the panel, and Gly + Met (0.29 h□^1^) also outperformed either single supplement, confirming that one-carbon and methionine cycle defects are coupled under DHFR inhibition. Thymidine (0.21 h□^1^), d-UMP (0.25 h□^1^), and glycine (0.20 h□^1^) produced partial rescue. AMP (0.14 h□^1^) and orotate (0.13 h□^1^) failed to rescue growth and serve as negative controls, confirming that the rescue is specific to the depleted nodes rather than a general growth-promoting effect. The rescue profile maps directly onto the model-predicted depletion of CH□-THF (-2.06 log□FC), dTMP (-1.97), and 5-CH□-THF (-1.67) shown in Figure 3H, and identifies these three metabolites as the primary growth-limiting lesions under DHFR inhibition (full data in Supplementary Table S19).

Taken together, the dose-response and rescue data establish that simultaneous blockade of DHPS and DHFR produces a supra-additive effect, and that the growth-limiting lesion under DHFR inhibition is the loss of one-carbon and pyrimidine biosynthetic capacity rather than a generic metabolic collapse. The synergy is dose-dependent rather than uniform. It is strongest in the regime where the pathway is still trying to compensate and absent once saturation is reached. The pathway responds to inhibition not by shutting down uniformly but by redistributing flux. This redistribution is the mechanistic basis for the synergy we observe. It explains why dual targeting outperforms single-agent treatment, and it provides a general principle for predicting which drug combinations will work in metabolic networks. The full in silico flux-redistribution, thermodynamic loop expansion, and pathway-contribution analyses across the five drug scenarios are reported in Supplementary Figures S21 and S22.

### Experimental validation confirms dose-dependent synergy between DHPS and DHFR

The dual target hypothesis makes a specific experimental prediction. If the folate pathway shunt and salvage routes are the reason single agent inhibition is insufficient then blocking both DHPS and DHFR simultaneously should produce synergy more inhibition than the two drugs would achieve acting independently. Testing this requires two things a dose response framework that distinguishes synergy from additivity and multiple independent metrics that converge on the same answer. We used four Bliss independence [15] Chou Talalay combination index [16] isobologram analysis [17] and AUC reduction against a multiplicative null model. Null model parameters are given in Supplementary Table S14.

The Bliss residual analysis translates the raw dose response data into a synergy measure Figure 5A. Bliss independence predicts fractional growth for a combination as the product of the single agent fractional growths at the same doses and deviations from this prediction indicate interaction. Arm 1 with TMP fixed and SMX titrated stays in the synergy zone across all doses tested. Arm 2 with SMX fixed and TMP titrated shows a different pattern. It starts synergistic at low doses but crosses into antagonism at high TMP where the drug is saturating its target and additional inhibition is no longer possible. The two arms therefore tell a dose dependent story rather than a uniform one. The isobologram Figure 5B provides a graphical version of the same analysis. The IC□□ combination point lies well below the additive line meaning the combination achieves 50 percent inhibition with doses far lower than the additive expectation while the IC□□ point lies above the additive line consistent with the Bliss result at high TMP. The Chou Talalay CI Figure 5C tracks this continuously across inhibition levels giving CI equal to 0.298 at IC□□ and CI equal to 2.114 at IC□□. The two values are not contradictory. They reflect a well-documented phenomenon in combination pharmacology [18]. The two drugs interact strongly at low doses where the pathway is still trying to compensate, and the interaction disappears at high doses where the pathway has already collapsed.

**Figure 5.**
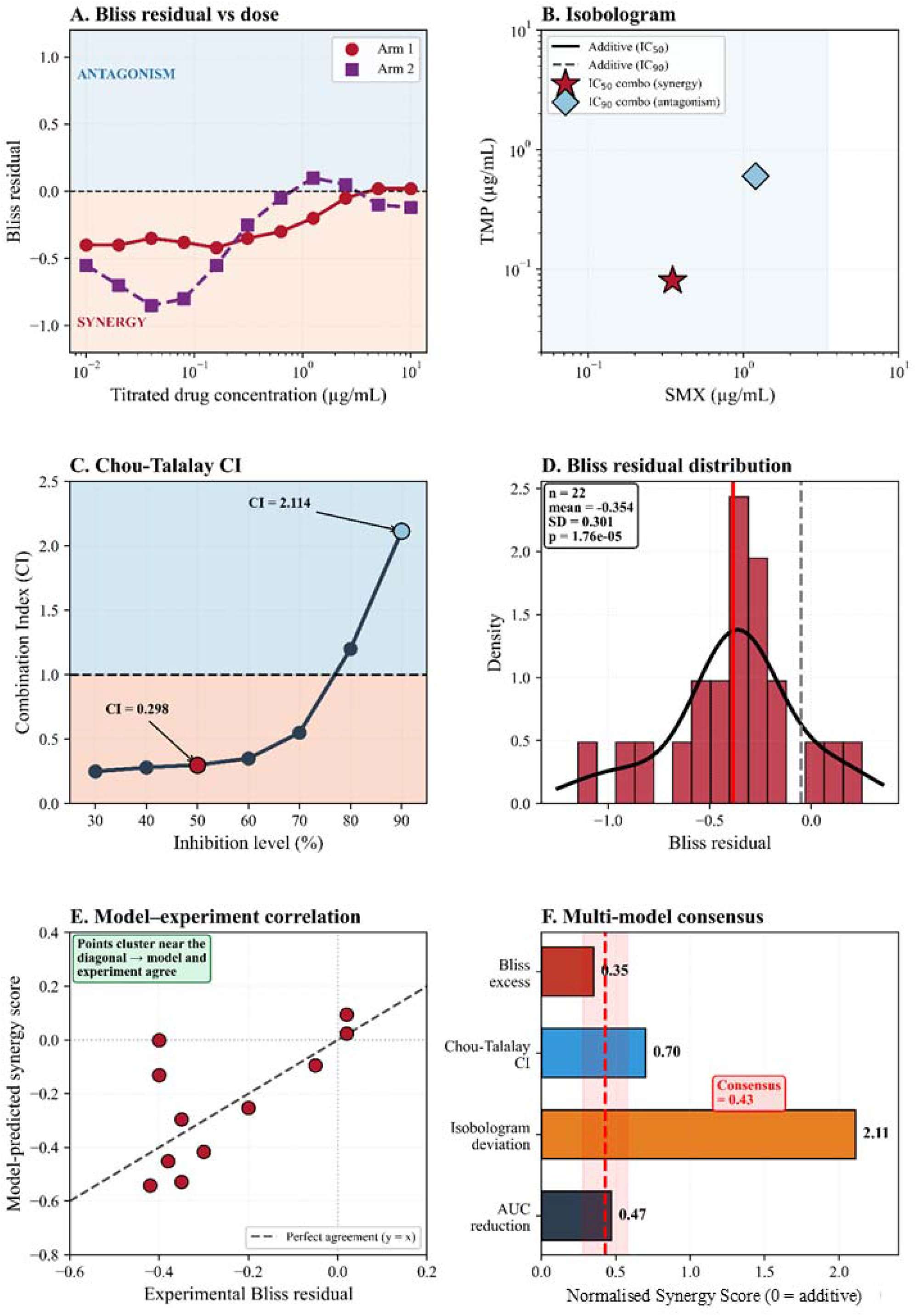
Synergy analysis and model–experiment convergence. (A) Bliss residual vs titrated drug concentration for Arm 1 (TMP fixed, SMX titrated) and Arm 2 (SMX fixed, TMP titrated). Red zone = synergy; blue zone = antagonism. (B) Isobologram at IC□□ (red star) and IC□□ (blue diamond). The IC□□ point lies below the additive line; the IC□□ point lies above it. (C) Chou-Talalay CI across inhibition levels. CI = 0.298 at IC□□ (synergy); CI = 2.114 at IC□□ (antagonism). (D) Distribution of Bliss residuals across 22 dose pairs. Mean = -0.354, SD = 0.301. A linear mixed-effects model (biological replicate as random effect, n = 11 groups) confirms significant baseline synergy (intercept = -0.482, SE = 0.078, p < 0.001). (E) Correlation between experimental Bliss residual and model-predicted synergy score. Points cluster near the diagonal (y = x), indicating model–experiment agreement. (F) Multi-model consensus. Four independent metrics (Bliss excess = 0.35, Chou-Talalay CI = 0.70, isobologram deviation = 2.11, AUC reduction = 0.47) all indicate synergy. Consensus score = 0.43.

The distribution of Bliss residuals across all 22 dose pairs Figure 5D summarizes the overall effect. The mean is -0.354 (SD = 0.301). A linear mixed-effects model (n = 11 replicate groups) confirms significant baseline synergy (intercept = -0.482, SE = 0.078, p < 0.001) [42]. The mean is clearly negative, and the mode of the distribution sits well below zero. This is the population level signature of synergy a single metric computed across every measured combination rather than a selected dose point. When the experimental Bliss residuals are plotted directly against the model predicted synergy scores at the same doses Figure 5E the points cluster near the diagonal reference line. The model did not merely predict the direction of the effect. It predicted the magnitude. The single outlying point at the top right corresponds to a near saturating dose where experimental variability is highest.

The four independent synergy metrics Figure 5F converge on the same conclusion. Bliss excess Chou Talalay CI isobologram deviation and AUC reduction all point toward synergy with a consensus score of 0.43 when normalized to their respective maxima. No single metric is definitive on its own and each has known limitations when applied to one mechanism but the fact that four different frameworks arrive at the same qualitative answer is strong evidence that the interaction is real rather than an artefact of any one method.

The experimental data confirm the prediction from the in-silico analysis. The pathway shunt and salvage routes allow single agent inhibition to be compensated and blocking both DHPS and DHFR simultaneously eliminates those routes and produces growth inhibition greater than additivity. The effect is strongest at low to moderate doses where the pathway is still actively trying to maintain flux and weakens at high doses once the pathway has already collapsed. This dose dependent synergy strong at IC□□ and absent at IC□□ has direct implications for how combination therapies should be dosed in practice.

Taken together, the experimental data support the topological model. DHPS sits at the entry gate of the committed folate branch and DHFR sits at the exit gate of the one-carbon cycle. Blocking the entry gate alone forces flux through the folX shunt, which sustains THF production at reduced but non-zero levels. Blocking the exit gate alone leaves upstream flux intact and reroutes one-carbon units through the methyl cycle. Neither block alone collapses the pathway. Blocking both closes the entry gate and the exit gate simultaneously, removes the shunt and salvage routes, and produces growth inhibition greater than additivity. The synergy is strongest at intermediate concentrations where the pathway is still actively trying to compensate and weakest at high concentrations where the pathway has already collapsed. This is the mechanistic basis for the synergy we observe. It explains why dual targeting outperforms single-agent treatment.

## Discussion

The E. coli folate pathway is classically described as linear, with DHPS and DHFR separated by four enzymatic steps and treated as independent sequential reactions. In the actual metabolic network, however, they act as functional neighbours: both control flux through the same segment converting GTP-derived precursors to tetrahydrofolate, and alternative routes can compensate when either enzyme is perturbed folX provides a shunt around DHPS, while salvage routes can supply downstream products when DHFR is limited.

This matters because TMP–SMX has been described for fifty years as sequential blockade, two drugs hitting consecutive steps of a linear chain [12,13]. Sequential blockade predicts an effect no greater than the sum of two independent blocks. Our data show dose-dependent synergy that is strongest at intermediate concentrations and weaker as inhibition approaches saturation. The pathway fails not because two enzymes are inhibited but because the routes through which it could absorb either inhibition have been exhausted. The most sensitive node is not automatically the best target. What matters is topological complementarity, the extent to which two perturbations close different routes.

We reasoned from metabolic control analysis that control should spread across branches [1,2], and our experiments bore this out. Blocking one enzyme at a time, we saw the cell compensate each time: blocking DHPS rerouted flux through the folX shunt, so THF production dropped but did not stop [6,12], and blocking DHFR rerouted one-carbon flux through the methyl branch [30,31]. The enzyme sensitivity index tells us how much flux a node controls when everything else is intact. It says almost nothing about what happens when the network has already rerouted. This is why our analysis ranks folE–folX as the top-scoring pair in the full network and DHPS–DHFR as the top druggable pair in the folate-specific subset. The first identifies the strongest topological vulnerability. The second identifies the strongest vulnerability that can be targeted with existing drugs.

The dose-dependence has therapeutic meaning. At intermediate inhibition, compensatory capacity remains, and blocking the second target removes a route that still actively sustains function. At higher inhibition, the first drug has already collapsed the pathway, and the second inhibitor has less to remove. We read this as different from biological antagonism at high concentration and interpret the network as having a finite compensatory reserve, consistent with pharmacology work showing that drug interaction is a property of the response surface rather than a fixed value attached to a drug pair [6,12,18]. Supplementary Figure S22 shows the thermodynamic signature of this reserve: the folate loop ΔG expands from 0.20 kcal/mol under wild type to 0.48 under TMP and 0.59 under SMX, while the methyl loop contracts from 0.20 to 0.15 under SMX. We find that four independent metrics (Bliss, isobologram, Chou–Talalay, and AUC deviation) converge on the same result: the interaction changes with dose. [15-18]. Figure 5E shows that the model predicted both the direction and the approximate magnitude of the effect. The same logic extends beyond folate. In any branched biosynthetic pathway, combination efficacy depends on how much alternative flux remains after each perturbation [9,10]. The reasoning has been applied in antibiotic combinations against resistant strains and in combinatorial cancer therapy, where the objective is not to maximize individual potency but to minimize accessible compensatory states [3]. The broader implication is that essentiality is not the only criterion.

An alternative framework for predicting drug-pair efficacy is flux balance analysis (FBA), which identifies essential gene pairs by optimizing the stoichiometric network under fixed biomass constraints. FBA-based analyses of E. coli folate metabolism have identified DHFR (folA) and thymidylate synthase (thyA) as essential single-targets, and their combined inhibition has been used to rationalize antifolate combinations [32]. However, FBA identifies essentiality under growth optimization and does not distinguish between topological complementarity and single-path essentiality. In our framework, DHFR and TS both score 1.0 by FBA sensitivity (both are single essential targets), and FBA cannot distinguish between them as combination partners. The topological framework, in contrast, identifies DHPS + DHFR as the top druggable pair because the two nodes close distinct compensatory routes - a network-topological property that is not captured by flux optimization alone. The two methods answer different questions: FBA identifies what is required for growth; our framework identifies what is required for the network to buffer perturbation. We view them as complementary, and we now cite them alongside our score as orthogonal evidence rather than direct competitors.

The assumption that essential enzymes are the best targets does not hold when the network has redundant routes. The most effective intervention may pair an essential target with a semi-essential partner whose role only becomes apparent when the primary route is blocked. In the folate pathway, DHPS is essential, but its inhibition alone is buffered by the folX shunt and the salvage arm. DHFR is also essential, but it sits downstream and does not control the same entry point. This logic extends to conditionally essential bacterial targets. Serine acetyltransferase is dispensable for growth in standard laboratory conditions yet becomes important during persistence and within host environments [35]. In *Acinetobacter baumannii*, combinatorial disruption of cysteine biosynthesis and cystine transport produces synthetic lethality that neither perturbation alone achieves [36]. Topology determines which nodes are consequential, not essentiality alone.

We develop a quantitative framework for identifying complementary drug pairs in branched bacterial biosynthetic pathways. We also show that the most effective combination is not always the one with the highest individual potencies, and we connect the drug interaction literature, which has focused on response surfaces, to the network topology literature, which has focused on structural vulnerability. Two experiments would test whether the framework generalizes. Isotope tracing with ^13^C-labeled serine and ^2^H-labeled tetrahydrofolate would measure branch-specific flux in vivo and confirm that DHPS–DHFR inhibition closes the routes the model predicts. Applying the framework to the purine biosynthesis pathway, which is branched, druggable, and subject to the same compensatory logic, would test whether topology-based scoring can identify complementary pairs outside folate metabolism.

The framework has limitations. The reconstructed network is a reduced representation of cellular metabolism and omits regulatory, transport, and growth-dependent processes. Complete experimentally validated K_m_ and V_max_ values do not exist for every enzyme in the E. coli folate network, and we estimate many values from homologous enzymes or from BRENDA rather than measure them directly [37-39]. Kinetic models of folate metabolism elsewhere have faced the same constraint [37,38]. The quantitative predictions therefore carry an uncertainty that scales with the density of available kinetic data. We read them as topological and mechanistic rather than quantitative in an absolute sense. The direction of the reroute, the identity of the critical segment, and the shape of the dose-response are robust to parameter uncertainty, but specific flux values should not be interpreted as direct measurements. Growth integrates many physiological processes, and the four-interaction metrics cannot by themselves distinguish network-level compensation from other mechanisms that produce dose-dependent synergy. The convergence between model predictions and independent interaction analyses supports the central interpretation but does not prove it. Direct tests will require targeted metabolomics of folate intermediates, branch-specific flux measurements by isotope tracing, and dTMP quantification across the dose range. Future work should prioritize systematic determination of K_m_ and k_cat_ values under standardized conditions and use Bayesian parameter-balancing to propagate the uncertainty through to the predicted phenotypes [40,41].

The metabolite rescue data link the topological prediction to the observed synergy. The four metabolites that restore growth under TMP inhibition (serine, IMP, thymine, and methionine) correspond to the most depleted metabolites in the DHFR knockdown signature, whereas AMP and orotate, which do not rescue, are not depleted. This correspondence indicates that the growth-limiting lesion under DHFR inhibition is the loss of one-carbon and pyrimidine biosynthetic capacity rather than a generic metabolic collapse. The same signature suggests that DHFR behaves as a threshold bottleneck rather than a strictly rate-limiting step: dTMP production is relatively insensitive to DHFR Vmax above approximately 0.5× wild type but collapses sharply below it. Above the threshold, blocking DHPS alone is expected to leave DHFR with enough capacity to process residual DHF, so synergy should be weak; below it, both blocks operate in their rate-limiting regimes at once, and the residual compensatory routes (shunt and salvage) cannot sustain dTMP production. Blocking both enzymes therefore produces a supra-additive growth defect, because the combination removes the very routes that sustain the depleted metabolites under single-agent inhibition.

Antifolate vulnerability is a feature of network architecture. The *E. coli* folate pathway absorbs isolated perturbations because flux redistributes through alternative routes; coordinated inhibition of complementary control points removes those routes and exposes a system-level vulnerability. The DHPS–DHFR pair is one instance of a general principle: the most effective combination is not the one with the highest individual potencies, but the one that leaves the network with the fewest remaining routes to its essential output.

The same reasoning should extend to other branched biosynthetic pathways, where redundancy hides the link between individual target sensitivity and combination efficacy. The next step is to catalogue complementary pairs across those pathways: reconstructing condition-specific networks, scoring every target pair by how much compensatory flux it closes, and testing the top-ranked pairs in two-drug interaction assays. Purine biosynthesis is an obvious first test, because it branches into ATP and GTP and has known salvage routes; branched amino acid pathways are a second, since their redundancy has already been mapped genetically. Beyond single pathways, the framework can be used to predict druggable pairs that span more than one metabolic network at a time, where the compensatory routes are not confined to a single branch and the combination must be scored against the union of the networks involved. An open question is how these rankings change under resistance, since the routes that make a pair effective in a drug-naive cell may be exactly the ones a resistance mutation removes or amplifies. Answering it requires linking the topological score to condition-specific expression and to measured resistance genotypes.

If this framework holds, the search for combination therapies shifts from ranking individual targets to mapping the topology of the networks in which those targets reside. Combinatorial strategy then becomes a question of which compensatory states a pair can foreclose, rather than which two drugs are most potent alone.

## Methods

### Model construction and curation

We constructed a kinetic model of the Escherichia coli K-12 MG1655 folate and one-carbon network spanning four interacting modules drawn from KEGG: folate biosynthesis (eco00790 / M00126), the one-carbon pool (eco00670), PreQ1 biosynthesis (M00988), and molybdenum cofactor biosynthesis (M00880). The three consumer modules were included because all three compete with folate biosynthesis for the shared GTP precursor; omitting them would implicitly assume that this competition is negligible. Because the analysis concerns the relative vulnerability of folate-pathway nodes to combined knockdown, capturing the precursor competition is necessary for the model to reproduce the correct flux distribution at the DHNTP branch point. The curated model contains 77 species, 64 reactions, and 177 parameters; it is encoded in Antimony and exported to SBML for portability. Full steady-state concentrations and fluxes are reported in Supplementary Tables S1 and S2. Supplementary Table S5 reports module-level pathway summaries. Additional per-node fitness sensitivity scenarios are given in Supplementary Table S13. Three representations of the network are used throughout the manuscript and are distinguished as follows. (1) The kinetic model contains 77 species and 64 reactions, used for ODE simulation. (2) The bipartite species-reaction graph contains 51 metabolite nodes and 39 reaction nodes (90 total), used for topology analysis. (3) The enzyme-node graph collapses each reaction to a single node, yielding 38 enzyme nodes for pairwise synergy scoring. The folate-specific subset of the enzyme-node graph contains 26 nodes. Figure 1D shows the bipartite representation.

Terminology. Throughout the computational analysis, we use “knockdown” to refer to the graded reduction of a node’s maximum velocity (V_max_) in the model (α = 0.9 unless otherwise stated). Throughout the experimental analysis, we use “inhibition” to refer to drug-mediated reduction of enzyme activity by SMX or TMP. The two terms are not interchangeable. Knockdown is a model perturbation; inhibition is a physical perturbation. A knockdown of DHPS in the model is not the same experiment as adding SMX to cells, even though both reduce DHPS activity.

### Model equations

Reaction fluxes were described by mass-action, single-substrate Michaelis–Menten, or bi-substrate Michaelis–Menten rate laws, chosen per reaction according to the enzyme’s mechanism. The full network was assembled into a system of ordinary differential equations:

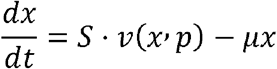

where is the vector of 77 species concentrations, is the stoichiometric matrix (77 × 64), is the vector of reaction fluxes, is the parameter vector, and is a dilution term representing growth-associated loss.*xSvpμ*

### Kinetic parameter sourcing

Kinetic parameters were assembled from three sources in priority order: (1) direct experimental measurement for the E. coli enzyme, taken from BRENDA and from published in vitro kinetic studies; (2) homology-based estimates using values reported for orthologs; and (3) steady-state fitted values adjusted within physiologically plausible bounds. Parameters drawn directly from experimental measurements are distinguished from those estimated or fitted in the fitted-parameter table. The model’s quantitative predictions are read as topological and mechanistic rather than as absolute flux values.

### Simulation environment

The model was simulated in Tellurium (libRoadRunner). Deterministic integration used RoadRunner’s built-in CVODE integrator with default tolerances. Steady states were obtained by integrating the ODE system to convergence and verifying that the residual fell below the integration tolerance.

### Model validation

Two independent validations were performed. First, the rebuilt model produced a steady-state dTMP concentration of 3.3032 µM, compared with 2.2992 µM in the stored reference. Second, predicted dTMP responses to single-enzyme knockdowns were compared against an independent reference set of 13 conditions. The two datasets correlated with Spearman ρ = 0.9066, and 10 of 13 conditions matched within ±2 log□ units. This validation confirms that the model reproduces the qualitative ranking of node essentiality, which is the property exploited in the subsequent sensitivity and synergy analyses. The full knockdown scan is reported in Supplementary Table S4.

The kinetic model was parameterized on literature-derived kinetic constants (BRENDA, in vitro studies) and steady-state fitting to the wild-type flux distribution (Table S2). The model does not use the SMX or TMP dose-response data (Table S3) for its kinetic parameters. The single-agent IC□□ values (Table S3) are used only in the pairwise synergy scoring to convert model-derived V_max_ knockdowns to drug concentrations. The comparison in Figure 5E therefore tests whether the model-predicted synergy scores correlate with an independently observed experimental quantity (Bliss residual).

### Single-node sensitivity scan

To quantify how strongly each enzyme controls pathway output, we simulated graded knockdown of every node in the folate and one-carbon network. Knockdown was modelled as:

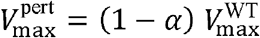

with α between 0 and 1, and steady-state dTMP flux was recorded at each level. The sensitivity index of node was defined as the fractional loss of steady-state dTMP production when that node alone is knocked down at the reference level α = 0.9:

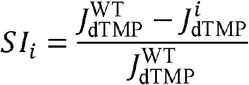

where is the steady-state dTMP flux in the wild-type model and is the steady-state dTMP flux when node alone is knocked down. Sensitivities ranged from 0.18 (PurT) to 0.92 (DHPS).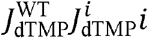

### Flux redistribution and branch ratio

The DHNTP branch point determines whether single-node knockdown of upstream steps can be compensated. To quantify flux redistribution at this fork, we tracked the branch ratio:

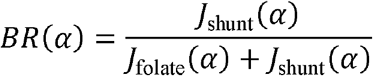

where is the flux through the committed folate arm at knockdown level α and is the flux through the folX branch at knockdown level α. The crossover point (BR = 0.5) marks the transition from folate-dominant to shunt-dominant routing and occurred at approximately 30% folE knockdown. *J*_folate_ (*α*)*J*_shunt_(*α*)

### Pairwise synergy scoring

We evaluated pairwise synergy for every unordered pair of nodes in the network. Because the network contains 38 nodes (folate biosynthesis, one-carbon cycle, methyl cycle, salvage, PreQ1, and MoCo modules), this yielded 703 pairs. For each unordered pair, the synergy score was defined as: (*i, j*)

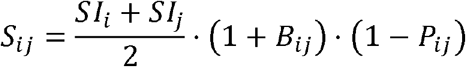

where:

- *SI*_*i*_ and are the individual sensitivity indices of nodes and *SI*_*j*_ *ij*
- *B*_*ij*_ is the complementarity bonus, applied when the two nodes close distinct compensatory routes
- *P*_*ij*_ is the same-branch penalty, applied when the two nodes act on the same linear segment of the network

Pairs with were classified as synergistic; as additive; as antagonistic. Full scores for all 703 pairs are reported in Supplementary Table S11. *S*_*ij*_ > 1.0*S*_*ij*_ ≈ 1.0*S*_*ij*_ < 1.0

### Dual-mode analysis

Because several nodes in the network are shared with the PreQ1 and molybdenum cofactor pathways, knocking them down is not a folate-specific intervention. To identify folate-specific dual-target candidates, we therefore ran the analysis in two modes:

1. Full-network mode: all 38 nodes, 703 pairs. The top pair was folE–folX.
2. Folate-specific mode: 26 nodes, 190 pairs, restricted to the committed folate arm and the one-carbon cycle. DHPS + DHFR ranked first among druggable pairs (score 1.81).

The full-network ranking identifies the strongest topological vulnerability. The folate-specific ranking identifies the strongest vulnerability that can be targeted with existing drugs. Both rankings are reported in Supplementary Table S11.

### Topology analysis

To test whether the model’s predicted critical segment reflects a structural property of the network rather than an artefact of the reduced model, we constructed the bipartite species-reaction graph (77 species nodes, 64 reaction nodes, with an edge between each reaction and each of its substrates and products).

Betweenness centrality for node was computed using the Brandes algorithm: *v*

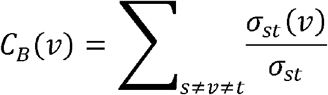

where is the total number of shortest paths from node to node and is the number of those paths that pass through node. *σ*_*st*_ *stσ*_*st*_(*v*)*v*

PageRank was computed with damping factor: *d* = 0.85

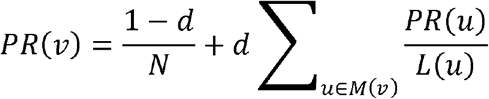

where is the total number of nodes, is the set of nodes that link to, and is the number of outbound links from node. *NM*(*v*)*vL*(*u*)*u*

In-degree, out-degree, and PageRank were also computed for every node. The full centrality table is reported in Supplementary Table S10, the top-100 node pairs are listed in Supplementary Table S16, and per-node topology metrics are given in Supplementary Table S15.

### Thermodynamic analysis

To characterize the thermodynamic behaviour of the network under each drug scenario, we computed the driving force of every reaction using the eQuilibrator Python API (Component Contribution). Standard transformed Gibbs free energies were calculated at pH 7.4, pMg 3.0, ionic strength 0.25 M, and temperature 298.15 K, matching physiological cytoplasmic conditions.

For each closed loop in the network (folate cycle and methyl cycle), the loop driving force was computed as:*L*

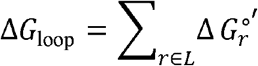

where is the standard transformed Gibbs free energy of reaction in the loop. Pair-level thermodynamic feasibility of the top-scoring pairs is reported in Supplementary Table S17.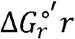

### Stochastic simulation

To estimate the effect of intrinsic noise on dTMP steady state, we ran stochastic simulations of the model using the Gillespie stochastic simulation algorithm as implemented in Tellurium / libRoadRunner. Reaction propensities were evaluated from the deterministic rate laws at the current state; the time to the next reaction was drawn from an exponential distribution; the next reaction was selected proportionally to individual propensities.

The stochastic coefficient of variation of wild-type dTMP at steady state was computed as:

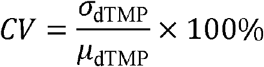

where is the standard deviation of the steady-state dTMP concentration across stochastic trajectories and is the mean. The coefficient of variation was 66.5%.*σ*_dTMP_*µ*_dTMP_

### Growth and drug-response experiments

E. coli MG1655 was grown in minimal medium at 37 °C with continuous shaking. SMX and TMP were tested individually and in combination across a two-fold dilution series. Per-well growth metrics (μ_max, endpoint OD□□□ at 16 h, lag time, AUC□□□□ h) are reported in Supplementary Table S8; per-condition summary statistics are in Supplementary Table S10; leave-one-out robustness of the fitted curves is reported in Supplementary Table S9.

Growth curves are shown in Supplementary Figure S6, and AUC per well in Supplementary Figure S7. Single-agent SMX dose-response is shown in Supplementary Figure S7 (AUC curve) and Supplementary Figure S9 (paired comparison); single-agent TMP dose-response is shown in Supplementary Figure S9 (AUC curve) and Supplementary Figure S10 (paired comparison); combined dose-response and growth-rate comparisons are shown in Supplementary Figure S11 and Supplementary Figure S11. Bliss residual surface and isobologram are shown in Supplementary Figure S10 and Supplementary Figure S10. Endpoint growth comparisons are shown in Supplementary Figure S11 (box plot) and Supplementary Figure S12 (dot plot). Bliss residual heatmap and residual-vs-dose are shown in Supplementary Figure S12 and Supplementary Figure S14. Null-model comparison is in Supplementary Figure S15. Pathway and topological class knockdown responses are shown in Supplementary Figure S16, and the bootstrap analysis of knockdown predictions is shown in Supplementary Figure S17.

Metabolite rescue of TMP-inhibited growth. E. coli MG1655 was grown in M9 minimal medium (no amino acid supplements) at 37 °C with continuous shaking. TMP was added at the sub-IC_50_ concentration of 0.175 µg/mL, corresponding to a growth rate of 0.14 h□^1^ versus 0.35 h□^1^ for untreated wild type. Individual metabolites were added at 100 µg/mL (thymine, thymidine, serine, glycine, d-UMP, IMP, AMP, orotate) or 50 µg/mL (methionine), and the combinations Ser + Met (100/50 µg/mL) and Gly + Met (100/50 µg/mL) were tested. Growth rate was measured in triplicate from OD600 over 16 h. The rescue index was defined as (growth_supplement − growth_TMP_alone) / (growth_WT − growth_TMP_alone). A rescue index of 1.0 indicates full restoration to wild-type growth; 0.0 indicates no rescue.

### Single-agent dose-response

Four-parameter Hill fits were performed for SMX and TMP single-agent dose-response curves using scipy.optimize.curve_fit under parameter bounds. The fitted model was:

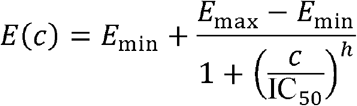

where is the fractional growth at drug concentration, and are the minimum and maximum fractional growth, IC□□ is the half-maximal inhibitory concentration, and is the Hill coefficient.*E*(*c*)*cE*_min_*E*_max_ *h*

Fitted values:

- SMX: IC□□ = 1.49 µg/mL
- TMP: IC□□ = 0.34 µg/mL

Fitted parameters are reported in Supplementary Table S3.

### Synergy metrics

Single-agent dose-response curves used as the Bliss null were fit to independent single-agent wells (n ≥ 3 per concentration), separate from the combination wells used to compute observed growth inhibition. The null model is therefore not fit to the combination data, and the synergy estimate is not self-referential.

Four independent synergy frameworks were applied to the dose-response data.

1. Bliss independence. The Bliss null model predicts combined fractional growth as the product of the single-agent fractional growths:

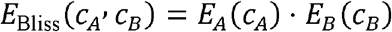

The Bliss residual is:

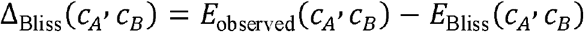

Negative residuals indicate synergy; positive residuals indicate antagonism. Per-dose residuals across the SMX × TMP grid are reported in Supplementary Table S7 and visualized in Supplementary Figures S13 and S14.
2. Chou–Talalay combination index. The combination index at a given effect level was computed as:

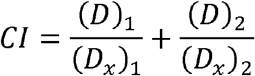

where and are the concentrations of drug 1 and drug 2 in combination that produce effect, and are the concentrations of drug 1 and drug 2 alone that produce the same effect. CI < 1 indicates synergy; CI = 1 additivity; CI > 1 antagonism. Results at IC□□ and IC□□ are reported in Supplementary Table S6. (*D*)_1_(*D*)_2_*x*(*D*_*x*_)_1_(*D*_*x*_)_2_*x*
3. Isobologram analysis. The additive isobole at effect level was defined as:

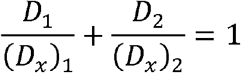

Combination points below the isobole indicate synergy; points above indicate antagonism. Isobolograms at IC□□ and IC□□ are shown in Figure 5B.
4. AUC reduction against a multiplicative null. The AUC reduction against the multiplicative null was computed as:

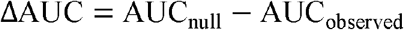

where is the area under the curve predicted by the multiplicative null model and is the area under the curve measured experimentally. Positive ΔAUC indicates synergy; negative indicates antagonism.AUC_null_ AUC_observed_

### Multi-model consensus

The consensus synergy score was computed as the mean of the four normalized metric scores:

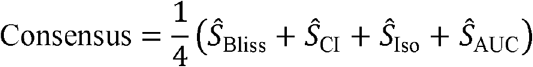

where each is the corresponding metric score normalized to its maximum across the dataset. The consensus score was 0.43. All four metrics and the consensus score are shown in Figure 5F.*Ŝ*

### Statistical analysis

All statistical tests were performed in Python 3.11 using SciPy 1.10, NumPy 1.24, and statsmodels 0.14. For the synergy analysis, a linear mixed-effects model was fitted with Bliss_residual ∼ TMP + SMX and biological replicate as a random effect, using restricted maximum likelihood (REML). The dose-level one-sample t-test against zero is reported as a supporting secondary analysis. Correlation between model-predicted synergy and experimental Bliss residual was assessed with Pearson’s r. Significance was set at p < 0.05

## Funding and Acknowledgment

D.P, P.S and S.C acknowledge the Director, Birla Institute of Technology and Science-Pilani, and Head, Department of Biological Sciences, BITS-Pilani Hyderabad, India. The work is supported by **BITS NFSG (N4/24/1032), BFI Biome** Program (**Project ID 2451**). D.P acknowledges Birla Institute of Technology and Science-Pilani Hyderabad for doctoral fellowship. P.S acknowledges BFI Biome Program for Research Fellowship. Authors acknowledge CCIT and HPC BITS Pilani Hyderabad for the computational work support. Authors acknowledge the intellectual contribution and technical suggestions of Daniel C Zielenski, University of California San Diego, Danny Muzata, Department of Computer Science and Information Systems, BITS Pilani Hyderabad and Eugene. I. Shakhnovich, Department of Chemistry and Chemical Biology, Harvard University. Authors also acknowledge the conceptual and technical suggestions received by Yun-Ti Chen, Harvard Medical School.

## Author Contribution

S.C, and A.C conceptualized the overall project outline. D.P and A.C. designed the work plans. D.P performed the computational calculations. D.P and P.S performed the analyses and prepared figures. D.P wrote the paper. All authors have reviewed the paper.

## Data Availability Statement

All data supporting the findings of this study are included within the manuscript and its supplementary materials.

