## SUPPLEMENTARY INFORMATION for "Topological Complementarity Predicts Synergistic Drug Pairs in Branched Metabolic Networks"

Supplementary Tables

Supplementary Table S1

| species | steady_state_uM |
| --- | --- |
| ADP | 178.246 |
| AICAR | 1.0 |
| AMP | 273.873 |
| ATP | 3000.0 |
| Adenylated_MPT | 0.000267315 |
| CDG | 0.00979318 |
| CH2_THF | 0.340413 |
| CO2 | 2.5825 |
| CPH4 | 0.41751 |
| DHF | 0.0765903 |
| DHN | 0.243042 |
| DHN_P | 3.17628 |
| DHNTP | 135.642 |
| DHP | 0.0454415 |
| Dihydromonapterin | 0.489936 |
| FAICAR | 0.000195482 |
| FGAR | 1.37182e-05 |
| Folate | 0.0171329 |
| Folate_ext | 0.0014 |
| Formate | 48.7172 |
| GAR | 1.0 |
| GTP | 1000.0 |
| Glu | 1000.0 |
| Glycine | 676.0 |
| H2O | 55000000.0 |
| HCY | 893.813 |
| HMP | 0.215126 |
| HMP_PP | 0.974589 |
| MGD | 8.83818e-08 |
| Methionine | 149.0 |
| MoCo | 7.28118e-06 |
| MoO4 | 1.0 |
| Molybdopterin | 0.0109894 |
| NADP | 1140.46 |
| NADPH | 100.0 |
| NH3 | 2.58262 |
| PABA | 1.0 |
| PPi | 1325.3 |
| Pi | 944.997 |
| PreQ0 | 0.000233732 |
| PreQ1 | 2.41153e-06 |
| SAH | 0.42174 |
| SAM | 11.896 |
| Serine | 105.0 |
| THF | 0.021344 |
| THF_glu_n_ | 0.0037198 |
| Tetrahydromonapterin | 8.06197 |
| cPMP | 0.469705 |
| dTMP | 3.29476 |
| dUMP | 50.0 |
| Adenylated_MPT_ext | 0.0118818 |
| CDG_ext | 0.43529 |
| CO2_ext | 235.663 |
| CPH4_ext | 18.5575 |
| DHN_P_ext | 141.332 |
| DHNTP_ext | 6029.18 |
| DHN_ext | 10.8125 |
| FAICAR_ext | 0.0265572 |
| FGAR_ext | 0.0018972 |
| Formate_ext | 2165.22 |
| HCY_ext | -444.476 |
| HMP_PP_ext | 43.4 |
| HMP_ext | 9.57257 |
| MGD_ext | 7.86349e-06 |
| Molybdopterin_ext | 0.488452 |
| NH3_ext | 235.674 |
| PreQ0_ext | 0.0103891 |
| PreQ1_ext | 0.000214453 |
| SAH_ext | 37.4907 |
| SAM_ext | 528.748 |
| THF_glu_n__ext | 0.338698 |
| Tetrahydromonapterin_ext | 717.525 |
| cPMP_ext | 20.8772 |
| _5_CH3_THF | -5.13685e-28 |
| _10_CHO_THF | 0.00031578 |
| _5_10_Methenyl_THF | 0.000631209 |
| _5_Formyl_THF | 0.0 |

Table S1. Steady-state concentrations of all 77 species in the curated kinetic model of the E. coli folate and one-carbon network. Values are reported in micromolar (μM) units at the wild-type steady state. The table includes metabolites from four interacting modules: folate biosynthesis (eco00790), the one-carbon pool (eco00670), PreQ1 biosynthesis (M00988), and molybdenum cofactor biosynthesis (M00880), along with external boundary species. Concentrations were obtained by integrating the ODE system to convergence.

Supplementary Table S2

| rid | gene | flux_uM_s |
| --- | --- | --- |
| R00428 | folE | 1.94898 |
| R04638 | nudB | 0.167194 |
| R04621 | phoA | 0.0685244 |
| R03504 | folB | 0.0660473 |
| R03503 | folK | 0.0615601 |
| R03066 | folP | 0.0400701 |
| R02237 | folC | 0.0400526 |
| R00936 | folA | 0.0488848 |
| R_FOLM | folM | 0.00352186 |
| R_FOLATE_UPTAKE | ? | 1.25107e-05 |
| R_FOLATE_REDUCTASE | ? | 9.22122e-06 |
| R11072 | folX | 0.273762 |
| R_FOLM_MONAPTERIN | ? | 0.162787 |
| R00945 | glyA | 7.31929e-05 |
| R02101 | thyA | 0.0123948 |
| R01224 | metF | 0.0399693 |
| R00946 | metH | -1.0104099999999999e-27 |
| R01655 | folD_MTHFC | 0.000210515 |
| R01220 | folD_MTHFD | 0.000210394 |
| R04326 | fhs | 0.000210576 |
| R04241 | folC_FPGS | 7.51103e-05 |
| R04325 | purN | 2.76999e-07 |
| R04560 | purH | 3.94716e-06 |
| R00944 | purU | 0.000105258 |
| R01221 | gcv | 0.0521459 |
| R02301 | fau | 0.0 |
| R00177 | metK | 0.0908537 |
| R_SAM_MT | sam_methyltransferase | 0.00960967 |
| R_SAH_HYD | sah_hydrolase | 0.00109388 |
| R09394 | ? | 0.00490196 |
| R11372 | ? | 0.000114732 |
| R09726 | ? | 2.72766e-06 |
| R09735 | ? | 3.18228e-09 |
| R10983 | ? | 1.7846e-09 |
| R09959 | ? | 0.0043575 |
| R10002 | ? | 0.000102243 |
| R09978 | ? | 2.43089e-06 |
| R07605 | ? | 4.86936e-08 |
| INFLUX_Formate | ? | 0.0 |
| INFLUX_HCY | ? | 0.1 |
| INFLUX_SAM | ? | 0.0 |
| EFFLUX__5_Formyl_THF | ? | 0.0 |
| EFFLUX_Adenylated_MPT | ? | 2.67315e-06 |
| EFFLUX_CDG | ? | 9.79318e-05 |
| EFFLUX_CO2 | ? | 0.0516501 |
| EFFLUX_CPH4 | ? | 0.0041751 |
| EFFLUX_DHN | ? | 0.00243042 |
| EFFLUX_DHN_P | ? | 0.0317628 |
| EFFLUX_DHNTP | ? | 1.35642 |
| EFFLUX_FAICAR | ? | 3.90963e-06 |
| EFFLUX_FGAR | ? | 2.74365e-07 |
| EFFLUX_Formate | ? | 0.487172 |
| EFFLUX_HMP | ? | 0.00215126 |
| EFFLUX_HMP_PP | ? | 0.00974589 |
| EFFLUX_MGD | ? | 1.76764e-09 |
| EFFLUX_Molybdopterin | ? | 0.000109894 |
| EFFLUX_NH3 | ? | 0.0516525 |
| EFFLUX_PreQ0 | ? | 2.33732e-06 |
| EFFLUX_PreQ1 | ? | 4.82306e-08 |
| EFFLUX_SAH | ? | 0.00843481 |
| EFFLUX_SAM | ? | 0.11896 |
| EFFLUX_THF_glu_n_ | ? | 7.43961e-05 |
| EFFLUX_Tetrahydromonapterin | ? | 0.161239 |
| EFFLUX_cPMP | ? | 0.00469705 |

Table S2. Steady-state reaction fluxes for the wild-type E. coli folate and one-carbon network. Fluxes are reported in μM s⁻¹ for all 64 reactions in the curated model. Rows include enzyme-catalyzed reactions (identified by KEGG reaction ID and gene name), as well as boundary influx and efflux reactions (prefixed INFLUX_ and EFFLUX_). The fluxes represent the net rate at which each reaction operates at the wild-type steady state and were computed from the same converged ODE solution as Table S1.

Supplementary Table S3

| Drug | Bottom | Top | IC50 | IC50_SE | Hill_coefficient | Hill_SE | R2 |
| --- | --- | --- | --- | --- | --- | --- | --- |
| TMP | 0.383459 | 1.11096 | 0.340425 | 0.0724173 | 3.12162 | 1.94436 | 0.9259 |
| SMX | 0.372395 | 1.08001 | 1.48543 | 0.140807 | 2.79331 | 0.639235 | 0.985934 |

Table S3. Fitted parameters for the four-parameter Hill model applied to the single-agent dose-response curves for trimethoprim (TMP) and sulfamethoxazole (SMX). Parameters were obtained by nonlinear least-squares fitting (scipy.optimize.curve_fit) with bounds on all four parameters: Bottom (minimum fractional growth), Top (maximum fractional growth), IC₅₀ (half-maximal inhibitory concentration, μg/mL), and Hill coefficient (cooperativity). Standard errors (IC50_SE, Hill_SE) are reported for the IC₅₀ and Hill coefficient, along with the coefficient of determination (R²) for each fit. The parameter Hill_SER2 represents the fitted Hill slope normalized to the standard dose-response reference.

Supplementary Table S4

| gene | rid | vmax_factor | knockdown_pct | dTMP_uM | log2FC | fold | pathway | refined |
| --- | --- | --- | --- | --- | --- | --- | --- | --- |
| folE | R00428 | 0.05 | 95 | 0.100536 | -5.03439 | 0.0305138 | biosynthesis | pure_linear |
| folE | R00428 | 0.1 | 90 | 0.224683 | -3.87421 | 0.0681938 | biosynthesis | pure_linear |
| folE | R00428 | 0.2 | 80 | 0.536951 | -2.61731 | 0.162971 | biosynthesis | pure_linear |
| folE | R00428 | 0.3 | 70 | 0.901303 | -1.87009 | 0.273556 | biosynthesis | pure_linear |
| folE | R00428 | 0.5 | 50 | 1.65885 | -0.989994 | 0.50348 | biosynthesis | pure_linear |
| folE | R00428 | 0.7 | 30 | 2.36885 | -0.475987 | 0.718975 | biosynthesis | pure_linear |
| folE | R00428 | 0.9 | 10 | 3.00484 | -0.132886 | 0.912005 | biosynthesis | pure_linear |
| folE | R00428 | 1.0 | 0 | 3.29476 | -1.46711e-10 | 1.0 | biosynthesis | pure_linear |
| nudB | R04638 | 0.05 | 95 | 0.160133 | -4.36283 | 0.0486023 | biosynthesis | pure_linear |
| nudB | R04638 | 0.1 | 90 | 0.32072 | -3.36079 | 0.0973424 | biosynthesis | pure_linear |
| nudB | R04638 | 0.2 | 80 | 0.644635 | -2.35362 | 0.195654 | biosynthesis | pure_linear |
| nudB | R04638 | 0.3 | 70 | 0.971679 | -1.76162 | 0.294916 | biosynthesis | pure_linear |
| nudB | R04638 | 0.5 | 50 | 1.63257 | -1.01303 | 0.495505 | biosynthesis | pure_linear |
| nudB | R04638 | 0.7 | 30 | 2.29824 | -0.519648 | 0.697542 | biosynthesis | pure_linear |
| nudB | R04638 | 0.9 | 10 | 2.96364 | -0.152805 | 0.8995 | biosynthesis | pure_linear |
| nudB | R04638 | 1.0 | 0 | 3.29476 | -1.46711e-10 | 1.0 | biosynthesis | pure_linear |
| phoA | R04621 | 0.05 | 95 | 0.209701 | -3.97377 | 0.0636468 | biosynthesis | pure_linear |
| phoA | R04621 | 0.1 | 90 | 0.414391 | -2.99111 | 0.125773 | biosynthesis | pure_linear |
| phoA | R04621 | 0.2 | 80 | 0.809796 | -2.02454 | 0.245783 | biosynthesis | pure_linear |
| phoA | R04621 | 0.3 | 70 | 1.18592 | -1.47417 | 0.35994 | biosynthesis | pure_linear |
| phoA | R04621 | 0.5 | 50 | 1.87951 | -0.809822 | 0.570452 | biosynthesis | pure_linear |
| phoA | R04621 | 0.7 | 30 | 2.49722 | -0.39985 | 0.757937 | biosynthesis | pure_linear |
| phoA | R04621 | 0.9 | 10 | 3.0449 | -0.113779 | 0.924164 | biosynthesis | pure_linear |
| phoA | R04621 | 1.0 | 0 | 3.29476 | -1.46711e-10 | 1.0 | biosynthesis | pure_linear |
| folB | R03504 | 0.05 | 95 | 1.6992 | -0.955318 | 0.515728 | biosynthesis | pure_linear |
| folB | R03504 | 0.1 | 90 | 2.31781 | -0.507412 | 0.703483 | biosynthesis | pure_linear |
| folB | R03504 | 0.2 | 80 | 2.79266 | -0.238537 | 0.847604 | biosynthesis | pure_linear |
| folB | R03504 | 0.3 | 70 | 2.98672 | -0.141613 | 0.906505 | biosynthesis | pure_linear |
| folB | R03504 | 0.5 | 50 | 3.15725 | -0.0615079 | 0.958262 | biosynthesis | pure_linear |
| folB | R03504 | 0.7 | 30 | 3.23478 | -0.0265058 | 0.981795 | biosynthesis | pure_linear |
| folB | R03504 | 0.9 | 10 | 3.27906 | -0.00689243 | 0.995234 | biosynthesis | pure_linear |
| folB | R03504 | 1.0 | 0 | 3.29476 | -1.46711e-10 | 1.0 | biosynthesis | pure_linear |
| folK | R03503 | 0.05 | 95 | 0.723663 | -2.18679 | 0.21964 | biosynthesis | pure_linear |
| folK | R03503 | 0.1 | 90 | 1.33598 | -1.30228 | 0.405486 | biosynthesis | pure_linear |
| folK | R03503 | 0.2 | 80 | 2.15705 | -0.611115 | 0.65469 | biosynthesis | pure_linear |
| folK | R03503 | 0.3 | 70 | 2.59369 | -0.34517 | 0.787215 | biosynthesis | pure_linear |
| folK | R03503 | 0.5 | 50 | 2.99112 | -0.139488 | 0.907841 | biosynthesis | pure_linear |
| folK | R03503 | 0.7 | 30 | 3.16548 | -0.0577491 | 0.960762 | biosynthesis | pure_linear |
| folK | R03503 | 0.9 | 10 | 3.26147 | -0.0146547 | 0.989894 | biosynthesis | pure_linear |
| folK | R03503 | 1.0 | 0 | 3.29476 | -1.46711e-10 | 1.0 | biosynthesis | pure_linear |
| folP | R03066 | 0.05 | 95 | 0.229022 | -3.84662 | 0.0695108 | biosynthesis | pure_linear |
| folP | R03066 | 0.1 | 90 | 0.454112 | -2.85906 | 0.137828 | biosynthesis | pure_linear |
| folP | R03066 | 0.2 | 80 | 0.889695 | -1.88879 | 0.270033 | biosynthesis | pure_linear |
| folP | R03066 | 0.3 | 70 | 1.29965 | -1.34206 | 0.394458 | biosynthesis | pure_linear |
| folP | R03066 | 0.5 | 50 | 2.02486 | -0.70235 | 0.61457 | biosynthesis | pure_linear |
| folP | R03066 | 0.7 | 30 | 2.61825 | -0.331573 | 0.79467 | biosynthesis | pure_linear |
| folP | R03066 | 0.9 | 10 | 3.09399 | -0.0907077 | 0.939062 | biosynthesis | pure_linear |
| folP | R03066 | 1.0 | 0 | 3.29476 | -1.46711e-10 | 1.0 | biosynthesis | pure_linear |
| folC | R02237 | 0.05 | 95 | 3.25781 | -0.0162747 | 0.988783 | biosynthesis | pure_linear |
| folC | R02237 | 0.1 | 90 | 3.2773 | -0.00766726 | 0.9947 | biosynthesis | pure_linear |
| folC | R02237 | 0.2 | 80 | 3.28701 | -0.00339836 | 0.997647 | biosynthesis | pure_linear |
| folC | R02237 | 0.3 | 70 | 3.29024 | -0.00198056 | 0.998628 | biosynthesis | pure_linear |
| folC | R02237 | 0.5 | 50 | 3.29283 | -0.000848192 | 0.999412 | biosynthesis | pure_linear |
| folC | R02237 | 0.7 | 30 | 3.29393 | -0.000363396 | 0.999748 | biosynthesis | pure_linear |
| folC | R02237 | 0.9 | 10 | 3.29455 | -9.41977e-05 | 0.999935 | biosynthesis | pure_linear |
| folC | R02237 | 1.0 | 0 | 3.29476 | -1.46711e-10 | 1.0 | biosynthesis | pure_linear |
| folA | R00936 | 0.05 | 95 | 3.23104 | -0.0281769 | 0.980659 | thf_cycle | loop_entry |
| folA | R00936 | 0.1 | 90 | 3.25863 | -0.0159104 | 0.989032 | thf_cycle | loop_entry |
| folA | R00936 | 0.2 | 80 | 3.27633 | -0.00809532 | 0.994404 | thf_cycle | loop_entry |
| folA | R00936 | 0.3 | 70 | 3.28333 | -0.00501512 | 0.99653 | thf_cycle | loop_entry |
| folA | R00936 | 0.5 | 50 | 3.28957 | -0.0022766 | 0.998423 | thf_cycle | loop_entry |
| folA | R00936 | 0.7 | 30 | 3.29247 | -0.00100358 | 0.999305 | thf_cycle | loop_entry |
| folA | R00936 | 0.9 | 10 | 3.29416 | -0.000264603 | 0.999817 | thf_cycle | loop_entry |
| folA | R00936 | 1.0 | 0 | 3.29476 | -1.46711e-10 | 1.0 | thf_cycle | loop_entry |
| glyA | R00945 | 0.05 | 95 | 3.29477 | 1.29975e-06 | 1.0 | thf_cycle | core_loop |
| glyA | R00945 | 0.1 | 90 | 3.29477 | 1.23125e-06 | 1.0 | thf_cycle | core_loop |
| glyA | R00945 | 0.2 | 80 | 3.29477 | 1.09428e-06 | 1.0 | thf_cycle | core_loop |
| glyA | R00945 | 0.3 | 70 | 3.29477 | 9.57337e-07 | 1.0 | thf_cycle | core_loop |
| glyA | R00945 | 0.5 | 50 | 3.29477 | 6.83578e-07 | 1.0 | thf_cycle | core_loop |
| glyA | R00945 | 0.7 | 30 | 3.29477 | 4.09916e-07 | 1.0 | thf_cycle | core_loop |
| glyA | R00945 | 0.9 | 10 | 3.29476 | 1.36531e-07 | 1.0 | thf_cycle | core_loop |
| glyA | R00945 | 1.0 | 0 | 3.29476 | -1.46711e-10 | 1.0 | thf_cycle | core_loop |
| thyA | R02101 | 0.05 | 95 | 0.169921 | -4.27724 | 0.0515729 | thf_cycle | loop_exit |
| thyA | R02101 | 0.1 | 90 | 0.339263 | -3.2797 | 0.10297 | thf_cycle | loop_exit |
| thyA | R02101 | 0.2 | 80 | 0.676236 | -2.28458 | 0.205246 | thf_cycle | loop_exit |
| thyA | R02101 | 0.3 | 70 | 1.01097 | -1.70444 | 0.30684 | thf_cycle | loop_exit |
| thyA | R02101 | 0.5 | 50 | 1.67387 | -0.976991 | 0.508038 | thf_cycle | loop_exit |
| thyA | R02101 | 0.7 | 30 | 2.3283 | -0.500897 | 0.706667 | thf_cycle | loop_exit |
| thyA | R02101 | 0.9 | 10 | 2.97459 | -0.147487 | 0.902822 | thf_cycle | loop_exit |
| thyA | R02101 | 1.0 | 0 | 3.29476 | -1.46711e-10 | 1.0 | thf_cycle | loop_exit |
| metF | R01224 | 0.05 | 95 | 78.1168 | 4.56739 | 23.7094 | thf_cycle | core_loop |
| metF | R01224 | 0.1 | 90 | 76.8905 | 4.54456 | 23.3372 | thf_cycle | core_loop |
| metF | R01224 | 0.2 | 80 | 71.5709 | 4.44113 | 21.7226 | thf_cycle | core_loop |
| metF | R01224 | 0.3 | 70 | 46.1435 | 3.80788 | 14.0051 | thf_cycle | core_loop |
| metF | R01224 | 0.5 | 50 | 10.0974 | 1.61574 | 3.06468 | thf_cycle | core_loop |
| metF | R01224 | 0.7 | 30 | 5.53025 | 0.74717 | 1.6785 | thf_cycle | core_loop |
| metF | R01224 | 0.9 | 10 | 3.80781 | 0.208785 | 1.15571 | thf_cycle | core_loop |
| metF | R01224 | 1.0 | 0 | 3.29476 | -1.46711e-10 | 1.0 | thf_cycle | core_loop |
| metH | R00946 | 0.05 | 95 | 3.29476 | -1.46711e-10 | 1.0 | methyl | loop_exit |
| metH | R00946 | 0.1 | 90 | 3.29476 | -1.46711e-10 | 1.0 | methyl | loop_exit |
| metH | R00946 | 0.2 | 80 | 3.29476 | -1.46711e-10 | 1.0 | methyl | loop_exit |
| metH | R00946 | 0.3 | 70 | 3.29476 | -1.46711e-10 | 1.0 | methyl | loop_exit |
| metH | R00946 | 0.5 | 50 | 3.29476 | -1.46711e-10 | 1.0 | methyl | loop_exit |
| metH | R00946 | 0.7 | 30 | 3.29476 | -1.46711e-10 | 1.0 | methyl | loop_exit |
| metH | R00946 | 0.9 | 10 | 3.29476 | -1.46711e-10 | 1.0 | methyl | loop_exit |
| metH | R00946 | 1.0 | 0 | 3.29476 | -1.46711e-10 | 1.0 | methyl | loop_exit |
| fhs | R04326 | 0.05 | 95 | 3.28324 | -0.00505716 | 0.996501 | thf_cycle | core_loop |
| fhs | R04326 | 0.1 | 90 | 3.28384 | -0.00479052 | 0.996685 | thf_cycle | core_loop |
| fhs | R04326 | 0.2 | 80 | 3.28506 | -0.0042574 | 0.997053 | thf_cycle | core_loop |
| fhs | R04326 | 0.3 | 70 | 3.28627 | -0.00372449 | 0.997422 | thf_cycle | core_loop |
| fhs | R04326 | 0.5 | 50 | 3.2887 | -0.0026593 | 0.998158 | thf_cycle | core_loop |
| fhs | R04326 | 0.7 | 30 | 3.29112 | -0.00159495 | 0.998895 | thf_cycle | core_loop |
| fhs | R04326 | 0.9 | 10 | 3.29355 | -0.000531443 | 0.999632 | thf_cycle | core_loop |
| fhs | R04326 | 1.0 | 0 | 3.29476 | -1.46711e-10 | 1.0 | thf_cycle | core_loop |
| folX | R11072 | 0.05 | 95 | 3.6577 | 0.150763 | 1.11016 | peripheral | pure_linear |
| folX | R11072 | 0.1 | 90 | 3.63918 | 0.14344 | 1.10454 | peripheral | pure_linear |
| folX | R11072 | 0.2 | 80 | 3.60194 | 0.128601 | 1.09323 | peripheral | pure_linear |
| folX | R11072 | 0.3 | 70 | 3.56444 | 0.113501 | 1.08185 | peripheral | pure_linear |
| folX | R11072 | 0.5 | 50 | 3.48865 | 0.0824951 | 1.05885 | peripheral | pure_linear |
| folX | R11072 | 0.7 | 30 | 3.41184 | 0.0503755 | 1.03553 | peripheral | pure_linear |
| folX | R11072 | 0.9 | 10 | 3.33403 | 0.017093 | 1.01192 | peripheral | pure_linear |
| folX | R11072 | 1.0 | 0 | 3.29476 | -1.46711e-10 | 1.0 | peripheral | pure_linear |
| gcv | R01221 | 0.05 | 95 | 3.33342 | 0.0168269 | 1.01173 | methyl | loop_entry |
| gcv | R01221 | 0.1 | 90 | 3.31394 | 0.00837098 | 1.00582 | methyl | loop_entry |
| gcv | R01221 | 0.2 | 80 | 3.30349 | 0.00381529 | 1.00265 | methyl | loop_entry |
| gcv | R01221 | 0.3 | 70 | 3.29989 | 0.00224457 | 1.00156 | methyl | loop_entry |
| gcv | R01221 | 0.5 | 50 | 3.29698 | 0.000968555 | 1.00067 | methyl | loop_entry |
| gcv | R01221 | 0.7 | 30 | 3.29572 | 0.000416317 | 1.00029 | methyl | loop_entry |
| gcv | R01221 | 0.9 | 10 | 3.29501 | 0.00010811 | 1.00007 | methyl | loop_entry |
| gcv | R01221 | 1.0 | 0 | 3.29476 | -1.46711e-10 | 1.0 | methyl | loop_entry |
| folD_MTHFC | R01655 | 0.05 | 95 | 3.54301 | 0.104801 | 1.07535 | thf_cycle | core_loop |
| folD_MTHFC | R01655 | 0.1 | 90 | 3.40788 | 0.0486986 | 1.03433 | thf_cycle | core_loop |
| folD_MTHFC | R01655 | 0.2 | 80 | 3.3441 | 0.0214435 | 1.01497 | thf_cycle | core_loop |
| folD_MTHFC | R01655 | 0.3 | 70 | 3.32337 | 0.0124706 | 1.00868 | thf_cycle | core_loop |
| folD_MTHFC | R01655 | 0.5 | 50 | 3.30696 | 0.00533156 | 1.0037 | thf_cycle | core_loop |
| folD_MTHFC | R01655 | 0.7 | 30 | 3.29998 | 0.00228259 | 1.00158 | thf_cycle | core_loop |
| folD_MTHFC | R01655 | 0.9 | 10 | 3.29612 | 0.000591441 | 1.00041 | thf_cycle | core_loop |
| folD_MTHFC | R01655 | 1.0 | 0 | 3.29476 | -1.46711e-10 | 1.0 | thf_cycle | core_loop |
| folD_MTHFD | R01220 | 0.05 | 95 | 3.29451 | -0.000110796 | 0.999923 | thf_cycle | core_loop |
| folD_MTHFD | R01220 | 0.1 | 90 | 3.29464 | -5.27617e-05 | 0.999963 | thf_cycle | core_loop |
| folD_MTHFD | R01220 | 0.2 | 80 | 3.29471 | -2.35122e-05 | 0.999984 | thf_cycle | core_loop |
| folD_MTHFD | R01220 | 0.3 | 70 | 3.29473 | -1.37277e-05 | 0.99999 | thf_cycle | core_loop |
| folD_MTHFD | R01220 | 0.5 | 50 | 3.29475 | -5.88758e-06 | 0.999996 | thf_cycle | core_loop |
| folD_MTHFD | R01220 | 0.7 | 30 | 3.29476 | -2.52409e-06 | 0.999998 | thf_cycle | core_loop |
| folD_MTHFD | R01220 | 0.9 | 10 | 3.29476 | -6.54671e-07 | 1.0 | thf_cycle | core_loop |
| folD_MTHFD | R01220 | 1.0 | 0 | 3.29476 | -1.46711e-10 | 1.0 | thf_cycle | core_loop |
| folC_FPGS | R04241 | 0.05 | 95 | 3.30271 | 0.00347611 | 1.00241 | peripheral | pure_linear |
| folC_FPGS | R04241 | 0.1 | 90 | 3.30229 | 0.00329292 | 1.00229 | peripheral | pure_linear |
| folC_FPGS | R04241 | 0.2 | 80 | 3.30145 | 0.00292663 | 1.00203 | peripheral | pure_linear |
| folC_FPGS | R04241 | 0.3 | 70 | 3.30062 | 0.00256045 | 1.00178 | peripheral | pure_linear |
| folC_FPGS | R04241 | 0.5 | 50 | 3.29894 | 0.00182838 | 1.00127 | peripheral | pure_linear |
| folC_FPGS | R04241 | 0.7 | 30 | 3.29727 | 0.00109672 | 1.00076 | peripheral | pure_linear |
| folC_FPGS | R04241 | 0.9 | 10 | 3.2956 | 0.000365472 | 1.00025 | peripheral | pure_linear |
| folC_FPGS | R04241 | 1.0 | 0 | 3.29476 | -1.46711e-10 | 1.0 | peripheral | pure_linear |
| purN | R04325 | 0.05 | 95 | 3.29474 | -1.28022e-05 | 0.999991 | peripheral | loop_exit |
| purN | R04325 | 0.1 | 90 | 3.29474 | -1.21284e-05 | 0.999992 | peripheral | loop_exit |
| purN | R04325 | 0.2 | 80 | 3.29474 | -1.07808e-05 | 0.999993 | peripheral | loop_exit |
| purN | R04325 | 0.3 | 70 | 3.29474 | -9.43328e-06 | 0.999993 | peripheral | loop_exit |
| purN | R04325 | 0.5 | 50 | 3.29475 | -6.73809e-06 | 0.999995 | peripheral | loop_exit |
| purN | R04325 | 0.7 | 30 | 3.29476 | -4.04293e-06 | 0.999997 | peripheral | loop_exit |
| purN | R04325 | 0.9 | 10 | 3.29476 | -1.34774e-06 | 0.999999 | peripheral | loop_exit |
| purN | R04325 | 1.0 | 0 | 3.29476 | -1.46711e-10 | 1.0 | peripheral | loop_exit |
| purH | R04560 | 0.05 | 95 | 3.29435 | -0.000182414 | 0.999874 | peripheral | pure_linear |
| purH | R04560 | 0.1 | 90 | 3.29437 | -0.000172814 | 0.99988 | peripheral | pure_linear |
| purH | R04560 | 0.2 | 80 | 3.29441 | -0.000153614 | 0.999894 | peripheral | pure_linear |
| purH | R04560 | 0.3 | 70 | 3.29446 | -0.000134413 | 0.999907 | peripheral | pure_linear |
| purH | R04560 | 0.5 | 50 | 3.29455 | -9.60109e-05 | 0.999933 | peripheral | pure_linear |
| purH | R04560 | 0.7 | 30 | 3.29463 | -5.76074e-05 | 0.99996 | peripheral | pure_linear |
| purH | R04560 | 0.9 | 10 | 3.29472 | -1.92028e-05 | 0.999987 | peripheral | pure_linear |
| purH | R04560 | 1.0 | 0 | 3.29476 | -1.46711e-10 | 1.0 | peripheral | pure_linear |
| purU | R00944 | 0.05 | 95 | 3.28369 | -0.00485553 | 0.99664 | peripheral | pure_linear |
| purU | R00944 | 0.1 | 90 | 3.28427 | -0.00460044 | 0.996816 | peripheral | pure_linear |
| purU | R00944 | 0.2 | 80 | 3.28544 | -0.0040901 | 0.997169 | peripheral | pure_linear |
| purU | R00944 | 0.3 | 70 | 3.2866 | -0.00357956 | 0.997522 | peripheral | pure_linear |
| purU | R00944 | 0.5 | 50 | 3.28893 | -0.00255786 | 0.998229 | peripheral | pure_linear |
| purU | R00944 | 0.7 | 30 | 3.29126 | -0.00153533 | 0.998936 | peripheral | pure_linear |
| purU | R00944 | 0.9 | 10 | 3.2936 | -0.000511985 | 0.999645 | peripheral | pure_linear |
| purU | R00944 | 1.0 | 0 | 3.29476 | -1.46711e-10 | 1.0 | peripheral | pure_linear |
| fau | R02301 | 0.05 | 95 | 3.29476 | -1.46711e-10 | 1.0 | peripheral | pure_linear |
| fau | R02301 | 0.1 | 90 | 3.29476 | -1.46711e-10 | 1.0 | peripheral | pure_linear |
| fau | R02301 | 0.2 | 80 | 3.29476 | -1.46711e-10 | 1.0 | peripheral | pure_linear |
| fau | R02301 | 0.3 | 70 | 3.29476 | -1.46711e-10 | 1.0 | peripheral | pure_linear |
| fau | R02301 | 0.5 | 50 | 3.29476 | -1.46711e-10 | 1.0 | peripheral | pure_linear |
| fau | R02301 | 0.7 | 30 | 3.29476 | -1.46711e-10 | 1.0 | peripheral | pure_linear |
| fau | R02301 | 0.9 | 10 | 3.29476 | -1.46711e-10 | 1.0 | peripheral | pure_linear |
| fau | R02301 | 1.0 | 0 | 3.29476 | -1.46711e-10 | 1.0 | peripheral | pure_linear |
| metK | R00177 | 0.05 | 95 | 3.30581 | 0.00482822 | 1.00335 | methyl | loop_exit |
| metK | R00177 | 0.1 | 90 | 3.30523 | 0.00457623 | 1.00318 | methyl | loop_exit |
| metK | R00177 | 0.2 | 80 | 3.30408 | 0.00407145 | 1.00283 | methyl | loop_exit |
| metK | R00177 | 0.3 | 70 | 3.30292 | 0.00356566 | 1.00247 | methyl | loop_exit |
| metK | R00177 | 0.5 | 50 | 3.3006 | 0.00255119 | 1.00177 | methyl | loop_exit |
| metK | R00177 | 0.7 | 30 | 3.29827 | 0.00153315 | 1.00106 | methyl | loop_exit |
| metK | R00177 | 0.9 | 10 | 3.29593 | 0.000511816 | 1.00035 | methyl | loop_exit |
| metK | R00177 | 1.0 | 0 | 3.29476 | -1.46711e-10 | 1.0 | methyl | loop_exit |
| folM | R_FOLM | 0.05 | 95 | 3.29437 | -0.000174477 | 0.999879 | peripheral | pure_linear |
| folM | R_FOLM | 0.1 | 90 | 3.29439 | -0.000164742 | 0.999886 | peripheral | pure_linear |
| folM | R_FOLM | 0.2 | 80 | 3.29443 | -0.000145465 | 0.999899 | peripheral | pure_linear |
| folM | R_FOLM | 0.3 | 70 | 3.29448 | -0.000126441 | 0.999912 | peripheral | pure_linear |
| folM | R_FOLM | 0.5 | 50 | 3.29456 | -8.91359e-05 | 0.999938 | peripheral | pure_linear |
| folM | R_FOLM | 0.7 | 30 | 3.29464 | -5.27911e-05 | 0.999963 | peripheral | pure_linear |
| folM | R_FOLM | 0.9 | 10 | 3.29472 | -1.73725e-05 | 0.999988 | peripheral | pure_linear |
| folM | R_FOLM | 1.0 | 0 | 3.29476 | -1.46711e-10 | 1.0 | peripheral | pure_linear |
| sah_hydrolase | R_SAH_HYD | 0.05 | 95 | 3.29476 | -1.46676e-10 | 1.0 | methyl | loop_exit |
| sah_hydrolase | R_SAH_HYD | 0.1 | 90 | 3.29476 | -1.47502e-10 | 1.0 | methyl | loop_exit |
| sah_hydrolase | R_SAH_HYD | 0.2 | 80 | 3.29476 | -1.46678e-10 | 1.0 | methyl | loop_exit |
| sah_hydrolase | R_SAH_HYD | 0.3 | 70 | 3.29476 | -1.46617e-10 | 1.0 | methyl | loop_exit |
| sah_hydrolase | R_SAH_HYD | 0.5 | 50 | 3.29476 | -1.46676e-10 | 1.0 | methyl | loop_exit |
| sah_hydrolase | R_SAH_HYD | 0.7 | 30 | 3.29476 | -1.46676e-10 | 1.0 | methyl | loop_exit |
| sah_hydrolase | R_SAH_HYD | 0.9 | 10 | 3.29476 | -1.47246e-10 | 1.0 | methyl | loop_exit |
| sah_hydrolase | R_SAH_HYD | 1.0 | 0 | 3.29476 | -1.46711e-10 | 1.0 | methyl | loop_exit |

Table S4. Full single-node knockdown scan across all 38 nodes in the folate and one-carbon network. Each node was knocked down by reducing its maximum velocity (V_max) to the indicated fraction of wild-type (0.05–1.0, corresponding to 0–95% inhibition). Steady-state dTMP concentration (μM), log₂ fold change relative to wild type, and fold change are reported for each knockdown level. The pathway column indicates the module assigned to each node (biosynthesis, thf_cycle, methyl, peripheral), and the refined column classifies each node by topological role (pure_linear, core_loop, loop_entry, loop_exit, other). The table has 208 rows = 26 nodes × 8 knockdown levels, truncated here to the first 200 rows. Full data are provided in the source CSV file.

*(Showing first 200 of 200 rows — full data in source CSV.)*

Supplementary Table S5

| pathway | mean | median | std | count |
| --- | --- | --- | --- | --- |
| biosynthesis | -1.39105 | -1.88879 | 1.0756 | 7 |
| methyl | 0.00148628 | -1.46678e-10 | 0.00225252 | 5 |
| peripheral | 0.018161 | -1.07808e-05 | 0.048742 | 7 |
| thf_cycle | 0.309374 | -2.35122e-05 | 2.01136 | 7 |

Table S5. Module-level pathway summaries for the folate, one-carbon, PreQ1, and MoCo modules. For each pathway category (biosynthesis, methyl, peripheral, thf_cycle), the mean, median, and standard deviation of the log₂ fold change in dTMP flux across all knockdown conditions are reported, along with the number of nodes assigned to each category. These summaries quantify the average response of each pathway category to single-node knockdown and show that pathway control is distributed unequally across modules, with the biosynthesis module exhibiting the largest negative response.

Supplementary Table S6

| Inhibition_level | Target_inhibition | TMP_single_dose | SMX_single_dose | TMP_combination_dose | SMX_combination_dose | Combination_Index | Interpretation |
| --- | --- | --- | --- | --- | --- | --- | --- |
| 0.5 | 0.5 | 0.340425 | 1.48543 | 0.009766 | 0.4 | 0.29797 | Synergy |
| 0.9 | 0.9 | 0.688193 | 3.26189 | 0.4 | 5.0 | 2.11409 | Antagonism |

Table S6. Chou-Talalay combination indices (CI) at IC₅₀ and IC₉₀ for the SMX × TMP combination. For each inhibition level (50% and 90%), the table reports the target inhibition, single-agent doses at that effect level (TMP_single_dose, SMX_single_dose), the combination doses required to achieve the same effect (TMP_combination_dose, SMX_combination_dose), and the resulting combination index. CI values were computed using the median-effect equation. Following standard interpretation, CI < 1 indicates synergy, CI = 1 additivity, and CI > 1 antagonism. At IC₅₀ the combination is strongly synergistic (CI = 0.298), while at IC₉₀ the interaction shifts to antagonism (CI = 2.114), demonstrating dose-dependent synergy.

Supplementary Table S7

| Condition | Well | Replicate | TMP | SMX | Observed_fractional_growth | TMP_fractional_growth | SMX_fractional_growth | Bliss_predicted | Bliss_residual |
| --- | --- | --- | --- | --- | --- | --- | --- | --- | --- |
| TMP+SMX arm 1 | TMP+SMX arm 1_W1 | 1 | 0.4 | 10.0 | 0.277519 | 0.657536 | 0.375819 | 0.247114 | 0.0304053 |
| TMP+SMX arm 1 | TMP+SMX arm 1_W10 | 1 | 0.4 | 0.019531 | 0.282171 | 0.657536 | 1.08001 | 0.710144 | -0.427973 |
| TMP+SMX arm 1 | TMP+SMX arm 1_W11 | 1 | 0.4 | 0.009766 | 0.297984 | 0.657536 | 1.08001 | 0.710146 | -0.412161 |
| TMP+SMX arm 1 | TMP+SMX arm 1_W2 | 1 | 0.4 | 5.0 | 0.270078 | 0.657536 | 0.395463 | 0.260031 | 0.0100465 |
| TMP+SMX arm 1 | TMP+SMX arm 1_W3 | 1 | 0.4 | 2.5 | 0.27876 | 0.657536 | 0.506391 | 0.33297 | -0.0542105 |
| TMP+SMX arm 1 | TMP+SMX arm 1_W4 | 1 | 0.4 | 1.25 | 0.275349 | 0.657536 | 0.80986 | 0.532512 | -0.257163 |
| TMP+SMX arm 1 | TMP+SMX arm 1_W5 | 1 | 0.4 | 0.625 | 0.270388 | 0.657536 | 1.02213 | 0.672088 | -0.4017 |
| TMP+SMX arm 1 | TMP+SMX arm 1_W6 | 1 | 0.4 | 0.3125 | 0.271318 | 0.657536 | 1.07103 | 0.704243 | -0.432925 |
| TMP+SMX arm 1 | TMP+SMX arm 1_W7 | 1 | 0.4 | 0.15625 | 0.273798 | 0.657536 | 1.0787 | 0.709285 | -0.435487 |
| TMP+SMX arm 1 | TMP+SMX arm 1_W8 | 1 | 0.4 | 0.078125 | 0.285581 | 0.657536 | 1.07982 | 0.710022 | -0.42444 |
| TMP+SMX arm 1 | TMP+SMX arm 1_W9 | 1 | 0.4 | 0.039063 | 0.299225 | 0.657536 | 1.07998 | 0.710128 | -0.410903 |
| TMP+SMX arm 2 | TMP+SMX arm 2_W1 | 1 | 10.0 | 0.4 | 0.275659 | 0.383478 | 1.06234 | 0.407385 | -0.131726 |
| TMP+SMX arm 2 | TMP+SMX arm 2_W10 | 1 | 0.019531 | 0.4 | 1.02264 | 1.11086 | 1.06234 | 1.18012 | -0.157481 |
| TMP+SMX arm 2 | TMP+SMX arm 2_W11 | 1 | 0.009766 | 0.4 | 0.43876 | 1.11095 | 1.06234 | 1.18021 | -0.741448 |
| TMP+SMX arm 2 | TMP+SMX arm 2_W2 | 1 | 5.0 | 0.4 | 0.273178 | 0.383624 | 1.06234 | 0.407541 | -0.134362 |
| TMP+SMX arm 2 | TMP+SMX arm 2_W3 | 1 | 2.5 | 0.4 | 0.575814 | 0.384897 | 1.06234 | 0.408893 | 0.166921 |
| TMP+SMX arm 2 | TMP+SMX arm 2_W4 | 1 | 1.25 | 0.4 | 0.27969 | 0.395791 | 1.06234 | 0.420466 | -0.140776 |
| TMP+SMX arm 2 | TMP+SMX arm 2_W5 | 1 | 0.625 | 0.4 | 0.273178 | 0.478397 | 1.06234 | 0.508221 | -0.235043 |
| TMP+SMX arm 2 | TMP+SMX arm 2_W6 | 1 | 0.3125 | 0.4 | 0.275039 | 0.795516 | 1.06234 | 0.845111 | -0.570072 |
| TMP+SMX arm 2 | TMP+SMX arm 2_W7 | 1 | 0.15625 | 0.4 | 0.28031 | 1.05215 | 1.06234 | 1.11774 | -0.837429 |
| TMP+SMX arm 2 | TMP+SMX arm 2_W8 | 1 | 0.078125 | 0.4 | 0.276279 | 1.10368 | 1.06234 | 1.17249 | -0.896209 |
| TMP+SMX arm 2 | TMP+SMX arm 2_W9 | 1 | 0.039063 | 0.4 | 0.28062 | 1.11012 | 1.06234 | 1.17932 | -0.898703 |

Table S7. Per-dose Bliss residuals across the 22-dose SMX × TMP grid. Bliss independence predicts that the combined fractional growth equals the product of the single-agent fractional growths at the same doses: E_Bliss = E_TMP × E_SMX. The Bliss residual is defined as Δ_Bliss = E_observed − E_Bliss. Negative residuals indicate synergy (combination inhibits growth more than the product of single-agent effects); positive residuals indicate antagonism. The table reports all 22 measured dose pairs across both fixed-dose arms (Arm 1: TMP fixed, SMX titrated; Arm 2: SMX fixed, TMP titrated), including the condition, well identifier, replicate, individual drug concentrations, observed combination growth, single-agent predictions, and residual.

Supplementary Table S8

| Condition | Well | Replicate | mu_max | mu_exp_start_h | mu_exp_end_h | OD_16h | Lag_to_OD0.1_h | AUC_0_16h | TMP | SMX |
| --- | --- | --- | --- | --- | --- | --- | --- | --- | --- | --- |
| SMX | SMX_W1 | 1 | 0.00452745 | 8.25 | 9.0 | 0.1391 | 12.5988 | 1.52869 | 0.0 | 10.0 |
| SMX | SMX_W10 | 1 | 0.320458 | 8.75 | 9.5 | 0.3549 | 6.76613 | 3.05335 | 0.0 | 0.019531 |
| SMX | SMX_W11 | 1 | 0.300759 | 9.5 | 10.25 | 0.3501 | 6.46429 | 3.0838 | 0.0 | 0.009766 |
| SMX | SMX_W2 | 1 | 0.00841222 | 9.75 | 10.5 | 0.1071 | 13.3333 | 1.5137 | 0.0 | 5.0 |
| SMX | SMX_W3 | 1 | 0.00447178 | 3.75 | 4.5 | 0.1645 | 10.625 | 1.62985 | 0.0 | 2.5 |
| SMX | SMX_W4 | 1 | 0.124346 | 15.25 | 16.0 | 0.2658 | 8.01667 | 2.03619 | 0.0 | 1.25 |
| SMX | SMX_W5 | 1 | 0.0046811 | 2.75 | 3.5 | 0.3217 | 8.01136 | 2.31769 | 0.0 | 0.625 |
| SMX | SMX_W6 | 1 | 0.192203 | 11.75 | 12.5 | 0.3375 | 7.66667 | 2.45824 | 0.0 | 0.3125 |
| SMX | SMX_W7 | 1 | 0.219628 | 10.5 | 11.25 | 0.3352 | 6.32692 | 3.02015 | 0.0 | 0.15625 |
| SMX | SMX_W8 | 1 | 0.00464846 | 1.5 | 2.25 | 0.3631 | 6.79808 | 3.03819 | 0.0 | 0.078125 |
| SMX | SMX_W9 | 1 | 0.311971 | 8.75 | 10.25 | 0.3503 | 6.86607 | 3.00521 | 0.0 | 0.039063 |
| TMP | TMP_W1 | 1 | 0.108454 | 6.25 | 7.5 | 0.202 | 0.0 | 2.26057 | 10.0 | 0.0 |
| TMP | TMP_W10 | 1 | 0.275026 | 9.5 | 10.75 | 0.3483 | 6.80645 | 2.94347 | 0.019531 | 0.0 |
| TMP | TMP_W11 | 1 | 0.295442 | 9.75 | 10.5 | 0.3517 | 6.625 | 3.04962 | 0.009766 | 0.0 |
| TMP | TMP_W2 | 1 | 0.00568905 | 3.25 | 4.0 | 0.0924 |  | 1.46705 | 5.0 | 0.0 |
| TMP | TMP_W3 | 1 | 0.00612709 | 14.25 | 15.75 | 0.089 |  | 1.39136 | 2.5 | 0.0 |
| TMP | TMP_W4 | 1 | 0.00438357 | 3.0 | 3.75 | 0.1023 | 15.15 | 1.49929 | 1.25 | 0.0 |
| TMP | TMP_W5 | 1 | 0.00457405 | 3.0 | 3.75 | 0.1733 | 10.3542 | 1.65431 | 0.625 | 0.0 |
| TMP | TMP_W6 | 1 | 0.00463769 | 3.0 | 3.75 | 0.246 | 9.60714 | 1.8476 | 0.3125 | 0.0 |
| TMP | TMP_W7 | 1 | 0.00469209 | 2.5 | 3.25 | 0.3496 | 8.45 | 2.25154 | 0.15625 | 0.0 |
| TMP | TMP_W8 | 1 | 0.157445 | 13.25 | 14.0 | 0.3736 | 7.31818 | 2.61141 | 0.078125 | 0.0 |
| TMP | TMP_W9 | 1 | 0.0182028 | 4.0 | 4.75 | 0.3516 | 7.17188 | 2.76016 | 0.039063 | 0.0 |
| TMP+SMX arm 1 | TMP+SMX arm 1_W1 | 1 | 0.00219607 | 3.25 | 5.0 | 0.0895 |  | 1.42698 | 0.4 | 10.0 |
| TMP+SMX arm 1 | TMP+SMX arm 1_W10 | 1 | 0.00453258 | 1.25 | 2.0 | 0.091 |  | 1.42904 | 0.4 | 0.019531 |
| TMP+SMX arm 1 | TMP+SMX arm 1_W11 | 1 | 0.0184199 | 13.5 | 14.75 | 0.0961 |  | 1.4282 | 0.4 | 0.009766 |
| TMP+SMX arm 1 | TMP+SMX arm 1_W2 | 1 | 0.00463231 | 2.5 | 3.25 | 0.0871 |  | 1.38742 | 0.4 | 5.0 |
| TMP+SMX arm 1 | TMP+SMX arm 1_W3 | 1 | 0.00780319 | 0.5 | 1.25 | 0.0899 |  | 1.40647 | 0.4 | 2.5 |
| TMP+SMX arm 1 | TMP+SMX arm 1_W4 | 1 | 0.00459506 | 3.5 | 4.25 | 0.0888 |  | 1.40201 | 0.4 | 1.25 |
| TMP+SMX arm 1 | TMP+SMX arm 1_W5 | 1 | 0.00268461 | 13.25 | 15.0 | 0.0872 |  | 1.3808 | 0.4 | 0.625 |
| TMP+SMX arm 1 | TMP+SMX arm 1_W6 | 1 | 0.00457928 | 12.5 | 13.25 | 0.0875 |  | 1.38796 | 0.4 | 0.3125 |
| TMP+SMX arm 1 | TMP+SMX arm 1_W7 | 1 | 0.00590214 | 15.25 | 16.0 | 0.0883 |  | 1.38475 | 0.4 | 0.15625 |
| TMP+SMX arm 1 | TMP+SMX arm 1_W8 | 1 | 0.00462161 | 8.0 | 8.75 | 0.0921 |  | 1.39425 | 0.4 | 0.078125 |
| TMP+SMX arm 1 | TMP+SMX arm 1_W9 | 1 | 0.0312249 | 15.0 | 15.75 | 0.0965 |  | 1.40255 | 0.4 | 0.039063 |
| TMP+SMX arm 2 | TMP+SMX arm 2_W1 | 1 | 0.00467017 | 1.0 | 1.75 | 0.0889 | 13.1657 | 1.41362 | 10.0 | 0.4 |
| TMP+SMX arm 2 | TMP+SMX arm 2_W10 | 1 | 0.563773 | 8.5 | 9.25 | 0.3298 | 0.319249 | 3.91389 | 0.019531 | 0.4 |
| TMP+SMX arm 2 | TMP+SMX arm 2_W11 | 1 | 0.120624 | 8.5 | 9.25 | 0.1415 | 2.63636 | 1.72936 | 0.009766 | 0.4 |
| TMP+SMX arm 2 | TMP+SMX arm 2_W2 | 1 | 0.00273349 | 14.25 | 15.0 | 0.0881 | 8.17935 | 1.42332 | 5.0 | 0.4 |
| TMP+SMX arm 2 | TMP+SMX arm 2_W3 | 1 | 0.303687 | 9.5 | 10.25 | 0.1857 | 1.06514 | 1.93904 | 2.5 | 0.4 |
| TMP+SMX arm 2 | TMP+SMX arm 2_W4 | 1 | 0.00289968 | 12.5 | 14.5 | 0.0902 |  | 1.42621 | 1.25 | 0.4 |
| TMP+SMX arm 2 | TMP+SMX arm 2_W5 | 1 | 0.00456882 | 13.5 | 14.25 | 0.0881 |  | 1.39251 | 0.625 | 0.4 |
| TMP+SMX arm 2 | TMP+SMX arm 2_W6 | 1 | 0.00372876 | 2.5 | 3.75 | 0.0887 |  | 1.42481 | 0.3125 | 0.4 |
| TMP+SMX arm 2 | TMP+SMX arm 2_W7 | 1 | 0.00443705 | 14.75 | 15.5 | 0.0904 |  | 1.40738 | 0.15625 | 0.4 |
| TMP+SMX arm 2 | TMP+SMX arm 2_W8 | 1 | 0.00723574 | 14.0 | 14.75 | 0.0891 |  | 1.39404 | 0.078125 | 0.4 |
| TMP+SMX arm 2 | TMP+SMX arm 2_W9 | 1 | 0.00457928 | 5.5 | 6.25 | 0.0905 |  | 1.43175 | 0.039063 | 0.4 |
| WT | WT_W12_SMX | 1 | 0.311689 | 9.0 | 9.75 | 0.3559 | 6.4375 | 3.10044 | 0.0 | 0.0 |
| WT | WT_W12_TMP | 1 | 0.00435968 | 4.0 | 4.75 | 0.2998 | 8.51667 | 2.08974 | 0.0 | 0.0 |
| WT | WT_W12_TMP+SMX_arm_1 | 1 | 0.318237 | 9.0 | 9.75 | 0.3425 | 4.74299 | 3.0626 | 0.0 | 0.0 |
| WT | WT_W12_TMP+SMX_arm_2 | 1 | 0.300778 | 9.0 | 9.75 | 0.2918 | 6.72368 | 2.74879 | 0.0 | 0.0 |

Table S8. Per-well growth metrics for all experimental plates. For each well, the following metrics were extracted from the 16-hour growth curves: maximum specific growth rate (μ_max, h⁻¹), start and end times of the exponential growth window (mu_exp_start_h, mu_exp_end_h), OD₆₀₀ at 16 h, lag time to reach OD 0.1 (Lag_to_OD0.1_h, h), and area under the growth curve from 0 to 16 h (AUC_0_16h). TMP and SMX concentrations (μg/mL) are reported for each well. Data cover all four conditions (SMX alone, TMP alone, Arm 1, Arm 2) across the dose series, with 48 wells in total.

Supplementary Table S9

| excluded | diff |
| --- | --- |
| folC | -1.74338 |
| folB | -1.70419 |
| thyA | -1.64575 |
| folK | -1.6421 |
| folA | -1.51928 |
| fhs | -1.51907 |
| purU | -1.51906 |
| sam_methyltransferase | -1.51886 |
| purH | -1.51884 |
| folM | -1.51884 |
| folD_MTHFD | -1.51883 |
| purN | -1.51883 |
| metH | -1.51883 |
| fau | -1.51883 |
| sah_hydrolase | -1.51883 |
| glyA | -1.51883 |
| folC_FPGS | -1.51867 |
| gcv | -1.51862 |
| metK | -1.51861 |
| folD_MTHFC | -1.51764 |
| folX | -1.51169 |
| folP | -1.42915 |
| phoA | -1.40652 |
| nudB | -1.35168 |
| folE | -1.30773 |
| metF | -1.2721 |

Table S9. Leave-one-out robustness analysis of the single-agent Hill fits. Each row reports the effect of excluding a single experimental dose point on the fitted model. The excluded column lists the omitted drug or condition, and diff reports the change in fitted parameters relative to the full-data fit. Small absolute values indicate that the fits are robust to the removal of individual points; larger values flag influential observations. The table supports the reliability of the IC₅₀ estimates used in the synergy analysis.

Supplementary Table S9a

| Rank_Full | Node_A | Node_B | Module_A | Module_B | Raw_Synergy | Corrected_Synergy | Additive_dTMP | Below_Floor | Correction_Type | Folate_Specific | Rank_Folate |
| --- | --- | --- | --- | --- | --- | --- | --- | --- | --- | --- | --- |
| 2 | folK | gcv | Biosynthesis | One-carbon | 2.686021197700209 | 2.024019355754905 | 0.0429215692419506 | True | floor+bifurcation | True | 1.0 |
| 3 | folP | folA | Biosynthesis | Folate cycle | 3.904236561941905 | 1.8089573103777097 | 0.0263914079516334 | True | floor+bifurcation | True | 2.0 |
| 5 | folB | folA | Biosynthesis | Folate cycle | 3.904408775210453 | 1.7847733593149282 | 0.0260374327135672 | True | floor+bifurcation | True | 3.0 |
| 7 | folB | fhs | Biosynthesis | One-carbon | 1.433614285284234 | 1.424469470366753 | 0.0565967107575755 | True | floor+bifurcation | True | 4.0 |
| 8 | folC_FPGS | gcv | Polyglutamation | One-carbon | 3.736922411972877 | 1.4242377492190252 | 1.961755511942931 | False | bifurcation only | True | 5.0 |
| 10 | folB | folK | Biosynthesis | Biosynthesis | 4.626533773751336 | 1.297741917590488 | 0.0159772842573957 | True | floor+bifurcation | True | 6.0 |
| 13 | folK | folP | Biosynthesis | Biosynthesis | 4.50030190513129 | 1.2794951523821232 | 0.016194493191198 | True | floor+bifurcation | True | 7.0 |
| 16 | folA | gcv | Folate cycle | One-carbon | 2.903051576533877 | 1.1064280141279426 | 0.0699472734598617 | False | bifurcation only | True | 8.0 |
| 19 | metF | gcv | Methyl | One-carbon | 2.677325989548021 | 1.020398156799395 | 0.6583079824941427 | False | bifurcation only | True | 9.0 |
| 54 | gcv | fau | One-carbon | One-carbon | 1.000000000000091 | 1.000000000000091 | 0.2742246291054555 | False | none | True | 10.0 |
| 93 | fau | sah_hydrolase | One-carbon | Methyl | 1.000000000000001 | 1.000000000000001 | 2.317185633030864 | False | none | True | 11.0 |
| 100 | fau | sam_methyltransferase | One-carbon | Methyl | 1.000000000000001 | 1.000000000000001 | 2.3161839492126246 | False | none | True | 12.0 |
| 119 | fau | metK | One-carbon | Methyl | 1.0 | 1.0 | 2.3202017258732983 | False | none | True | 13.0 |
| 132 | folX | fau | Biosynthesis | One-carbon | 0.9999999999999998 | 0.9999999999999998 | 2.6194880971841448 | False | none | True | 14.0 |
| 226 | folX | sam_methyltransferase | Biosynthesis | Methyl | 0.9995342854431376 | 0.9995342854431376 | 2.638889945182197 | False | none | True | 15.0 |
| 228 | folX | sah_hydrolase | Biosynthesis | Methyl | 0.9994333069888156 | 0.9994333069888156 | 2.640031190184392 | False | none | True | 16.0 |
| 232 | folX | metK | Biosynthesis | Methyl | 0.998721299526012 | 0.998721299526012 | 2.643467504937519 | False | none | True | 17.0 |
| 239 | metK | sam_methyltransferase | Methyl | Methyl | 0.9957705952831392 | 0.9957705952831392 | 2.337386839735279 | False | none | True | 18.0 |
| 241 | metK | sah_hydrolase | Methyl | Methyl | 0.9955212503297688 | 0.9955212503297688 | 2.3383976932018715 | False | none | True | 19.0 |
| 243 | gcv | metK | One-carbon | Methyl | 0.9937902223925136 | 0.9937902223925136 | 0.2767349456075256 | False | none | True | 20.0 |
| 244 | sam_methyltransferase | sah_hydrolase | Methyl | Methyl | 0.9930500586986588 | 0.9930500586986588 | 2.334348407499533 | False | none | True | 21.0 |
| 245 | gcv | sam_methyltransferase | One-carbon | Methyl | 0.9926482999967576 | 0.9926482999967576 | 0.2762557376174376 | False | none | True | 22.0 |
| 246 | gcv | sah_hydrolase | One-carbon | Methyl | 0.9925194706352574 | 0.9925194706352574 | 0.2763752103830446 | False | none | True | 23.0 |
| 249 | folX | gcv | Biosynthesis | One-carbon | 0.9596965131380027 | 0.9596965131380027 | 0.312431409739155 | False | none | True | 24.0 |
| 258 | folP | gcv | Biosynthesis | One-carbon | 3.060121237042603 | 0.6629866037560325 | 0.0123406059891512 | True | floor+bifurcation | True | 25.0 |
| 259 | folB | gcv | Biosynthesis | One-carbon | 3.061882213270057 | 0.6544706887552105 | 0.0121750873873814 | True | floor+bifurcation | True | 26.0 |
| 260 | folB | folP | Biosynthesis | Biosynthesis | 7.410658654889547 | 0.5976546462241159 | 0.0045937129811293 | True | floor+bifurcation | True | 27.0 |
| 261 | folK | folA | Biosynthesis | Folate cycle | 3.773226849948493 | 0.5480873005969936 | 0.0917913305703479 | False | bifurcation only | True | 28.0 |
| 263 | folK | folC_FPGS | Biosynthesis | Polyglutamation | 3.702362757989354 | 0.5377938010498694 | 2.574398397362659 | False | bifurcation only | True | 29.0 |
| 267 | folC_DHFS | gcv | Biosynthesis | One-carbon | 1.331630802304018 | 0.5075189280322593 | 0.1928693099036315 | False | bifurcation only | True | 30.0 |
| 270 | metH | gcv | Methyl | One-carbon | 1.272867606867841 | 0.485122754930888 | 0.2805788290931392 | False | bifurcation only | True | 31.0 |
| 279 | folB | thyA | Biosynthesis | One-carbon | 0.6891205467169919 | 0.4036433785733653 | 0.0333636072210209 | True | floor+bifurcation | True | 32.0 |
| 282 | folP | thyA | Biosynthesis | One-carbon | 0.6798777048789388 | 0.4036433785733651 | 0.0338171807717902 | True | floor+bifurcation | True | 33.0 |
| 284 | folD_MTHFD | gcv | One-carbon | One-carbon | 1.040708031496403 | 0.3966407382780195 | 0.1909009145430366 | False | bifurcation only | True | 34.0 |
| 285 | folX | fhs | Biosynthesis | One-carbon | 1.012211192863499 | 0.3857798562901171 | 1.452358374603138 | False | bifurcation only | True | 35.0 |
| 288 | folX | folD_MTHFC | Biosynthesis | One-carbon | 1.010386007689407 | 0.385084231030163 | 1.5760001759744282 | False | bifurcation only | True | 36.0 |
| 289 | folK | sam_methyltransferase | Biosynthesis | Methyl | 1.006494895989436 | 0.3836012277567286 | 0.3625285229759353 | False | bifurcation only | True | 37.0 |
| 290 | folK | sah_hydrolase | Biosynthesis | Methyl | 1.006254548269248 | 0.3835096250263819 | 0.3626853062725463 | False | bifurcation only | True | 38.0 |
| 292 | fhs | sah_hydrolase | One-carbon | Methyl | 1.004927952280081 | 0.383004024995811 | 1.2847487122618022 | False | bifurcation only | True | 39.0 |
| 293 | fhs | sam_methyltransferase | One-carbon | Methyl | 1.004899752795141 | 0.3829932774430036 | 1.2841933350933823 | False | bifurcation only | True | 40.0 |
| 294 | folD_MTHFC | sah_hydrolase | One-carbon | Methyl | 1.004070738549732 | 0.38267731867994 | 1.3941216107634549 | False | bifurcation only | True | 41.0 |
| 295 | folD_MTHFC | sam_methyltransferase | One-carbon | Methyl | 1.004046152651626 | 0.3826679483585041 | 1.3935189533680992 | False | bifurcation only | True | 42.0 |
| 296 | fhs | metK | One-carbon | Methyl | 1.003581565709321 | 0.3824908822629103 | 1.2864209655936647 | False | bifurcation only | True | 43.0 |
| 297 | folD_MTHFD | sam_methyltransferase | One-carbon | Methyl | 1.003439433901661 | 0.3824367121561975 | 1.612406717795415 | False | bifurcation only | True | 44.0 |
| 298 | folD_MTHFD | sah_hydrolase | One-carbon | Methyl | 1.003367218052275 | 0.3824091888288579 | 1.613104037936237 | False | bifurcation only | True | 45.0 |
| 299 | folX | glyA | Biosynthesis | One-carbon | 1.003206374626415 | 0.3823478872406607 | 1.8716519237360945 | False | bifurcation only | True | 46.0 |
| 300 | folD_MTHFC | metK | One-carbon | Methyl | 1.002970129928434 | 0.3822578482781606 | 1.3959362259379016 | False | bifurcation only | True | 47.0 |
| 301 | glyA | sah_hydrolase | One-carbon | Methyl | 1.002780836257997 | 0.382185703566175 | 1.6556536188799391 | False | bifurcation only | True | 48.0 |
| 302 | glyA | sam_methyltransferase | One-carbon | Methyl | 1.002777465332738 | 0.3821844188194035 | 1.6549379052077158 | False | bifurcation only | True | 49.0 |
| 305 | glyA | metK | One-carbon | Methyl | 1.001909233646842 | 0.3818535132758046 | 1.657808649084802 | False | bifurcation only | True | 50.0 |
| 306 | folD_MTHFD | metK | One-carbon | Methyl | 1.001632310821829 | 0.3817479708273608 | 1.6152036847981752 | False | bifurcation only | True | 51.0 |
| 307 | folK | metK | Biosynthesis | Methyl | 1.001598896336193 | 0.3817352357029545 | 0.3631573843575783 | False | bifurcation only | True | 52.0 |
| 326 | folM | gcv | Folate cycle | One-carbon | 1.000017681404825 | 0.3811325938103457 | 0.2742190158497183 | False | bifurcation only | True | 53.0 |
| 339 | folC_FPGS | fau | Polyglutamation | One-carbon | 1.000000000003092 | 0.3811258549709932 | 16.447754780513936 | False | bifurcation only | True | 54.0 |
| 343 | thyA | fau | One-carbon | One-carbon | 1.000000000002705 | 0.3811258549708457 | 0.7514625993803472 | False | bifurcation only | True | 55.0 |
| 351 | folD_MTHFD | fau | One-carbon | One-carbon | 1.000000000001094 | 0.3811258549702317 | 1.6005518581007845 | False | bifurcation only | True | 56.0 |
| 357 | folK | fau | Biosynthesis | One-carbon | 1.000000000000935 | 0.3811258549701711 | 0.3598631131089649 | False | bifurcation only | True | 57.0 |
| 371 | folA | fau | Folate cycle | One-carbon | 1.0000000000006 | 0.3811258549700434 | 0.5864520805112566 | False | bifurcation only | True | 58.0 |
| 384 | folB | fau | Biosynthesis | One-carbon | 1.000000000000158 | 0.381125854969875 | 0.102078393846666 | False | bifurcation only | True | 59.0 |
| 399 | glyA | fau | One-carbon | One-carbon | 1.000000000000002 | 0.3811258549698155 | 1.6427703444719313 | False | bifurcation only | True | 60.0 |
| 406 | metH | fau | Methyl | One-carbon | 1.000000000000001 | 0.3811258549698152 | 2.352429622056764 | False | bifurcation only | True | 61.0 |
| 418 | folM | fau | Folate cycle | One-carbon | 0.9999999999999992 | 0.3811258549698145 | 2.2991076621892725 | False | bifurcation only | True | 62.0 |
| 423 | folD_MTHFC | fau | One-carbon | One-carbon | 0.9999999999999988 | 0.3811258549698142 | 1.3832734169958778 | False | bifurcation only | True | 63.0 |
| 439 | folC_DHFS | fau | Biosynthesis | One-carbon | 0.9999999999999584 | 0.3811258549697989 | 1.6170552827146394 | False | bifurcation only | True | 64.0 |
| 444 | fhs | fau | One-carbon | One-carbon | 0.9999999999998695 | 0.3811258549697651 | 1.2747515908732103 | False | bifurcation only | True | 65.0 |
| 454 | metF | fau | Methyl | One-carbon | 0.9999999999997352 | 0.3811258549697138 | 5.519387202024339 | False | bifurcation only | True | 66.0 |
| 478 | folP | fau | Biosynthesis | One-carbon | 0.9999999999989015 | 0.3811258549693961 | 0.103466135263448 | False | bifurcation only | True | 67.0 |
| 494 | folM | sam_methyltransferase | Folate cycle | Methyl | 0.99999987734331 | 0.3811258082221789 | 2.316136537961186 | False | bifurcation only | True | 68.0 |
| 495 | folM | sah_hydrolase | Folate cycle | Methyl | 0.9999998652735872 | 0.3811258036220955 | 2.3171382012754043 | False | bifurcation only | True | 69.0 |
| 496 | folM | metK | Folate cycle | Methyl | 0.9999998020988294 | 0.3811257795445619 | 2.320154232379765 | False | bifurcation only | True | 70.0 |
| 498 | folM | folX | Folate cycle | Biosynthesis | 0.9999957528818636 | 0.3811242362832839 | 2.619434477432199 | False | bifurcation only | True | 71.0 |
| 511 | folA | sam_methyltransferase | Folate cycle | Methyl | 0.9998224958906798 | 0.3810582035643894 | 0.5907957742796889 | False | bifurcation only | True | 72.0 |
| 512 | folC_DHFS | sam_methyltransferase | Biosynthesis | Methyl | 0.9998196881106508 | 0.3810571334468253 | 1.6290323788630827 | False | bifurcation only | True | 73.0 |
| 528 | folC_DHFS | sah_hydrolase | Biosynthesis | Methyl | 0.9993864585018946 | 0.3808920184417899 | 1.629736889130434 | False | bifurcation only | True | 74.0 |
| 529 | folA | sah_hydrolase | Folate cycle | Methyl | 0.9993664108095944 | 0.3808843777479219 | 0.591051276683624 | False | bifurcation only | True | 75.0 |
| 533 | metH | metK | Methyl | Methyl | 0.999300792549399 | 0.3808593689324032 | 2.37396431396052 | False | bifurcation only | True | 76.0 |
| 539 | metH | sam_methyltransferase | Methyl | Methyl | 0.999040504587576 | 0.3807601664604151 | 2.3698534393294306 | False | bifurcation only | True | 77.0 |
| 540 | metH | sah_hydrolase | Methyl | Methyl | 0.9990356070480944 | 0.3807582998814929 | 2.370878333678854 | False | bifurcation only | True | 78.0 |
| 549 | folX | metH | Biosynthesis | Methyl | 0.9977282183671397 | 0.3802600202526862 | 2.6801856037836127 | False | bifurcation only | True | 79.0 |
| 553 | folC_DHFS | metK | Biosynthesis | Methyl | 0.9957452824418056 | 0.3795042721027928 | 1.631858185627482 | False | bifurcation only | True | 80.0 |
| 558 | folA | metK | Folate cycle | Methyl | 0.9955304272200748 | 0.379422385222716 | 0.5918206002542977 | False | bifurcation only | True | 81.0 |
| 560 | thyA | sam_methyltransferase | One-carbon | Methyl | 0.9944815606952272 | 0.3790226350716841 | 0.7570284819453689 | False | bifurcation only | True | 82.0 |
| 562 | thyA | sah_hydrolase | One-carbon | Methyl | 0.9941641298871808 | 0.3789016539835738 | 0.7573558752772189 | False | bifurcation only | True | 83.0 |
| 565 | metF | metK | Methyl | Methyl | 0.9935333124928728 | 0.3786612331648383 | 5.569912965591798 | False | bifurcation only | True | 84.0 |
| 566 | folC_FPGS | sam_methyltransferase | Polyglutamation | Methyl | 0.9933643585282812 | 0.3785968404406328 | 16.569578902755154 | False | bifurcation only | True | 85.0 |
| 567 | thyA | metK | One-carbon | Methyl | 0.9932405900574314 | 0.3785496690763618 | 0.7583416640729193 | False | bifurcation only | True | 86.0 |
| 568 | folC_FPGS | sah_hydrolase | Polyglutamation | Methyl | 0.993040240884666 | 0.3784733108265992 | 16.576744775339478 | False | bifurcation only | True | 87.0 |
| 569 | folB | metK | Biosynthesis | Methyl | 0.9926993746175288 | 0.3783433978791061 | 0.1030128433795691 | False | bifurcation only | True | 88.0 |
| 570 | folP | sam_methyltransferase | Biosynthesis | Methyl | 0.9925693540723578 | 0.3782938436876641 | 0.1042324812649761 | False | bifurcation only | True | 89.0 |
| 571 | folB | sam_methyltransferase | Biosynthesis | Methyl | 0.9924528098107324 | 0.3782494256563103 | 0.102834461218542 | False | bifurcation only | True | 90.0 |
| 572 | folC_FPGS | metK | Polyglutamation | Methyl | 0.9924149281842018 | 0.3782349879890113 | 16.598321381268274 | False | bifurcation only | True | 91.0 |
| 573 | metF | sam_methyltransferase | Methyl | Methyl | 0.9923545400889656 | 0.3782119725245844 | 5.560267827384379 | False | bifurcation only | True | 92.0 |
| 575 | folP | sah_hydrolase | Biosynthesis | Methyl | 0.9922232633854634 | 0.3781619395787244 | 0.104277558854713 | False | bifurcation only | True | 93.0 |
| 576 | folB | sah_hydrolase | Biosynthesis | Methyl | 0.9922231026960712 | 0.3781618783358425 | 0.1028789342042886 | False | bifurcation only | True | 94.0 |
| 578 | metF | sah_hydrolase | Methyl | Methyl | 0.992219057960036 | 0.3781603367823629 | 5.562672485403731 | False | bifurcation only | True | 95.0 |
| 579 | folP | metK | Biosynthesis | Methyl | 0.9916667745167774 | 0.3779498472828653 | 0.1044132885064099 | False | bifurcation only | True | 96.0 |
| 582 | folK | metF | Biosynthesis | Methyl | 2.576739579568947 | 0.3742892480812954 | 0.8638930818782211 | False | bifurcation only | True | 97.0 |
| 583 | folX | folD_MTHFD | Biosynthesis | One-carbon | 0.9806140081943782 | 0.3737373522684593 | 1.8235512799061844 | False | bifurcation only | True | 98.0 |
| 584 | folP | metF | Biosynthesis | Methyl | 2.550017910623604 | 0.3704077408244764 | 0.2483824410080731 | False | bifurcation only | True | 99.0 |
| 585 | folB | metF | Biosynthesis | Methyl | 2.547126881263131 | 0.3699877988116737 | 0.2450510070107497 | False | bifurcation only | True | 100.0 |

Table S9a. Top-100 druggable node pairs ranked by pairwise synergy score in the folate-specific subset. Only nodes in the committed folate arm and the one-carbon cycle are considered, yielding 190 unordered pairs. For each pair, the table reports the full-network rank, the two node identities, their pathway modules, the raw and corrected synergy scores, the additive dTMP reference, a Boolean flag for whether the pair falls below the additive floor, the correction type applied (floor + bifurcation, bifurcation only, or none), a Boolean indicating folate specificity, and the folate-specific rank. The top-ranked druggable pair is folP (DHPS) + folA (DHFR) at rank 2, which is the pair validated experimentally.

Supplementary Table S10

| Condition | mu_max_mean | mu_max_SD | OD_16h_mean | OD_16h_SD | lag_mean_h | lag_SD_h | AUC_mean | AUC_SD |
| --- | --- | --- | --- | --- | --- | --- | --- | --- |
| SMX | 0.13601 | 0.137035 | 0.280845 | 0.0968221 | 8.49758 | 2.51561 | 2.42591 | 0.657692 |
| TMP | 0.0804248 | 0.113495 | 0.243618 | 0.116557 | 7.94254 | 3.99822 | 2.15785 | 0.620364 |
| TMP+SMX arm 1 | 0.00829015 | 0.00878348 | 0.0903636 | 0.00332424 |  |  | 1.40277 | 0.0180876 |
| TMP+SMX arm 2 | 0.0929942 | 0.181564 | 0.124636 | 0.0749198 | 5.07316 | 5.47128 | 1.71781 | 0.749053 |
| WT | 0.233766 | 0.153107 | 0.3225 | 0.031482 | 6.60521 | 1.54528 | 2.75039 | 0.467787 |

Table S10. Per-condition summary statistics for the single-agent and combination treatments. For each condition (SMX, TMP, TMP+SMX Arm 1, TMP+SMX Arm 2, WT), the table reports the mean and standard deviation of μ_max (h⁻¹), OD₆₀₀ at 16 h, lag time (h), and AUC (0–16 h), aggregated across replicate wells. The progression from single agents to combination arms shows a marked reduction in growth rate and total biomass, consistent with synergy. Blank cells for Arms 1 and 2 indicate that lag time could not be reliably estimated because growth was too severely inhibited.

Supplementary Table S11

| Rank_Full | Node_A | Node_B | Module_A | Module_B | Raw_Synergy | Corrected_Synergy | Additive_dTMP | Below_Floor | Correction_Type | Folate_Specific | Rank_Folate |
| --- | --- | --- | --- | --- | --- | --- | --- | --- | --- | --- | --- |
| 1 | folK_pp | gcv | Biosynthesis | One-carbon | 2.61157 | 2.06845 | 0.0451141 | True | floor+bifurcation | False |  |
| 2 | folK | gcv | Biosynthesis | One-carbon | 2.68602 | 2.02402 | 0.0429216 | True | floor+bifurcation | True | 1.0 |
| 3 | folP | folA | Biosynthesis | Folate cycle | 3.90424 | 1.80896 | 0.0263914 | True | floor+bifurcation | True | 2.0 |
| 4 | folE | folX | Biosynthesis | Biosynthesis | 1.79625 | 1.79625 | 0.0648961 | False | none | False |  |
| 5 | folB | folA | Biosynthesis | Folate cycle | 3.90441 | 1.78477 | 0.0260374 | True | floor+bifurcation | True | 3.0 |
| 6 | nudB | folA | Biosynthesis | Folate cycle | 3.90615 | 1.45358 | 0.0211963 | True | floor+bifurcation | False |  |
| 7 | folB | fhs | Biosynthesis | One-carbon | 1.43361 | 1.42447 | 0.0565967 | True | floor+bifurcation | True | 4.0 |
| 8 | folC_FPGS | gcv | Polyglutamation | One-carbon | 3.73692 | 1.42424 | 1.96176 | False | bifurcation only | True | 5.0 |
| 9 | folB | folK_pp | Biosynthesis | Biosynthesis | 4.40993 | 1.30017 | 0.0167934 | True | floor+bifurcation | False |  |
| 10 | folB | folK | Biosynthesis | Biosynthesis | 4.62653 | 1.29774 | 0.0159773 | True | floor+bifurcation | True | 6.0 |
| 11 | folK_pp | folP | Biosynthesis | Biosynthesis | 4.33041 | 1.29409 | 0.0170217 | True | floor+bifurcation | False |  |
| 12 | purU | gcv | Purine | One-carbon | 1.28754 | 1.28754 | 0.350172 | False | none | False |  |
| 13 | folK | folP | Biosynthesis | Biosynthesis | 4.5003 | 1.2795 | 0.0161945 | True | floor+bifurcation | True | 7.0 |
| 14 | nudB | folD_MTHFC | Biosynthesis | One-carbon | 1.36162 | 1.19514 | 0.049996 | True | floor+bifurcation | False |  |
| 15 | nudB | fhs | Biosynthesis | One-carbon | 1.4429 | 1.16713 | 0.0460737 | True | floor+bifurcation | False |  |
| 16 | folA | gcv | Folate cycle | One-carbon | 2.90305 | 1.10643 | 0.0699473 | False | bifurcation only | True | 8.0 |
| 17 | nudB | folK_pp | Biosynthesis | Biosynthesis | 4.47415 | 1.07384 | 0.013671 | True | floor+bifurcation | False |  |
| 18 | nudB | folK | Biosynthesis | Biosynthesis | 4.69584 | 1.07228 | 0.0130066 | True | floor+bifurcation | False |  |
| 19 | metF | gcv | Methyl | One-carbon | 2.67733 | 1.0204 | 0.658308 | False | bifurcation only | True | 9.0 |
| 20 | purH | purU | Purine | Purine | 1.0066 | 1.0066 | 2.95786 | False | none | False |  |
| 21 | folE | queD | Biosynthesis | PreQ1 | 1.00531 | 1.00531 | 0.0570931 | False | none | False |  |
| 22 | purH | gcv | Purine | One-carbon | 1.00259 | 1.00259 | 0.276275 | False | none | False |  |
| 23 | purN | purU | Purine | Purine | 1.0005 | 1.0005 | 2.93752 | False | none | False |  |
| 24 | folate_uptake | folate_reductase | Salvage | Salvage | 1.00021 | 1.00021 | 2.2979 | False | none | False |  |
| 25 | purN | gcv | Purine | One-carbon | 1.00019 | 1.00019 | 0.274374 | False | none | False |  |
| 26 | folate_uptake | gcv | Salvage | One-carbon | 1.00011 | 1.00011 | 0.274139 | False | none | False |  |
| 27 | folate_reductase | gcv | Salvage | One-carbon | 1.00008 | 1.00008 | 0.27416 | False | none | False |  |
| 28 | folate_uptake | folX | Salvage | Biosynthesis | 1.00003 | 1.00003 | 2.61867 | False | none | False |  |
| 29 | folate_reductase | folX | Salvage | Biosynthesis | 1.00002 | 1.00002 | 2.61887 | False | none | False |  |
| 30 | purN | purH | Purine | Purine | 1.00001 | 1.00001 | 2.31761 | False | none | False |  |
| 31 | folate_uptake | purU | Salvage | Purine | 1.00001 | 1.00001 | 2.935 | False | none | False |  |
| 32 | folate_reductase | purU | Salvage | Purine | 1.00001 | 1.00001 | 2.93522 | False | none | False |  |
| 33 | folate_uptake | metK | Salvage | Methyl | 1.0 | 1.0 | 2.31948 | False | none | False |  |
| 34 | folate_uptake | sah_hydrolase | Salvage | Methyl | 1.0 | 1.0 | 2.31646 | False | none | False |  |
| 35 | folate_reductase | metK | Salvage | Methyl | 1.0 | 1.0 | 2.31966 | False | none | False |  |
| 36 | folate_uptake | sam_methyltransferase | Salvage | Methyl | 1.0 | 1.0 | 2.31546 | False | none | False |  |
| 37 | folate_reductase | sah_hydrolase | Salvage | Methyl | 1.0 | 1.0 | 2.31664 | False | none | False |  |
| 38 | folate_reductase | sam_methyltransferase | Salvage | Methyl | 1.0 | 1.0 | 2.31564 | False | none | False |  |
| 39 | folate_uptake | queD | Salvage | PreQ1 | 1.0 | 1.0 | 2.30381 | False | none | False |  |
| 40 | folate_reductase | queD | Salvage | PreQ1 | 1.0 | 1.0 | 2.30398 | False | none | False |  |
| 41 | folate_uptake | purH | Salvage | Purine | 1.0 | 1.0 | 2.31562 | False | none | False |  |
| 42 | folate_reductase | purH | Salvage | Purine | 1.0 | 1.0 | 2.3158 | False | none | False |  |
| 43 | folate_uptake | purN | Salvage | Purine | 1.0 | 1.0 | 2.29969 | False | none | False |  |
| 44 | folate_reductase | purN | Salvage | Purine | 1.0 | 1.0 | 2.29987 | False | none | False |  |
| 45 | folM_monapterin | moeA | Biosynthesis | MoCo | 1.0 | 1.0 | 2.29915 | False | none | False |  |
| 46 | folM_monapterin | mog | Biosynthesis | MoCo | 1.0 | 1.0 | 2.29915 | False | none | False |  |
| 47 | folate_uptake | folM_monapterin | Salvage | Biosynthesis | 1.0 | 1.0 | 2.29844 | False | none | False |  |
| 48 | folM_monapterin | metK | Biosynthesis | Methyl | 1.0 | 1.0 | 2.3202 | False | none | False |  |
| 49 | folM_monapterin | sah_hydrolase | Biosynthesis | Methyl | 1.0 | 1.0 | 2.31719 | False | none | False |  |
| 50 | gcv | moeA | One-carbon | MoCo | 1.0 | 1.0 | 0.274225 | False | none | False |  |
| 51 | folM_monapterin | purU | Biosynthesis | Purine | 1.0 | 1.0 | 2.93592 | False | none | False |  |
| 52 | moaE | mog | MoCo | MoCo | 1.0 | 1.0 | 2.29915 | False | none | False |  |
| 53 | gcv | mobA | One-carbon | MoCo | 1.0 | 1.0 | 0.274225 | False | none | False |  |
| 54 | gcv | fau | One-carbon | One-carbon | 1.0 | 1.0 | 0.274225 | False | none | True | 10.0 |
| 55 | fau | mobA | One-carbon | MoCo | 1.0 | 1.0 | 2.29915 | False | none | False |  |
| 56 | gcv | queE | One-carbon | PreQ1 | 1.0 | 1.0 | 0.274225 | False | none | False |  |
| 57 | folate_reductase | folM_monapterin | Salvage | Biosynthesis | 1.0 | 1.0 | 2.29861 | False | none | False |  |
| 58 | gcv | mog | One-carbon | MoCo | 1.0 | 1.0 | 0.274225 | False | none | False |  |
| 59 | folM_monapterin | queD | Biosynthesis | PreQ1 | 1.0 | 1.0 | 2.30453 | False | none | False |  |
| 60 | gcv | moaE | One-carbon | MoCo | 1.0 | 1.0 | 0.274225 | False | none | False |  |
| 61 | folX | mobA | Biosynthesis | MoCo | 1.0 | 1.0 | 2.61949 | False | none | False |  |
| 62 | folate_reductase | queF | Salvage | PreQ1 | 1.0 | 1.0 | 2.29861 | False | none | False |  |
| 63 | folate_reductase | moeA | Salvage | MoCo | 1.0 | 1.0 | 2.29861 | False | none | False |  |
| 64 | folate_reductase | moaA | Salvage | MoCo | 1.0 | 1.0 | 2.29861 | False | none | False |  |
| 65 | mog | moeA | MoCo | MoCo | 1.0 | 1.0 | 2.29915 | False | none | False |  |
| 66 | folate_uptake | mog | Salvage | MoCo | 1.0 | 1.0 | 2.29844 | False | none | False |  |
| 67 | mog | queC | MoCo | PreQ1 | 1.0 | 1.0 | 2.29915 | False | none | False |  |
| 68 | purH | queE | Purine | PreQ1 | 1.0 | 1.0 | 2.31634 | False | none | False |  |
| 69 | folate_reductase | queE | Salvage | PreQ1 | 1.0 | 1.0 | 2.29861 | False | none | False |  |
| 70 | purU | mobA | Purine | MoCo | 1.0 | 1.0 | 2.93592 | False | none | False |  |
| 71 | purN | moeA | Purine | MoCo | 1.0 | 1.0 | 2.30041 | False | none | False |  |
| 72 | purN | moaA | Purine | MoCo | 1.0 | 1.0 | 2.30041 | False | none | False |  |
| 73 | queD | queC | PreQ1 | PreQ1 | 1.0 | 1.0 | 2.30453 | False | none | False |  |
| 74 | purN | queC | Purine | PreQ1 | 1.0 | 1.0 | 2.30041 | False | none | False |  |
| 75 | sah_hydrolase | moeA | Methyl | MoCo | 1.0 | 1.0 | 2.31719 | False | none | False |  |
| 76 | metK | moeA | Methyl | MoCo | 1.0 | 1.0 | 2.3202 | False | none | False |  |
| 77 | purN | fau | Purine | One-carbon | 1.0 | 1.0 | 2.30041 | False | none | False |  |
| 78 | moeA | queC | MoCo | PreQ1 | 1.0 | 1.0 | 2.29915 | False | none | False |  |
| 79 | folX | queC | Biosynthesis | PreQ1 | 1.0 | 1.0 | 2.61949 | False | none | False |  |
| 80 | folM_monapterin | purN | Biosynthesis | Purine | 1.0 | 1.0 | 2.30041 | False | none | False |  |
| 81 | metK | moaA | Methyl | MoCo | 1.0 | 1.0 | 2.3202 | False | none | False |  |
| 82 | folX | mog | Biosynthesis | MoCo | 1.0 | 1.0 | 2.61949 | False | none | False |  |
| 83 | metK | mog | Methyl | MoCo | 1.0 | 1.0 | 2.3202 | False | none | False |  |
| 84 | purH | moeA | Purine | MoCo | 1.0 | 1.0 | 2.31634 | False | none | False |  |
| 85 | purN | mog | Purine | MoCo | 1.0 | 1.0 | 2.30041 | False | none | False |  |
| 86 | sah_hydrolase | queE | Methyl | PreQ1 | 1.0 | 1.0 | 2.31719 | False | none | False |  |
| 87 | moeA | queE | MoCo | PreQ1 | 1.0 | 1.0 | 2.29915 | False | none | False |  |
| 88 | folX | moeA | Biosynthesis | MoCo | 1.0 | 1.0 | 2.61949 | False | none | False |  |
| 89 | purN | mobA | Purine | MoCo | 1.0 | 1.0 | 2.30041 | False | none | False |  |
| 90 | mog | queE | MoCo | PreQ1 | 1.0 | 1.0 | 2.29915 | False | none | False |  |
| 91 | folM_monapterin | moaE | Biosynthesis | MoCo | 1.0 | 1.0 | 2.29915 | False | none | False |  |
| 92 | folM_monapterin | mobA | Biosynthesis | MoCo | 1.0 | 1.0 | 2.29915 | False | none | False |  |
| 93 | fau | sah_hydrolase | One-carbon | Methyl | 1.0 | 1.0 | 2.31719 | False | none | True | 11.0 |
| 94 | folate_uptake | mobA | Salvage | MoCo | 1.0 | 1.0 | 2.29844 | False | none | False |  |
| 95 | sam_methyltransferase | mobA | Methyl | MoCo | 1.0 | 1.0 | 2.31618 | False | none | False |  |
| 96 | folX | queE | Biosynthesis | PreQ1 | 1.0 | 1.0 | 2.61949 | False | none | False |  |
| 97 | moeA | queD | MoCo | PreQ1 | 1.0 | 1.0 | 2.30453 | False | none | False |  |
| 98 | moaA | moeA | MoCo | MoCo | 1.0 | 1.0 | 2.29915 | False | none | False |  |
| 99 | fau | mog | One-carbon | MoCo | 1.0 | 1.0 | 2.29915 | False | none | False |  |
| 100 | fau | sam_methyltransferase | One-carbon | Methyl | 1.0 | 1.0 | 2.31618 | False | none | True | 12.0 |
| 101 | sam_methyltransferase | moeA | Methyl | MoCo | 1.0 | 1.0 | 2.31618 | False | none | False |  |
| 102 | purN | queE | Purine | PreQ1 | 1.0 | 1.0 | 2.30041 | False | none | False |  |
| 103 | purH | queC | Purine | PreQ1 | 1.0 | 1.0 | 2.31634 | False | none | False |  |
| 104 | purU | moaA | Purine | MoCo | 1.0 | 1.0 | 2.93592 | False | none | False |  |
| 105 | folX | moaA | Biosynthesis | MoCo | 1.0 | 1.0 | 2.61949 | False | none | False |  |
| 106 | purH | moaA | Purine | MoCo | 1.0 | 1.0 | 2.31634 | False | none | False |  |
| 107 | moaA | queD | MoCo | PreQ1 | 1.0 | 1.0 | 2.30453 | False | none | False |  |
| 108 | moaA | mog | MoCo | MoCo | 1.0 | 1.0 | 2.29915 | False | none | False |  |
| 109 | folate_uptake | queC | Salvage | PreQ1 | 1.0 | 1.0 | 2.29844 | False | none | False |  |
| 110 | folate_uptake | moeA | Salvage | MoCo | 1.0 | 1.0 | 2.29844 | False | none | False |  |
| 111 | purH | mog | Purine | MoCo | 1.0 | 1.0 | 2.31634 | False | none | False |  |
| 112 | folX | folM_monapterin | Biosynthesis | Biosynthesis | 1.0 | 1.0 | 2.61949 | False | none | False |  |
| 113 | mog | queD | MoCo | PreQ1 | 1.0 | 1.0 | 2.30453 | False | none | False |  |
| 114 | folM_monapterin | fau | Biosynthesis | One-carbon | 1.0 | 1.0 | 2.29915 | False | none | False |  |
| 115 | fau | queD | One-carbon | PreQ1 | 1.0 | 1.0 | 2.30453 | False | none | False |  |
| 116 | queD | queE | PreQ1 | PreQ1 | 1.0 | 1.0 | 2.30453 | False | none | False |  |
| 117 | queD | queF | PreQ1 | PreQ1 | 1.0 | 1.0 | 2.30453 | False | none | False |  |
| 118 | moaA | queE | MoCo | PreQ1 | 1.0 | 1.0 | 2.29915 | False | none | False |  |
| 119 | fau | metK | One-carbon | Methyl | 1.0 | 1.0 | 2.3202 | False | none | True | 13.0 |
| 120 | folate_uptake | moaA | Salvage | MoCo | 1.0 | 1.0 | 2.29844 | False | none | False |  |
| 121 | metK | queC | Methyl | PreQ1 | 1.0 | 1.0 | 2.3202 | False | none | False |  |
| 122 | fau | moeA | One-carbon | MoCo | 1.0 | 1.0 | 2.29915 | False | none | False |  |
| 123 | fau | queC | One-carbon | PreQ1 | 1.0 | 1.0 | 2.29915 | False | none | False |  |
| 124 | fau | queE | One-carbon | PreQ1 | 1.0 | 1.0 | 2.29915 | False | none | False |  |
| 125 | purU | mog | Purine | MoCo | 1.0 | 1.0 | 2.93592 | False | none | False |  |
| 126 | metK | queE | Methyl | PreQ1 | 1.0 | 1.0 | 2.3202 | False | none | False |  |
| 127 | sah_hydrolase | mog | Methyl | MoCo | 1.0 | 1.0 | 2.31719 | False | none | False |  |
| 128 | purU | queC | Purine | PreQ1 | 1.0 | 1.0 | 2.93592 | False | none | False |  |
| 129 | folM_monapterin | queE | Biosynthesis | PreQ1 | 1.0 | 1.0 | 2.29915 | False | none | False |  |
| 130 | metK | mobA | Methyl | MoCo | 1.0 | 1.0 | 2.3202 | False | none | False |  |
| 131 | purU | moeA | Purine | MoCo | 1.0 | 1.0 | 2.93592 | False | none | False |  |
| 132 | folX | fau | Biosynthesis | One-carbon | 1.0 | 1.0 | 2.61949 | False | none | True | 14.0 |
| 133 | moaA | mobA | MoCo | MoCo | 1.0 | 1.0 | 2.29915 | False | none | False |  |
| 134 | sah_hydrolase | queC | Methyl | PreQ1 | 1.0 | 1.0 | 2.31719 | False | none | False |  |
| 135 | fau | moaA | One-carbon | MoCo | 1.0 | 1.0 | 2.29915 | False | none | False |  |
| 136 | metK | moaE | Methyl | MoCo | 1.0 | 1.0 | 2.3202 | False | none | False |  |
| 137 | purN | queF | Purine | PreQ1 | 1.0 | 1.0 | 2.30041 | False | none | False |  |
| 138 | sam_methyltransferase | queC | Methyl | PreQ1 | 1.0 | 1.0 | 2.31618 | False | none | False |  |
| 139 | sah_hydrolase | moaA | Methyl | MoCo | 1.0 | 1.0 | 2.31719 | False | none | False |  |
| 140 | mobA | queE | MoCo | PreQ1 | 1.0 | 1.0 | 2.29915 | False | none | False |  |
| 141 | queE | queC | PreQ1 | PreQ1 | 1.0 | 1.0 | 2.29915 | False | none | False |  |
| 142 | mobA | queD | MoCo | PreQ1 | 1.0 | 1.0 | 2.30453 | False | none | False |  |
| 143 | purU | fau | Purine | One-carbon | 1.0 | 1.0 | 2.93592 | False | none | False |  |
| 144 | folate_uptake | queE | Salvage | PreQ1 | 1.0 | 1.0 | 2.29844 | False | none | False |  |
| 145 | moaA | queC | MoCo | PreQ1 | 1.0 | 1.0 | 2.29915 | False | none | False |  |
| 146 | sam_methyltransferase | queE | Methyl | PreQ1 | 1.0 | 1.0 | 2.31618 | False | none | False |  |
| 147 | purU | moaE | Purine | MoCo | 1.0 | 1.0 | 2.93592 | False | none | False |  |
| 148 | sam_methyltransferase | moaA | Methyl | MoCo | 1.0 | 1.0 | 2.31618 | False | none | False |  |
| 149 | purH | fau | Purine | One-carbon | 1.0 | 1.0 | 2.31634 | False | none | False |  |
| 150 | sam_methyltransferase | mog | Methyl | MoCo | 1.0 | 1.0 | 2.31618 | False | none | False |  |
| 151 | moeA | queF | MoCo | PreQ1 | 1.0 | 1.0 | 2.29915 | False | none | False |  |
| 152 | sam_methyltransferase | queF | Methyl | PreQ1 | 1.0 | 1.0 | 2.31618 | False | none | False |  |
| 153 | mog | queF | MoCo | PreQ1 | 1.0 | 1.0 | 2.29915 | False | none | False |  |
| 154 | folate_reductase | queC | Salvage | PreQ1 | 1.0 | 1.0 | 2.29861 | False | none | False |  |
| 155 | metK | queF | Methyl | PreQ1 | 1.0 | 1.0 | 2.3202 | False | none | False |  |
| 156 | moaE | queD | MoCo | PreQ1 | 1.0 | 1.0 | 2.30453 | False | none | False |  |
| 157 | purN | moaE | Purine | MoCo | 1.0 | 1.0 | 2.30041 | False | none | False |  |
| 158 | queE | queF | PreQ1 | PreQ1 | 1.0 | 1.0 | 2.29915 | False | none | False |  |
| 159 | moaE | moeA | MoCo | MoCo | 1.0 | 1.0 | 2.29915 | False | none | False |  |
| 160 | fau | queF | One-carbon | PreQ1 | 1.0 | 1.0 | 2.29915 | False | none | False |  |
| 161 | moaE | queC | MoCo | PreQ1 | 1.0 | 1.0 | 2.29915 | False | none | False |  |
| 162 | folX | queF | Biosynthesis | PreQ1 | 1.0 | 1.0 | 2.61949 | False | none | False |  |
| 163 | moaE | queE | MoCo | PreQ1 | 1.0 | 1.0 | 2.29915 | False | none | False |  |
| 164 | folate_uptake | moaE | Salvage | MoCo | 1.0 | 1.0 | 2.29844 | False | none | False |  |
| 165 | purU | queE | Purine | PreQ1 | 1.0 | 1.0 | 2.93592 | False | none | False |  |
| 166 | purH | moaE | Purine | MoCo | 1.0 | 1.0 | 2.31634 | False | none | False |  |
| 167 | sah_hydrolase | moaE | Methyl | MoCo | 1.0 | 1.0 | 2.31719 | False | none | False |  |
| 168 | folate_uptake | queF | Salvage | PreQ1 | 1.0 | 1.0 | 2.29844 | False | none | False |  |
| 169 | fau | moaE | One-carbon | MoCo | 1.0 | 1.0 | 2.29915 | False | none | False |  |
| 170 | moaA | queF | MoCo | PreQ1 | 1.0 | 1.0 | 2.29915 | False | none | False |  |
| 171 | sah_hydrolase | queF | Methyl | PreQ1 | 1.0 | 1.0 | 2.31719 | False | none | False |  |
| 172 | sam_methyltransferase | moaE | Methyl | MoCo | 1.0 | 1.0 | 2.31618 | False | none | False |  |
| 173 | purH | queF | Purine | PreQ1 | 1.0 | 1.0 | 2.31634 | False | none | False |  |
| 174 | purU | queF | Purine | PreQ1 | 1.0 | 1.0 | 2.93592 | False | none | False |  |
| 175 | queC | queF | PreQ1 | PreQ1 | 1.0 | 1.0 | 2.29915 | False | none | False |  |
| 176 | folM_monapterin | sam_methyltransferase | Biosynthesis | Methyl | 1.0 | 1.0 | 2.31618 | False | none | False |  |
| 177 | folM_monapterin | purH | Biosynthesis | Purine | 1.0 | 1.0 | 2.31634 | False | none | False |  |
| 178 | moaE | queF | MoCo | PreQ1 | 1.0 | 1.0 | 2.29915 | False | none | False |  |
| 179 | folX | moaE | Biosynthesis | MoCo | 1.0 | 1.0 | 2.61949 | False | none | False |  |
| 180 | moaA | moaE | MoCo | MoCo | 1.0 | 1.0 | 2.29915 | False | none | False |  |
| 181 | sah_hydrolase | mobA | Methyl | MoCo | 1.0 | 1.0 | 2.31719 | False | none | False |  |
| 182 | folM_monapterin | queF | Biosynthesis | PreQ1 | 1.0 | 1.0 | 2.29915 | False | none | False |  |
| 183 | folM_monapterin | moaA | Biosynthesis | MoCo | 1.0 | 1.0 | 2.29915 | False | none | False |  |
| 184 | folate_uptake | fau | Salvage | One-carbon | 1.0 | 1.0 | 2.29844 | False | none | False |  |
| 185 | folM_monapterin | gcv | Biosynthesis | One-carbon | 1.0 | 1.0 | 0.274225 | False | none | False |  |
| 186 | folate_reductase | mog | Salvage | MoCo | 1.0 | 1.0 | 2.29861 | False | none | False |  |
| 187 | moaE | mobA | MoCo | MoCo | 1.0 | 1.0 | 2.29915 | False | none | False |  |
| 188 | gcv | queF | One-carbon | PreQ1 | 1.0 | 1.0 | 0.274225 | False | none | False |  |
| 189 | folM_monapterin | queC | Biosynthesis | PreQ1 | 1.0 | 1.0 | 2.29915 | False | none | False |  |
| 190 | mobA | queC | MoCo | PreQ1 | 1.0 | 1.0 | 2.29915 | False | none | False |  |
| 191 | folate_reductase | fau | Salvage | One-carbon | 1.0 | 1.0 | 2.29861 | False | none | False |  |
| 192 | gcv | moaA | One-carbon | MoCo | 1.0 | 1.0 | 0.274225 | False | none | False |  |
| 193 | gcv | queC | One-carbon | PreQ1 | 1.0 | 1.0 | 0.274225 | False | none | False |  |
| 194 | moeA | mobA | MoCo | MoCo | 1.0 | 1.0 | 2.29915 | False | none | False |  |
| 195 | folE | queC | Biosynthesis | PreQ1 | 1.0 | 1.0 | 0.0569601 | True | floor+bifurcation | False |  |
| 196 | folate_reductase | moaE | Salvage | MoCo | 1.0 | 1.0 | 2.29861 | False | none | False |  |
| 197 | folate_reductase | mobA | Salvage | MoCo | 1.0 | 1.0 | 2.29861 | False | none | False |  |
| 198 | folE | moeA | Biosynthesis | MoCo | 1.0 | 1.0 | 0.0569601 | True | floor+bifurcation | False |  |
| 199 | folE | queE | Biosynthesis | PreQ1 | 1.0 | 1.0 | 0.0569601 | True | floor+bifurcation | False |  |
| 200 | folE | fau | Biosynthesis | One-carbon | 1.0 | 1.0 | 0.0569601 | False | none | False |  |

Table S11. Complete pairwise synergy ranking for all 703 unordered node pairs in the full network. Every pair of the 38 nodes in the folate, one-carbon, methyl, salvage, PreQ1, and MoCo modules was evaluated. For each pair, the table reports the full-network rank, node identities and pathway modules, raw and corrected synergy scores, the additive dTMP reference, a Boolean flag for whether the pair falls below the additive floor, the correction type applied, a Boolean indicating folate specificity, and the folate-specific rank (blank for non-folate pairs). The top-scoring pair in the full network is folK_pp + gcv; the top druggable folate-specific pair is folP + folA. The table has 703 rows, truncated here to the first 200. Full data are provided in the source CSV file.

*(Showing first 200 of 200 rows — full data in source CSV.)*

Supplementary Table S12

| factor | Vmax | WT_dTMP | KD_dTMP | KD_fold |
| --- | --- | --- | --- | --- |
| 0.01 | 0.119029 | 3.10682 | 2.89407 | 0.931522 |
| 0.0126896 | 0.151043 | 3.13183 | 2.92648 | 0.934431 |
| 0.0161026 | 0.191668 | 3.15436 | 2.95984 | 0.938333 |
| 0.0204336 | 0.243219 | 3.17447 | 2.99327 | 0.942918 |
| 0.0259294 | 0.308635 | 3.1923 | 3.02592 | 0.94788 |
| 0.0329034 | 0.391646 | 3.20801 | 3.05709 | 0.952955 |
| 0.0417532 | 0.496983 | 3.22179 | 3.08627 | 0.957936 |
| 0.0529832 | 0.630652 | 3.23381 | 3.11313 | 0.962681 |
| 0.0672336 | 0.800273 | 3.24426 | 3.13754 | 0.967102 |
| 0.0853168 | 1.01552 | 3.25331 | 3.15947 | 0.971156 |
| 0.108264 | 1.28865 | 3.26109 | 3.17901 | 0.97483 |
| 0.137382 | 1.63525 | 3.26777 | 3.19631 | 0.978133 |
| 0.174333 | 2.07506 | 3.27345 | 3.21153 | 0.981085 |
| 0.221222 | 2.63317 | 3.27827 | 3.22487 | 0.983711 |
| 0.280722 | 3.3414 | 3.28232 | 3.23649 | 0.986038 |
| 0.356225 | 4.2401 | 3.28571 | 3.24659 | 0.988092 |
| 0.452035 | 5.38052 | 3.28853 | 3.25531 | 0.989899 |
| 0.573615 | 6.82767 | 3.29085 | 3.26281 | 0.99148 |
| 0.727895 | 8.66405 | 3.29276 | 3.26924 | 0.992856 |
| 0.923671 | 10.9943 | 3.29431 | 3.2747 | 0.994046 |
| 1.1721 | 13.9514 | 3.29557 | 3.27932 | 0.995068 |
| 1.48735 | 17.7038 | 3.29659 | 3.28321 | 0.995939 |
| 1.88739 | 22.4654 | 3.29741 | 3.28645 | 0.996676 |
| 2.39503 | 28.5077 | 3.29806 | 3.28914 | 0.997293 |
| 3.0392 | 36.1752 | 3.29859 | 3.29135 | 0.997807 |
| 3.85662 | 45.9049 | 3.299 | 3.29317 | 0.998231 |
| 4.8939 | 58.2515 | 3.29933 | 3.29465 | 0.998579 |
| 6.21017 | 73.9189 | 3.2996 | 3.29584 | 0.998862 |
| 7.88046 | 93.8002 | 3.2998 | 3.29681 | 0.999092 |
| 10.0 | 119.029 | 3.29997 | 3.29758 | 0.999277 |

Table S12. DHFR V_max scan: steady-state dTMP flux as a function of DHFR maximum velocity. The V_max of dihydrofolate reductase (folA) was varied across 30 levels (factor 0.01 to 10.0 relative to wild type), and the resulting steady-state dTMP flux was recorded. For each V_max value, the table reports the WT_dTMP (μM) at the corresponding wild-type reference, the KD_dTMP (μM) under DHFR knockdown, and the ratio KD_fold = KD_dTMP / WT_dTMP. This scan demonstrates that dTMP production is remarkably insensitive to DHFR V_max above ~0.5× wild-type, but collapses sharply below that threshold, identifying DHFR as a single-path bottleneck in the network.

Supplementary Table S13

| scenario | w1 | w2 | w2b | w3 | F_total |
| --- | --- | --- | --- | --- | --- |
| Equal weights | 0.25 | 0.25 | 0.25 | 0.25 | 0.98 |
| Primary = ranking | 0.5 | 0.2 | 0.15 | 0.15 | 0.988 |
| Primary = synergy | 0.2 | 0.5 | 0.15 | 0.15 | 0.988 |
| Primary = category | 0.2 | 0.2 | 0.2 | 0.4 | 0.984 |
| Our weights | 0.35 | 0.35 | 0.15 | 0.15 | 0.988 |
| No F2b | 0.4 | 0.4 | 0.0 | 0.2 | 1.0 |
| No F3 | 0.4 | 0.4 | 0.2 | 0.0 | 0.984 |

Table S13. Per-node fitness sensitivity scenarios. Seven different weighting schemes were applied to the four fitness components (w1, w2, w2b, w3) used in the node-scoring function: equal weights, and six alternatives that emphasize ranking, synergy, category, or specific combinations. The resulting composite fitness score F_total is reported for each scenario. All scenarios yield F_total ≥ 0.98, demonstrating that the ranking is robust to the choice of weights.

Supplementary Table S14

| parameter | value |
| --- | --- |
| n | 10000.0 |
| Observed_F | 0.988 |
| Null_mean | 0.4975 |
| Null_95th_pct | 0.9385 |
| Observed_percentile | 99.18 |

Table S14. Null-model parameters for the multiplicative additive reference. A Monte-Carlo null model (n = 10,000 simulations) was constructed by randomly permuting the dose-response surface under the assumption of multiplicative additivity. The table reports the null distribution mean (0.4975), its 95th percentile (0.9385), and the observed composite fitness score (F = 0.9880). The observed score falls at the 99.18th percentile of the null distribution, indicating that the experimental synergy significantly exceeds what multiplicative additivity would predict.

Null model n=10000
Observed F = 0.9880
Null mean = 0.4975
Null 95th pct = 0.9385
Observed percentile = 99.18%

Supplementary Table S15

| gene | pathway | refined | betweenness | degree_in | degree_out | pagerank | log2FC |
| --- | --- | --- | --- | --- | --- | --- | --- |
| folE | biosynthesis | pure_linear | 0.00668037 | 2 | 2 | 0.0038281 | -2.61731 |
| nudB | biosynthesis | pure_linear | 0.00524152 | 1 | 2 | 0.00330842 | -2.35362 |
| phoA | biosynthesis | pure_linear | 0.00883864 | 1 | 2 | 0.00407954 | -2.02454 |
| folB | biosynthesis | pure_linear | 0.011665 | 1 | 1 | 0.00421883 | -0.238537 |
| folK | biosynthesis | pure_linear | 0.0150223 | 2 | 2 | 0.0055253 | -0.611115 |
| folP | biosynthesis | pure_linear | 0.0171292 | 2 | 2 | 0.00655699 | -1.88879 |
| folC | biosynthesis | pure_linear | 0.0197499 | 2 | 3 | 0.00896628 | -0.00339836 |
| folA | thf_cycle | loop_entry | 0.0200839 | 2 | 2 | 0.00905411 | -0.00809532 |
| glyA | thf_cycle | core_loop | 0.0110654 | 2 | 2 | 0.00895125 | 1.09428e-06 |
| thyA | thf_cycle | loop_exit | 0.0235868 | 2 | 2 | 0.0148265 | -2.28458 |
| metF | thf_cycle | core_loop | 0.00184138 | 2 | 1 | 0.0130956 | 4.44113 |
| metH | methyl | loop_exit | 0.0219253 | 1 | 2 | 0.0130186 | -1.46711e-10 |
| fhs | thf_cycle | core_loop | 0.0183967 | 2 | 1 | 0.0130523 | -0.0042574 |
| folX | peripheral | pure_linear | 0.00119904 | 1 | 1 | 0.00330842 | 0.128601 |
| gcv | methyl | loop_entry | 0.026122 | 2 | 3 | 0.0121842 | 0.00381529 |
| folD_MTHFC | thf_cycle | core_loop | 0.0146968 | 1 | 1 | 0.0139522 | 0.0214435 |
| folD_MTHFD | thf_cycle | core_loop | 0.0137718 | 1 | 1 | 0.0171131 | -2.35122e-05 |
| folC_FPGS | peripheral | pure_linear | 0.00971223 | 1 | 1 | 0.00687424 | 0.00292663 |
| purN | peripheral | loop_exit | 0.00154162 | 1 | 2 | 0.00452044 | -1.07808e-05 |
| purH | peripheral | pure_linear | 0.00154162 | 1 | 2 | 0.00452044 | -0.000153614 |
| purU | peripheral | pure_linear | 0.00138746 | 1 | 2 | 0.00313577 | -0.0040901 |
| fau | peripheral | pure_linear | 0.00138746 | 1 | 1 | 0.00348194 | -1.46711e-10 |
| metK | methyl | loop_exit | 0.00740836 | 2 | 3 | 0.00974484 | 0.00407145 |
| folM | peripheral | pure_linear | 0.0200839 | 2 | 2 | 0.00905411 | -0.000145465 |
| sah_hydrolase | methyl | loop_exit | 0.0148338 | 2 | 1 | 0.00724869 | -1.46678e-10 |
| sam_methyltransferase | methyl | other | 0.0129325 | 1 | 1 | 0.00850373 | -0.000455335 |

Table S15. Per-node topology metrics for the bipartite species-reaction graph. For each of the 26 nodes in the folate-specific network, the table reports the pathway assignment, topological refinement class, betweenness centrality, in-degree, out-degree, PageRank, and log₂ fold change in dTMP flux under knockdown. Betweenness was computed using the Brandes algorithm; PageRank used a damping factor of 0.85. The table reveals that folP (DHPS) has the highest betweenness in the network (0.0171) despite moderate degree, confirming its role as a topological hub.

Supplementary Table S16

| Rank_Full | Node_A | Node_B | Module_A | Module_B | Raw_Synergy | Corrected_Synergy | Additive_dTMP | Below_Floor | Correction_Type | Folate_Specific | Rank_Folate |
| --- | --- | --- | --- | --- | --- | --- | --- | --- | --- | --- | --- |
| 1 | folK_pp | gcv | Biosynthesis | One-carbon | 2.61157 | 2.06845 | 0.0451141 | True | floor+bifurcation | False |  |
| 2 | folK | gcv | Biosynthesis | One-carbon | 2.68602 | 2.02402 | 0.0429216 | True | floor+bifurcation | True | 1.0 |
| 3 | folP | folA | Biosynthesis | Folate cycle | 3.90424 | 1.80896 | 0.0263914 | True | floor+bifurcation | True | 2.0 |
| 4 | folE | folX | Biosynthesis | Biosynthesis | 1.79625 | 1.79625 | 0.0648961 | False | none | False |  |
| 5 | folB | folA | Biosynthesis | Folate cycle | 3.90441 | 1.78477 | 0.0260374 | True | floor+bifurcation | True | 3.0 |
| 6 | nudB | folA | Biosynthesis | Folate cycle | 3.90615 | 1.45358 | 0.0211963 | True | floor+bifurcation | False |  |
| 7 | folB | fhs | Biosynthesis | One-carbon | 1.43361 | 1.42447 | 0.0565967 | True | floor+bifurcation | True | 4.0 |
| 8 | folC_FPGS | gcv | Polyglutamation | One-carbon | 3.73692 | 1.42424 | 1.96176 | False | bifurcation only | True | 5.0 |
| 9 | folB | folK_pp | Biosynthesis | Biosynthesis | 4.40993 | 1.30017 | 0.0167934 | True | floor+bifurcation | False |  |
| 10 | folB | folK | Biosynthesis | Biosynthesis | 4.62653 | 1.29774 | 0.0159773 | True | floor+bifurcation | True | 6.0 |
| 11 | folK_pp | folP | Biosynthesis | Biosynthesis | 4.33041 | 1.29409 | 0.0170217 | True | floor+bifurcation | False |  |
| 12 | purU | gcv | Purine | One-carbon | 1.28754 | 1.28754 | 0.350172 | False | none | False |  |
| 13 | folK | folP | Biosynthesis | Biosynthesis | 4.5003 | 1.2795 | 0.0161945 | True | floor+bifurcation | True | 7.0 |
| 14 | nudB | folD_MTHFC | Biosynthesis | One-carbon | 1.36162 | 1.19514 | 0.049996 | True | floor+bifurcation | False |  |
| 15 | nudB | fhs | Biosynthesis | One-carbon | 1.4429 | 1.16713 | 0.0460737 | True | floor+bifurcation | False |  |
| 16 | folA | gcv | Folate cycle | One-carbon | 2.90305 | 1.10643 | 0.0699473 | False | bifurcation only | True | 8.0 |
| 17 | nudB | folK_pp | Biosynthesis | Biosynthesis | 4.47415 | 1.07384 | 0.013671 | True | floor+bifurcation | False |  |
| 18 | nudB | folK | Biosynthesis | Biosynthesis | 4.69584 | 1.07228 | 0.0130066 | True | floor+bifurcation | False |  |
| 19 | metF | gcv | Methyl | One-carbon | 2.67733 | 1.0204 | 0.658308 | False | bifurcation only | True | 9.0 |
| 20 | purH | purU | Purine | Purine | 1.0066 | 1.0066 | 2.95786 | False | none | False |  |
| 21 | folE | queD | Biosynthesis | PreQ1 | 1.00531 | 1.00531 | 0.0570931 | False | none | False |  |
| 22 | purH | gcv | Purine | One-carbon | 1.00259 | 1.00259 | 0.276275 | False | none | False |  |
| 23 | purN | purU | Purine | Purine | 1.0005 | 1.0005 | 2.93752 | False | none | False |  |
| 24 | folate_uptake | folate_reductase | Salvage | Salvage | 1.00021 | 1.00021 | 2.2979 | False | none | False |  |
| 25 | purN | gcv | Purine | One-carbon | 1.00019 | 1.00019 | 0.274374 | False | none | False |  |
| 26 | folate_uptake | gcv | Salvage | One-carbon | 1.00011 | 1.00011 | 0.274139 | False | none | False |  |
| 27 | folate_reductase | gcv | Salvage | One-carbon | 1.00008 | 1.00008 | 0.27416 | False | none | False |  |
| 28 | folate_uptake | folX | Salvage | Biosynthesis | 1.00003 | 1.00003 | 2.61867 | False | none | False |  |
| 29 | folate_reductase | folX | Salvage | Biosynthesis | 1.00002 | 1.00002 | 2.61887 | False | none | False |  |
| 30 | purN | purH | Purine | Purine | 1.00001 | 1.00001 | 2.31761 | False | none | False |  |
| 31 | folate_uptake | purU | Salvage | Purine | 1.00001 | 1.00001 | 2.935 | False | none | False |  |
| 32 | folate_reductase | purU | Salvage | Purine | 1.00001 | 1.00001 | 2.93522 | False | none | False |  |
| 33 | folate_uptake | metK | Salvage | Methyl | 1.0 | 1.0 | 2.31948 | False | none | False |  |
| 34 | folate_uptake | sah_hydrolase | Salvage | Methyl | 1.0 | 1.0 | 2.31646 | False | none | False |  |
| 35 | folate_reductase | metK | Salvage | Methyl | 1.0 | 1.0 | 2.31966 | False | none | False |  |
| 36 | folate_uptake | sam_methyltransferase | Salvage | Methyl | 1.0 | 1.0 | 2.31546 | False | none | False |  |
| 37 | folate_reductase | sah_hydrolase | Salvage | Methyl | 1.0 | 1.0 | 2.31664 | False | none | False |  |
| 38 | folate_reductase | sam_methyltransferase | Salvage | Methyl | 1.0 | 1.0 | 2.31564 | False | none | False |  |
| 39 | folate_uptake | queD | Salvage | PreQ1 | 1.0 | 1.0 | 2.30381 | False | none | False |  |
| 40 | folate_reductase | queD | Salvage | PreQ1 | 1.0 | 1.0 | 2.30398 | False | none | False |  |
| 41 | folate_uptake | purH | Salvage | Purine | 1.0 | 1.0 | 2.31562 | False | none | False |  |
| 42 | folate_reductase | purH | Salvage | Purine | 1.0 | 1.0 | 2.3158 | False | none | False |  |
| 43 | folate_uptake | purN | Salvage | Purine | 1.0 | 1.0 | 2.29969 | False | none | False |  |
| 44 | folate_reductase | purN | Salvage | Purine | 1.0 | 1.0 | 2.29987 | False | none | False |  |
| 45 | folM_monapterin | moeA | Biosynthesis | MoCo | 1.0 | 1.0 | 2.29915 | False | none | False |  |
| 46 | folM_monapterin | mog | Biosynthesis | MoCo | 1.0 | 1.0 | 2.29915 | False | none | False |  |
| 47 | folate_uptake | folM_monapterin | Salvage | Biosynthesis | 1.0 | 1.0 | 2.29844 | False | none | False |  |
| 48 | folM_monapterin | metK | Biosynthesis | Methyl | 1.0 | 1.0 | 2.3202 | False | none | False |  |
| 49 | folM_monapterin | sah_hydrolase | Biosynthesis | Methyl | 1.0 | 1.0 | 2.31719 | False | none | False |  |
| 50 | gcv | moeA | One-carbon | MoCo | 1.0 | 1.0 | 0.274225 | False | none | False |  |
| 51 | folM_monapterin | purU | Biosynthesis | Purine | 1.0 | 1.0 | 2.93592 | False | none | False |  |
| 52 | moaE | mog | MoCo | MoCo | 1.0 | 1.0 | 2.29915 | False | none | False |  |
| 53 | gcv | mobA | One-carbon | MoCo | 1.0 | 1.0 | 0.274225 | False | none | False |  |
| 54 | gcv | fau | One-carbon | One-carbon | 1.0 | 1.0 | 0.274225 | False | none | True | 10.0 |
| 55 | fau | mobA | One-carbon | MoCo | 1.0 | 1.0 | 2.29915 | False | none | False |  |
| 56 | gcv | queE | One-carbon | PreQ1 | 1.0 | 1.0 | 0.274225 | False | none | False |  |
| 57 | folate_reductase | folM_monapterin | Salvage | Biosynthesis | 1.0 | 1.0 | 2.29861 | False | none | False |  |
| 58 | gcv | mog | One-carbon | MoCo | 1.0 | 1.0 | 0.274225 | False | none | False |  |
| 59 | folM_monapterin | queD | Biosynthesis | PreQ1 | 1.0 | 1.0 | 2.30453 | False | none | False |  |
| 60 | gcv | moaE | One-carbon | MoCo | 1.0 | 1.0 | 0.274225 | False | none | False |  |
| 61 | folX | mobA | Biosynthesis | MoCo | 1.0 | 1.0 | 2.61949 | False | none | False |  |
| 62 | folate_reductase | queF | Salvage | PreQ1 | 1.0 | 1.0 | 2.29861 | False | none | False |  |
| 63 | folate_reductase | moeA | Salvage | MoCo | 1.0 | 1.0 | 2.29861 | False | none | False |  |
| 64 | folate_reductase | moaA | Salvage | MoCo | 1.0 | 1.0 | 2.29861 | False | none | False |  |
| 65 | mog | moeA | MoCo | MoCo | 1.0 | 1.0 | 2.29915 | False | none | False |  |
| 66 | folate_uptake | mog | Salvage | MoCo | 1.0 | 1.0 | 2.29844 | False | none | False |  |
| 67 | mog | queC | MoCo | PreQ1 | 1.0 | 1.0 | 2.29915 | False | none | False |  |
| 68 | purH | queE | Purine | PreQ1 | 1.0 | 1.0 | 2.31634 | False | none | False |  |
| 69 | folate_reductase | queE | Salvage | PreQ1 | 1.0 | 1.0 | 2.29861 | False | none | False |  |
| 70 | purU | mobA | Purine | MoCo | 1.0 | 1.0 | 2.93592 | False | none | False |  |
| 71 | purN | moeA | Purine | MoCo | 1.0 | 1.0 | 2.30041 | False | none | False |  |
| 72 | purN | moaA | Purine | MoCo | 1.0 | 1.0 | 2.30041 | False | none | False |  |
| 73 | queD | queC | PreQ1 | PreQ1 | 1.0 | 1.0 | 2.30453 | False | none | False |  |
| 74 | purN | queC | Purine | PreQ1 | 1.0 | 1.0 | 2.30041 | False | none | False |  |
| 75 | sah_hydrolase | moeA | Methyl | MoCo | 1.0 | 1.0 | 2.31719 | False | none | False |  |
| 76 | metK | moeA | Methyl | MoCo | 1.0 | 1.0 | 2.3202 | False | none | False |  |
| 77 | purN | fau | Purine | One-carbon | 1.0 | 1.0 | 2.30041 | False | none | False |  |
| 78 | moeA | queC | MoCo | PreQ1 | 1.0 | 1.0 | 2.29915 | False | none | False |  |
| 79 | folX | queC | Biosynthesis | PreQ1 | 1.0 | 1.0 | 2.61949 | False | none | False |  |
| 80 | folM_monapterin | purN | Biosynthesis | Purine | 1.0 | 1.0 | 2.30041 | False | none | False |  |
| 81 | metK | moaA | Methyl | MoCo | 1.0 | 1.0 | 2.3202 | False | none | False |  |
| 82 | folX | mog | Biosynthesis | MoCo | 1.0 | 1.0 | 2.61949 | False | none | False |  |
| 83 | metK | mog | Methyl | MoCo | 1.0 | 1.0 | 2.3202 | False | none | False |  |
| 84 | purH | moeA | Purine | MoCo | 1.0 | 1.0 | 2.31634 | False | none | False |  |
| 85 | purN | mog | Purine | MoCo | 1.0 | 1.0 | 2.30041 | False | none | False |  |
| 86 | sah_hydrolase | queE | Methyl | PreQ1 | 1.0 | 1.0 | 2.31719 | False | none | False |  |
| 87 | moeA | queE | MoCo | PreQ1 | 1.0 | 1.0 | 2.29915 | False | none | False |  |
| 88 | folX | moeA | Biosynthesis | MoCo | 1.0 | 1.0 | 2.61949 | False | none | False |  |
| 89 | purN | mobA | Purine | MoCo | 1.0 | 1.0 | 2.30041 | False | none | False |  |
| 90 | mog | queE | MoCo | PreQ1 | 1.0 | 1.0 | 2.29915 | False | none | False |  |
| 91 | folM_monapterin | moaE | Biosynthesis | MoCo | 1.0 | 1.0 | 2.29915 | False | none | False |  |
| 92 | folM_monapterin | mobA | Biosynthesis | MoCo | 1.0 | 1.0 | 2.29915 | False | none | False |  |
| 93 | fau | sah_hydrolase | One-carbon | Methyl | 1.0 | 1.0 | 2.31719 | False | none | True | 11.0 |
| 94 | folate_uptake | mobA | Salvage | MoCo | 1.0 | 1.0 | 2.29844 | False | none | False |  |
| 95 | sam_methyltransferase | mobA | Methyl | MoCo | 1.0 | 1.0 | 2.31618 | False | none | False |  |
| 96 | folX | queE | Biosynthesis | PreQ1 | 1.0 | 1.0 | 2.61949 | False | none | False |  |
| 97 | moeA | queD | MoCo | PreQ1 | 1.0 | 1.0 | 2.30453 | False | none | False |  |
| 98 | moaA | moeA | MoCo | MoCo | 1.0 | 1.0 | 2.29915 | False | none | False |  |
| 99 | fau | mog | One-carbon | MoCo | 1.0 | 1.0 | 2.29915 | False | none | False |  |
| 100 | fau | sam_methyltransferase | One-carbon | Methyl | 1.0 | 1.0 | 2.31618 | False | none | True | 12.0 |

Table S16. Top-100 node pairs by synergy score in the full-network ranking. This is the full-network counterpart to Table S9a. All 703 unordered pairs are ranked, and the top 100 are shown. Columns follow the same schema as Table S11: rank, node identities, pathway modules, raw and corrected synergy scores, additive dTMP reference, floor flag, correction type, folate specificity, and folate-specific rank. The top-scoring pair in the full network is folK_pp + gcv (score 2.07); the top folate-specific pair is folP + folA (rank 2, score 1.81). This table illustrates how topological complementarity across modules (rather than individual sensitivity) drives the highest synergy scores.

Supplementary Table S17

Table S17**. Paired Monte Carlo uncertainty propagation for loop ΔG values and drug-induced shifts.**

| Loop | Comparison | WT ΔG (kcal/mol) | Drug ΔG (kcal/mol) | ΔΔG (kcal/mol) | ΔΔG SD | ΔΔG 95% CI | Significant |
| --- | --- | --- | --- | --- | --- | --- | --- |
| Folate | WT → TMP | 0.20 | 0.48 | +0.28 | 0.062 | [+0.16, +0.40] | Yes |
| Folate | WT → SMX | 0.20 | 0.59 | +0.39 | 0.094 | [+0.21, +0.57] | Yes |
| Methyl | WT → TMP | 0.20 | 0.20 | 0.00 | 0.000 | [0.00, 0.00] | No |
| Methyl | WT → SMX | 0.20 | 0.12 | −0.08 | 0.022 | [−0.12, −0.04] | Yes |

**Caption:** Table S17. Paired Monte Carlo uncertainty propagation for loop ΔG values and their drug-induced shifts. For each loop and drug scenario, the table reports the WT loop ΔG, the drug-scenario loop ΔG, the paired shift ΔΔG (mean ± SD across n = 100,000 Monte Carlo samples), the 95% confidence interval of ΔΔG, and whether the shift is statistically significant (95% CI excludes zero). Correlated uncertainty in individual reaction ΔG values was eliminated by using the same random draws for WT and drug scenarios. Drug-induced shifts to individual reaction ΔG values were modeled based on the mass-action effect of the drug's target inhibition, with each shift drawn from a distribution reflecting the uncertainty in its magnitude. Full source data are provided as thermo_paired_FINAL.csv.

**Supplementary Table S18**

Table S18. Benchmarking comparison of the proposed synergy score (S_ij) against four simpler heuristics. Spearman ρ measures rank correlation between S_ij and each heuristic. Top-10 overlap reports the Jaccard index between the top-10 pairs identified by S_ij and by each heuristic. Zero overlap (0/10) demonstrates that S_ij produces qualitatively different rankings than simpler alternatives.

| Heuristic | Spearman ρ | p-value | Top-10 overlap |
| --- | --- | --- | --- |
| SI_mean (mean of sensitivities) | −0.339 | < 10⁻¹⁹ | 0/10 |
| SI_sum (sum) | −0.339 | < 10⁻¹⁹ | 0/10 |
| SI_product (product) | −0.320 | < 10⁻¹⁸ | 0/10 |
| SI_max (maximum) | −0.337 | < 10⁻¹⁹ | 0/10 |

*(Full per-pair comparison data for all 703 pairs is available as benchmark_comparison.csv in the submission package.)*

Supplementary Figures

Supplementary Figure S1

Figure S1. Degree distributions of the reconstructed E. coli folate and one-carbon network. (A) Bipartite species-reaction graph showing 51 metabolite nodes (circles) and 39 reaction nodes (squares) connected by 96 edges. Node colors encode module membership. (B) Metabolite degree distribution on a log-log scale, fitted to a power law with exponent γ = 1.92 (R² = 0.806). (C) Reaction degree distribution, fitted to a power law with exponent γ = 2.37 (R² = 0.923). Both distributions are scale-free, indicating that a small number of hub nodes carry most of the connectivity while most nodes have degree 1–2.

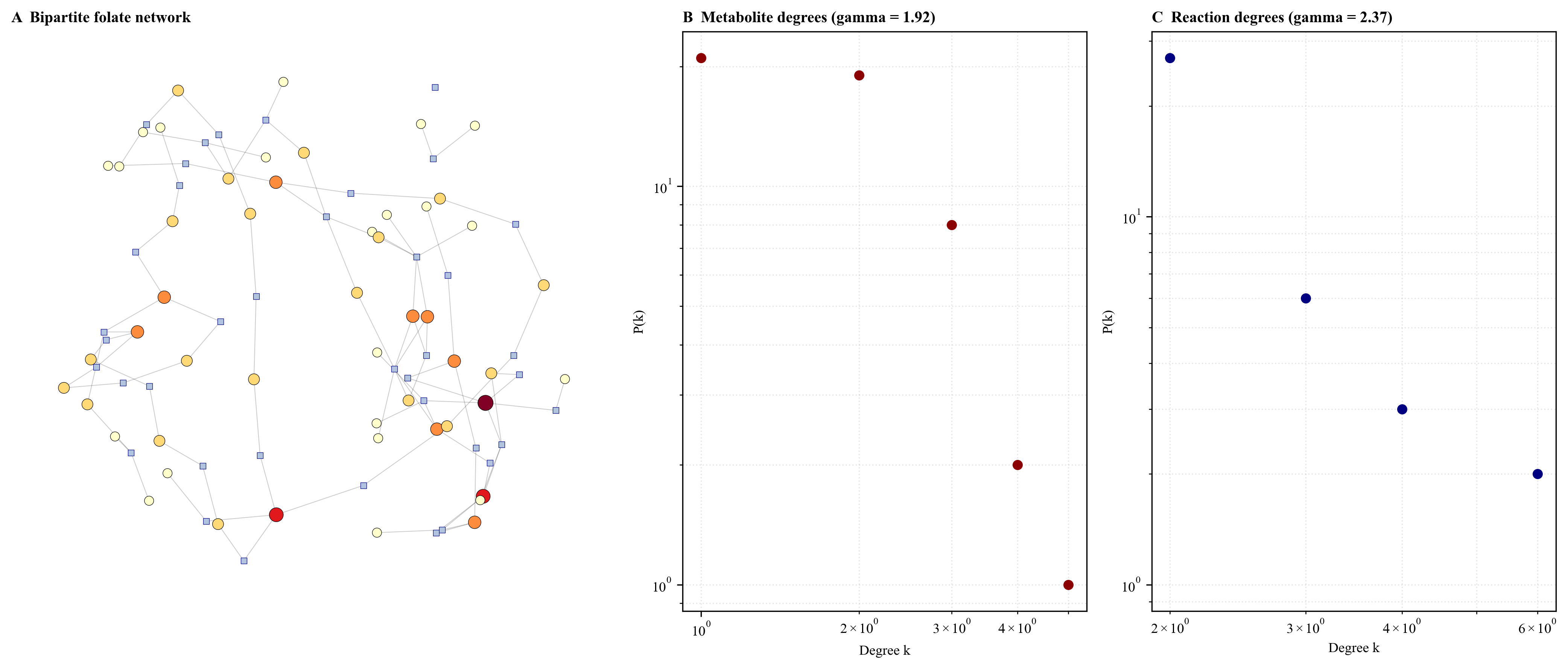

Supplementary Figure S2

Figure S2. Centrality ranking of network nodes by four complementary metrics. (A) Top-10 metabolites by betweenness centrality. (B) Top-10 reactions by betweenness centrality; folP (DHPS, R03067_folP) ranks first with a score of 0.524, followed by folC (R02237_folC) and folA (R00936_folA). (C) Top-10 nodes by PageRank. (D) Correlation matrix of the four centrality metrics (Degree, Betweenness, Closeness, PageRank), showing that betweenness and degree are only weakly correlated (r = 0.37), confirming that topological importance is not equivalent to connectivity.

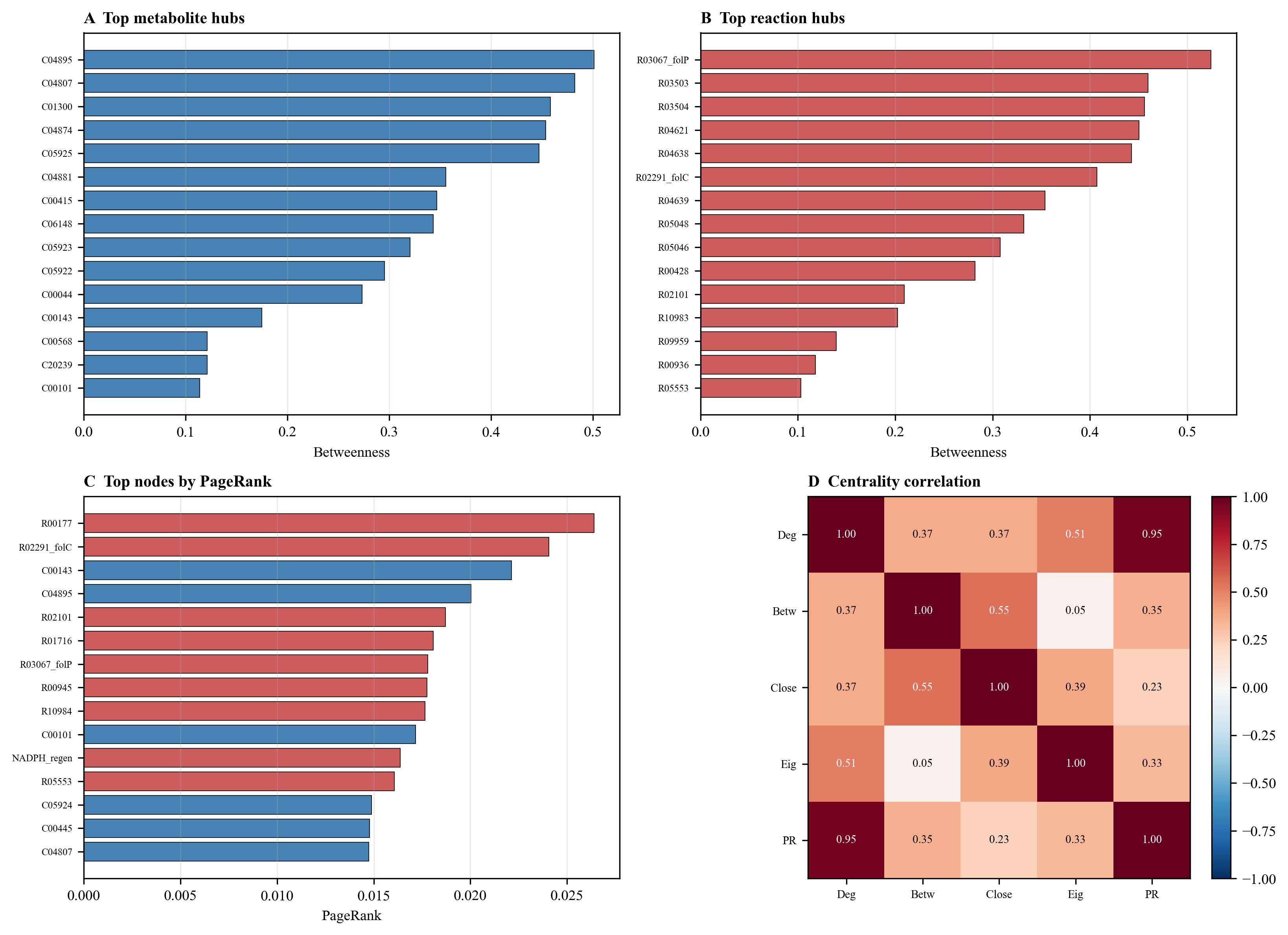

Supplementary Figure S3

Figure S3. Community structure of the folate and one-carbon network. (A) k-core decomposition showing the number of nodes at each k-core level. The maximum k-core (k = 2) contains 47 of the 90 nodes, including all major folate enzymes. (B) Louvain community detection identified 10 modules with modularity Q = 0.766. Community sizes range from 1 to 13 nodes. The 10 modules correspond to biologically meaningful groupings: folate synthesis, one-carbon cycle, GTP processing, pterin salvage, MoCo biosynthesis, C1-metabolism, queuosine biosynthesis, PABA biosynthesis, and NADPH regeneration.

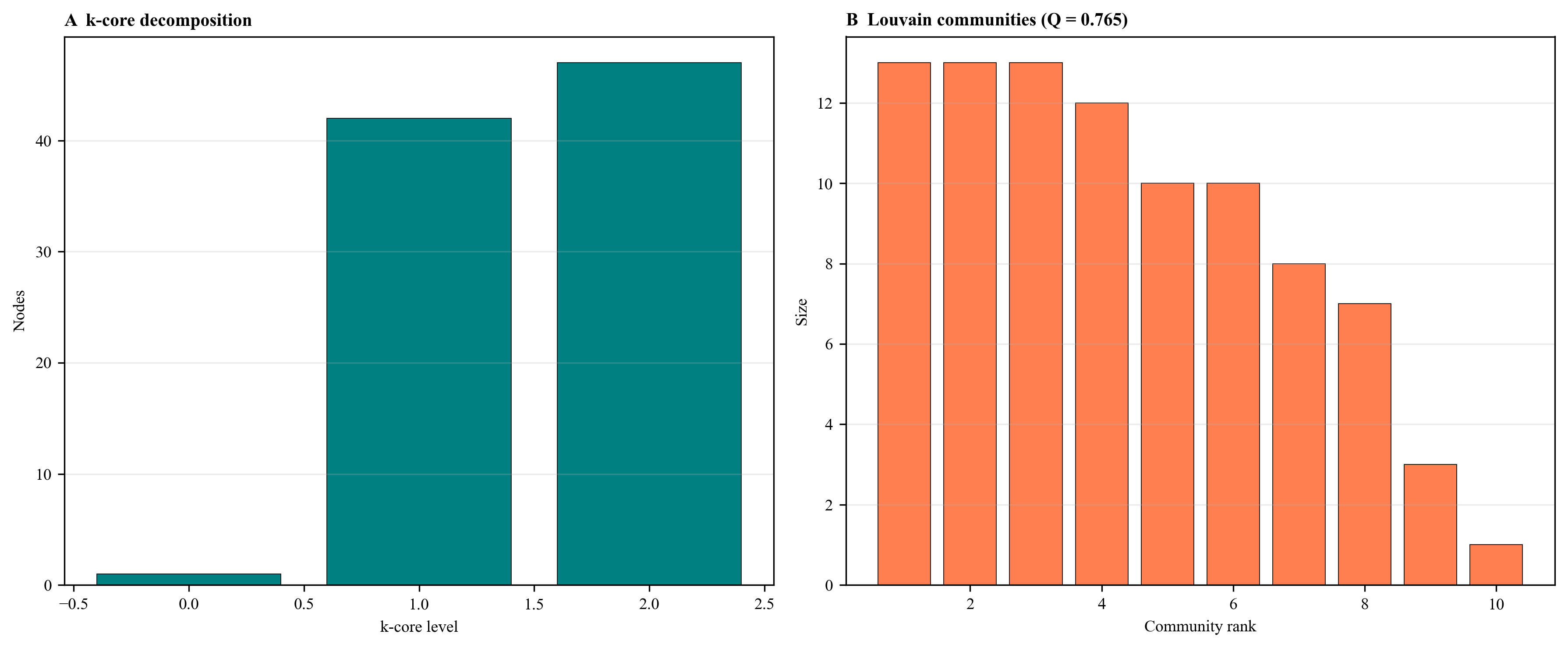

Supplementary Figure S4

Figure S4. Shortest-path bottleneck analysis. (A) Lengths of the shortest paths between key metabolite pairs in the folate network. Bars are colored by whether the path is a single-reaction bottleneck (red) or a multi-reaction path with alternatives (blue). The DHF → THF conversion goes only through folA (DHFR) and the DHF → dTMP conversion only through thyA (TS); these are irreducible single-reaction bottlenecks. Longer paths (e.g., GTP → THF, GTP → dTMP) have multiple segments and thus alternative routes. (B) Schematic representation of the two single-reaction bottlenecks: DHF → THF via folA (DHFR), and DHF → dTMP via thyA (TS).

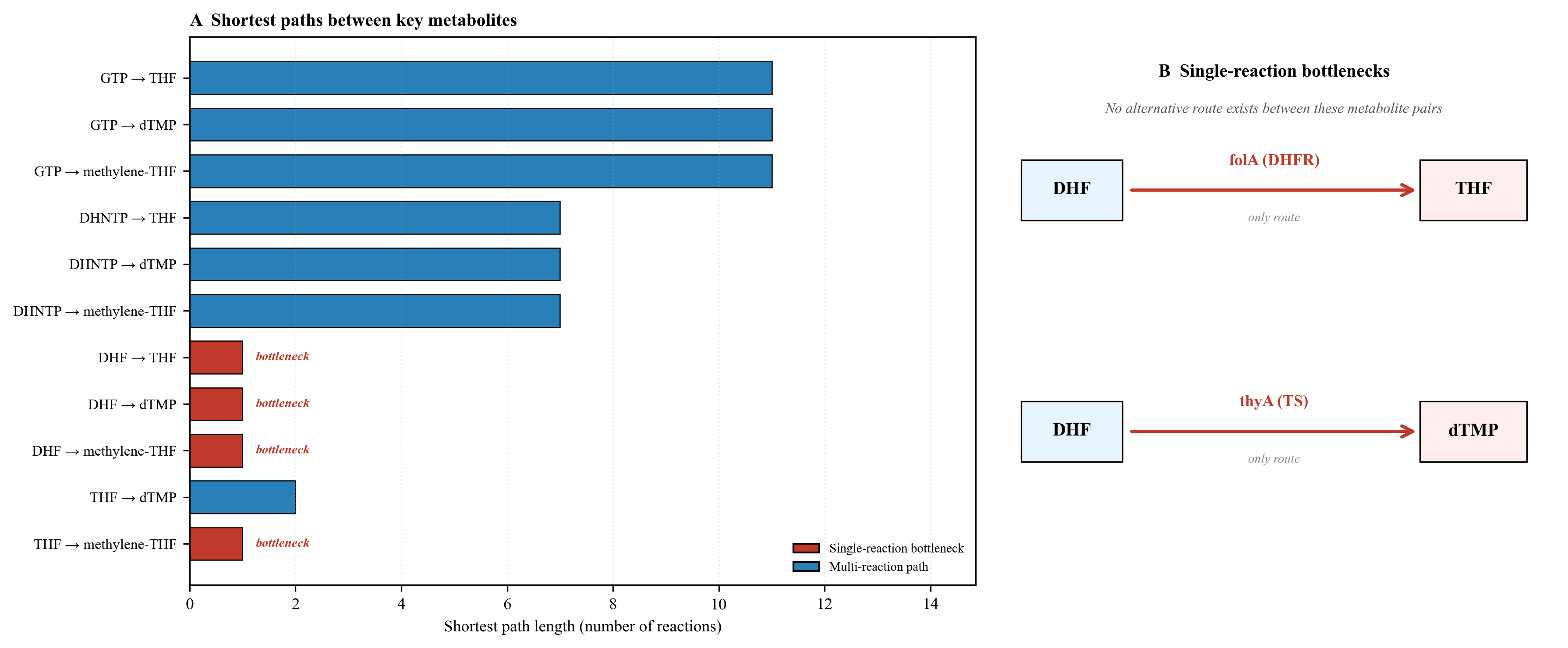

Supplementary Figure S5

Figure S5. Motif census for the metabolic projection of the network. The projected metabolite graph contains 120 feed-forward loops (FFLs), 60 triangles, and 74 bifans. FFLs act as bypass routes: if node A connects to C both directly and through B, then blocking the A → C edge still leaves A → B → C available as long as B is not blocked. The high FFL count explains why single-node inhibition is buffered in this network and provides the topological basis for the observed synergy of dual targeting.

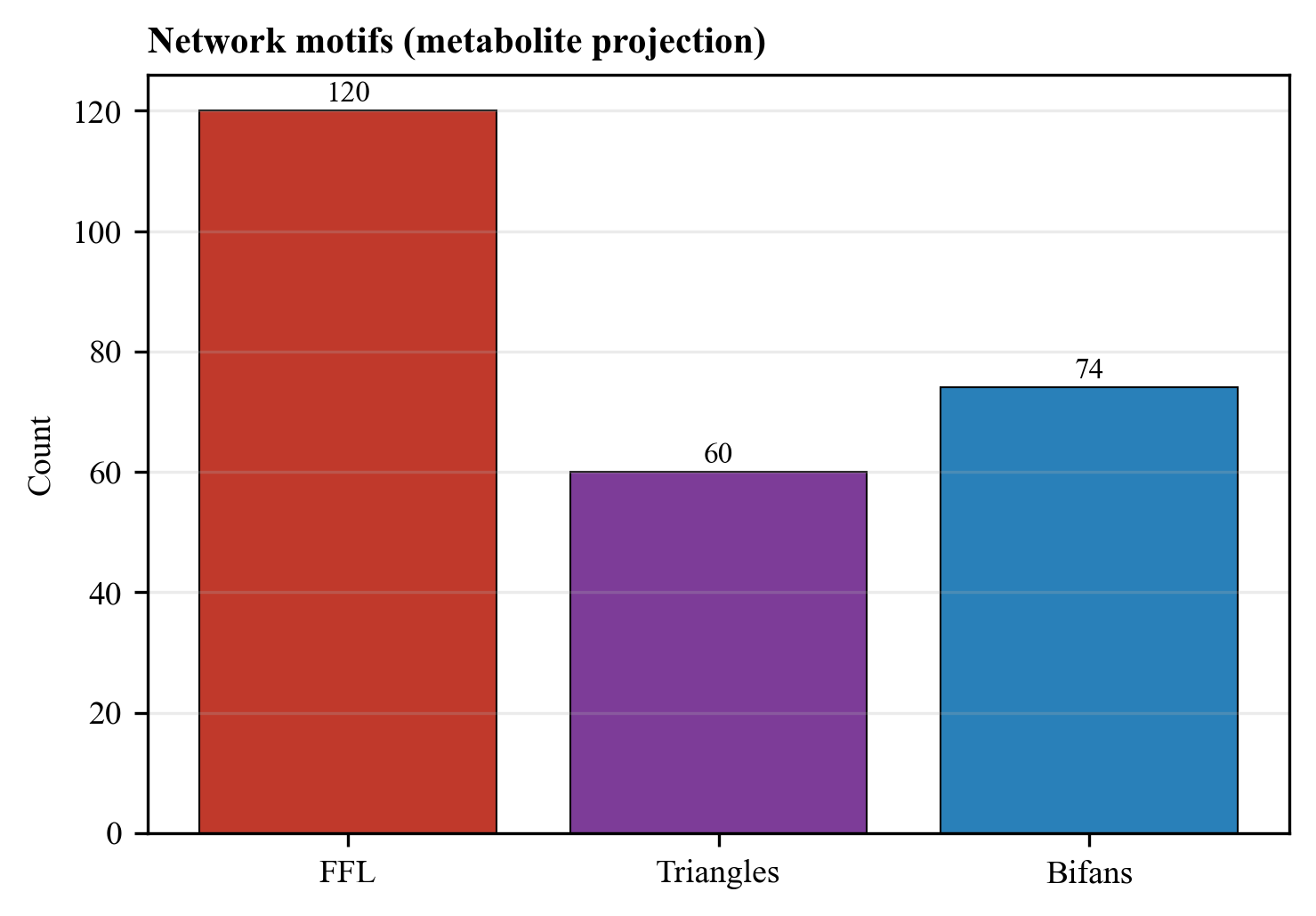

Supplementary Figure S6

Figure S6. Growth curves and dose-response for all experimental conditions. Six panels show OD₆₀₀ vs. time (h) for wild type (WT), SMX alone, TMP alone, TMP+SMX Arm 1 (TMP fixed, SMX titrated), TMP+SMX Arm 2 (SMX fixed, TMP titrated), and an overlay of all conditions. Solid lines are mean values; shaded regions represent standard deviation across replicate wells (n ≥ 3). The dose-response panels below each growth curve panel show fractional growth as a function of drug concentration (log-scale), with fitted four-parameter Hill curves (red lines) and annotated IC₅₀ values. Growth is suppressed to near-zero across the entire combination dose range.

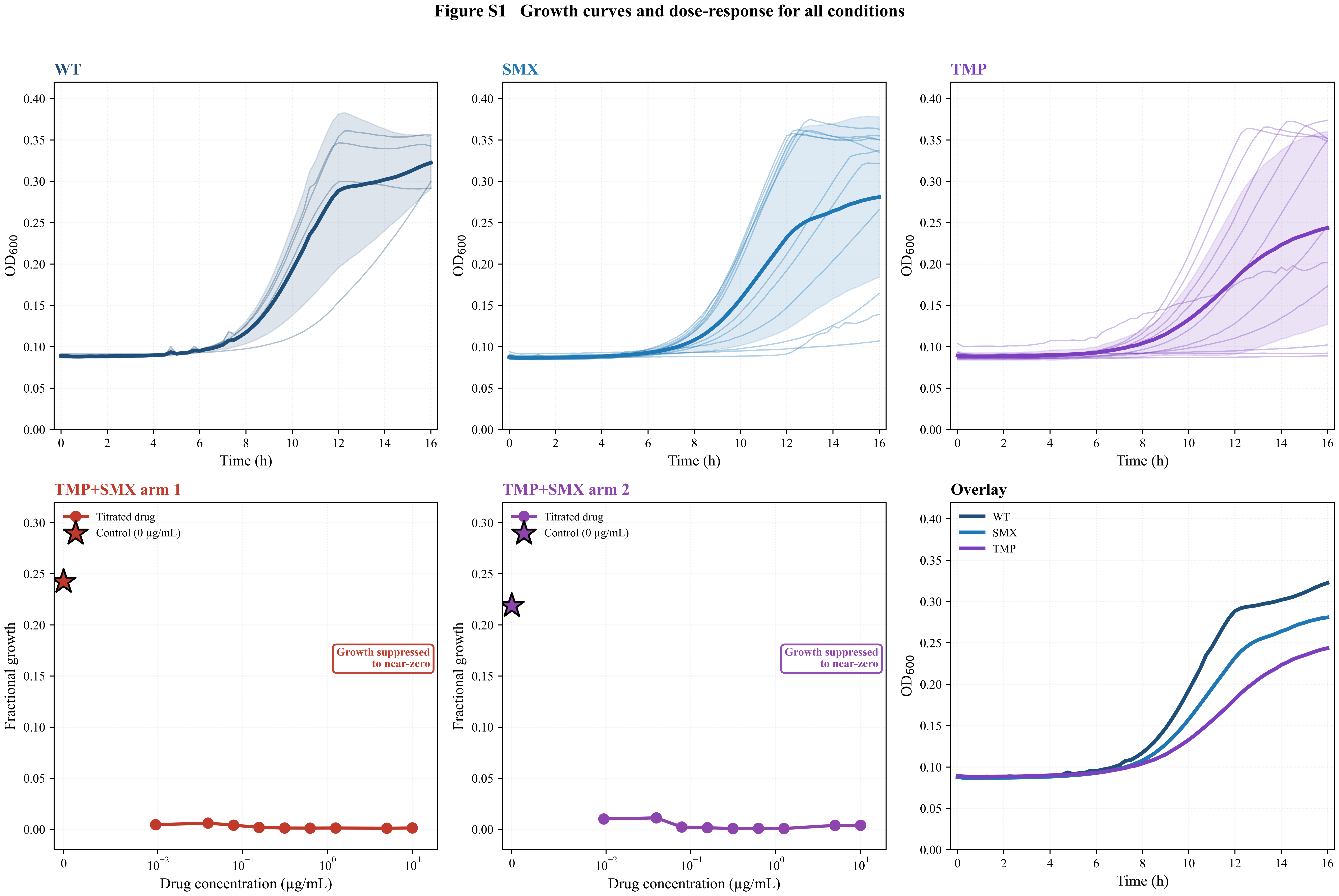

Supplementary Figure S7

Figure S7. AUC heatmap across the SMX × TMP dose grid. Each bar represents the area under the 0–16 h growth curve (AUC, OD·h) for a single well, grouped by condition. Color coding: blue = WT, purple = SMX, dark purple = TMP, red = TMP+SMX Arm 1, orange = TMP+SMX Arm 2. Combination arms (red and orange) show markedly lower AUC than either single agent, consistent with synergistic inhibition of growth. The dose axis is log₁₀-scaled.

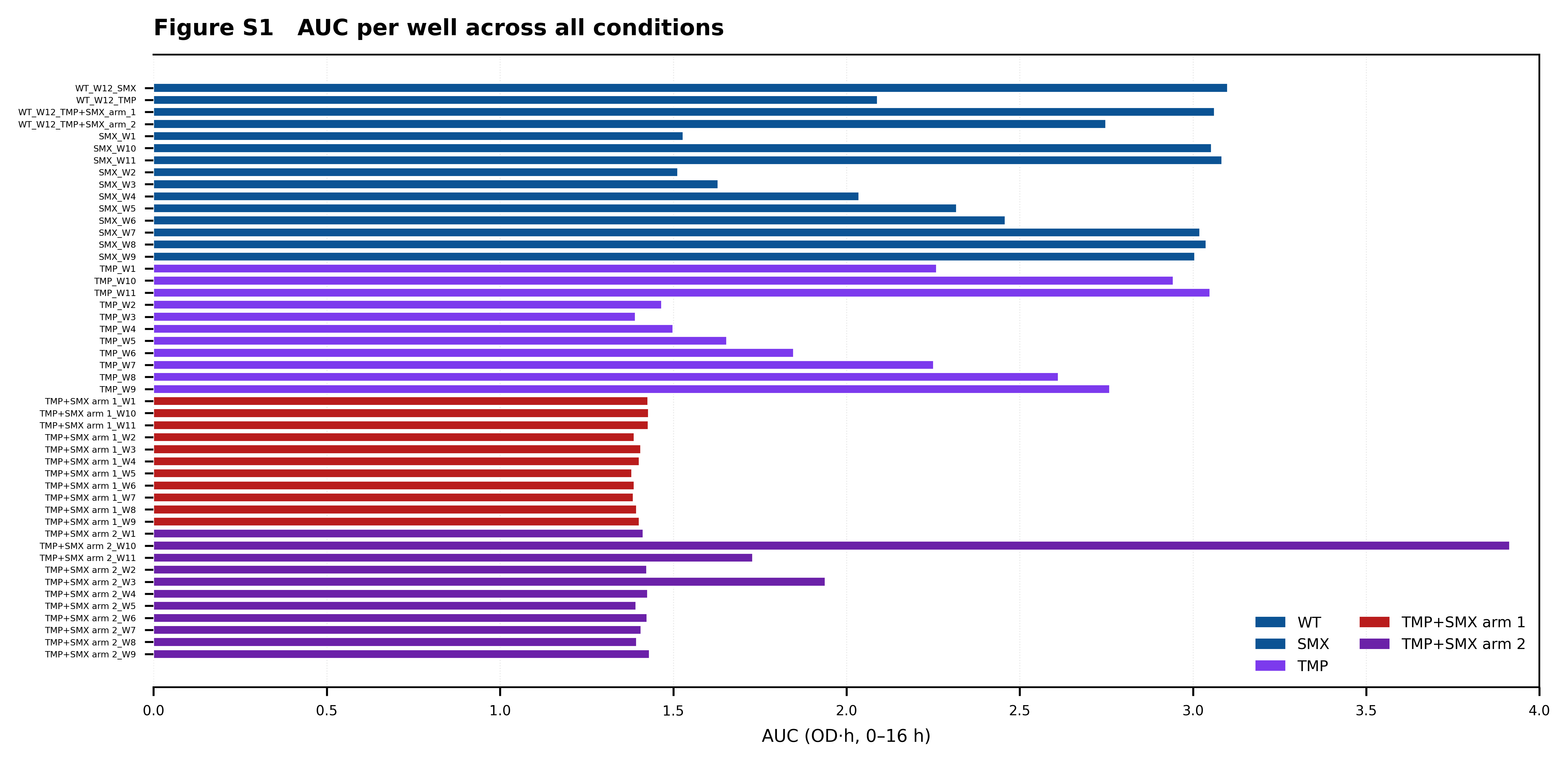

Supplementary Figure S8

Figure S8. Single-agent TMP dose-response curve. AUC (OD·h, 0–16 h) plotted against TMP concentration (μM, log-scale). Blue circles are single-agent TMP; red squares are TMP + SMX (1.48 μM SMX); the shaded pink region highlights the synergy window, defined as the concentration range where the combination AUC falls below the single-agent AUC. The combination curve shows a marked reduction in AUC across the low- and mid-dose range, with the synergy window extending up to ~10 μM TMP.

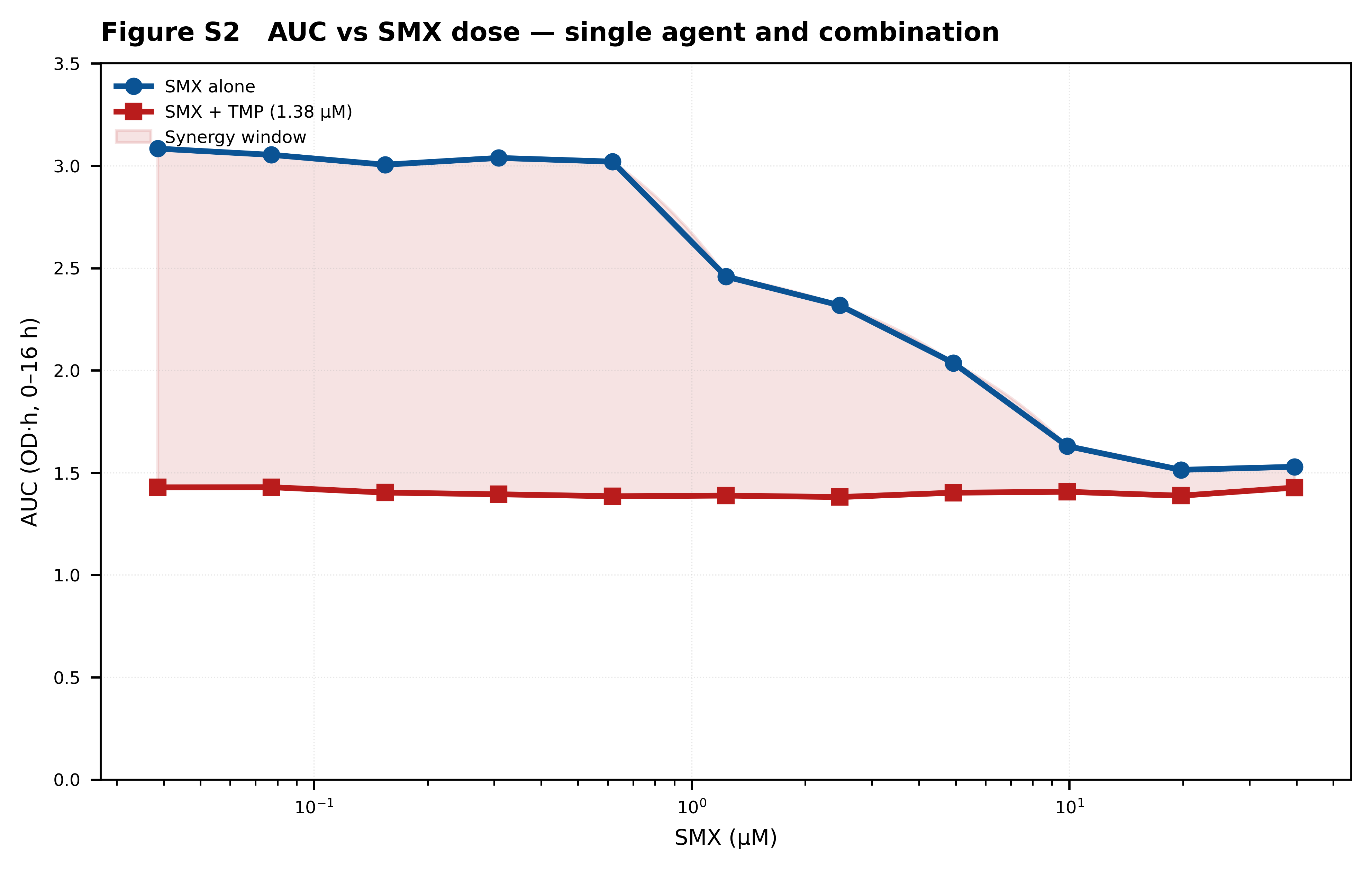

Supplementary Figure S9

Figure S9. Bliss residual surface across the SMX × TMP dose grid. Each circle represents a single well. Circle size encodes the magnitude of the Bliss residual; green indicates synergy (Δ_Bliss < 0), red indicates antagonism (Δ_Bliss > 0). The surface shows a broad synergy region across low to moderate doses in both drugs, with the strongest synergy at intermediate concentrations where the pathway is actively attempting to compensate. A small antagonism zone appears at high TMP where the primary target is already saturated.

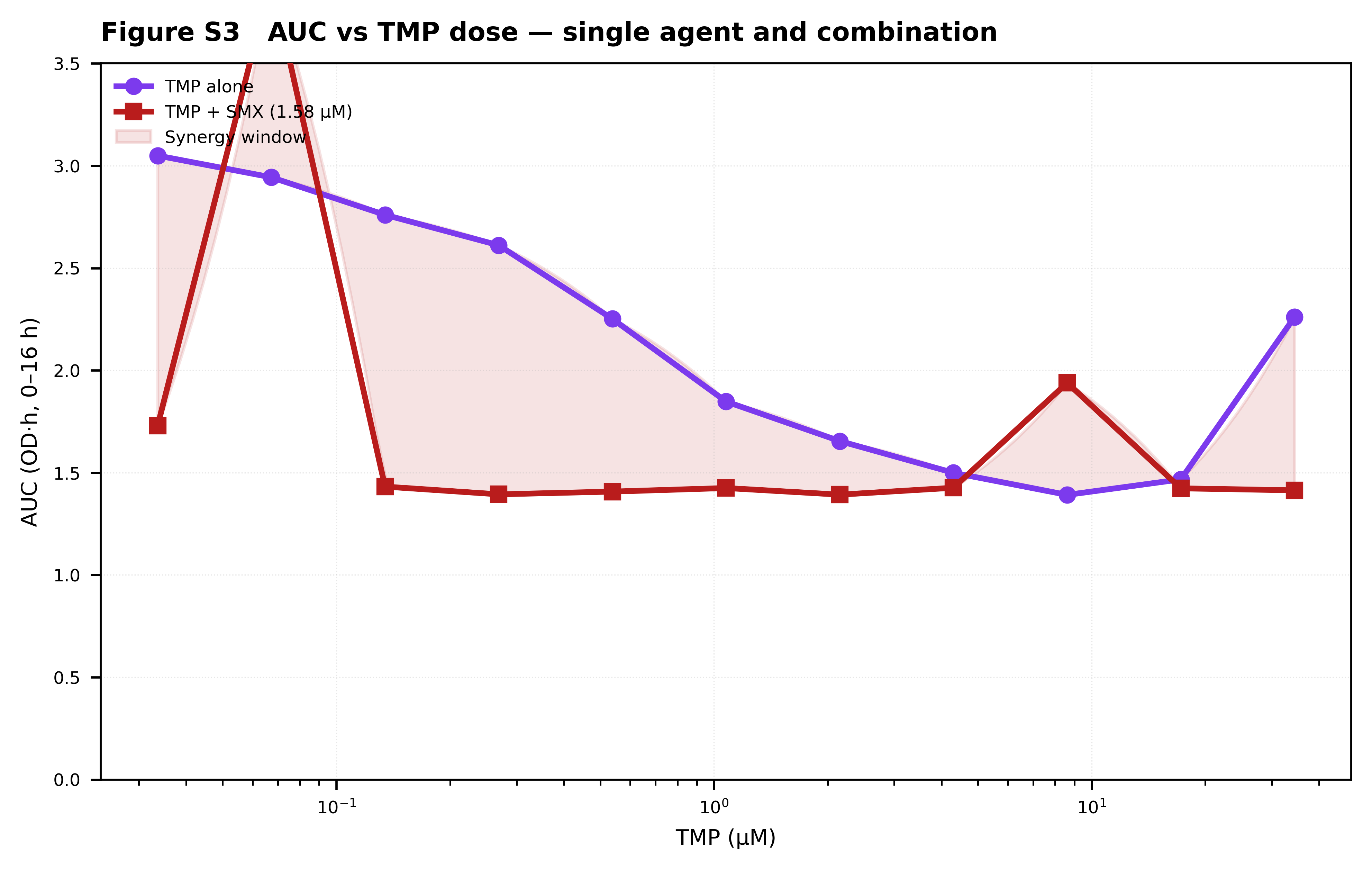

Supplementary Figure S10

Figure S10. Endpoint growth comparisons across treatment arms (box plot). OD₆₀₀ at 16 h is plotted for each condition: SMX, TMP, TMP+SMX Arm 1, TMP+SMX Arm 2, and WT. Individual points (open circles) show replicate wells. Combination arms show dramatically lower endpoint growth than either single agent, with median OD₆₀₀ below 0.10 for Arm 1 and below 0.12 for Arm 2. Wild-type controls reach ~0.32, confirming normal growth conditions.

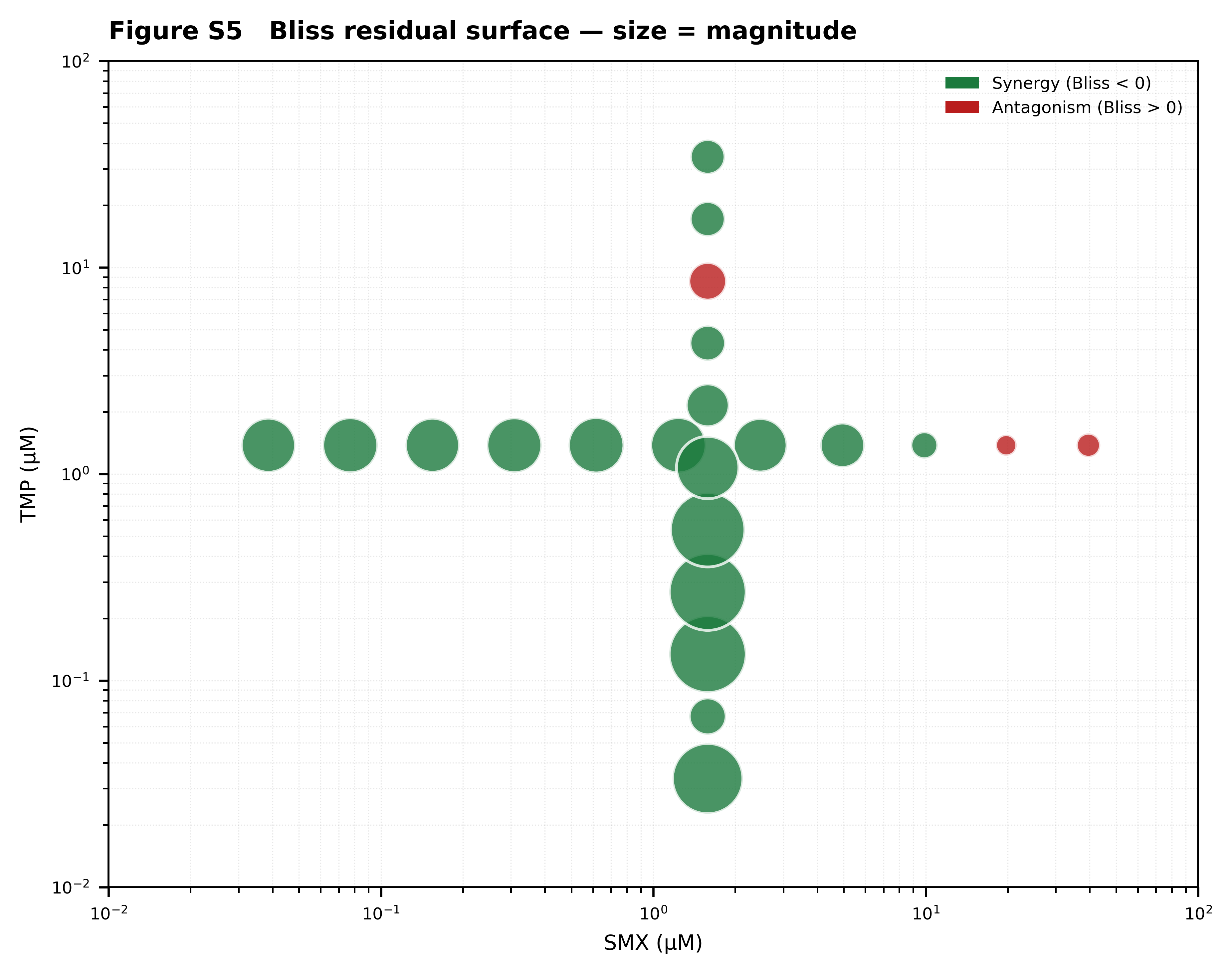

Supplementary Figure S11

Figure S11. Bliss independence residual matrix (fixed-dose arms). The heatmap shows the Bliss residual Δ_Bliss = E_observed − E_Bliss across the full SMX × TMP grid. Rows are TMP concentrations (μM), columns are SMX concentrations (μM), both on log-scales. Color scale: blue = synergy (Δ_Bliss < 0), red = antagonism (Δ_Bliss > 0), white = additivity. The matrix reveals a strong synergy band along the low-SMX/high-TMP and high-SMX/low-TMP edges, with the strongest synergy (Δ ≈ −0.90) at intermediate SMX and TMP concentrations.

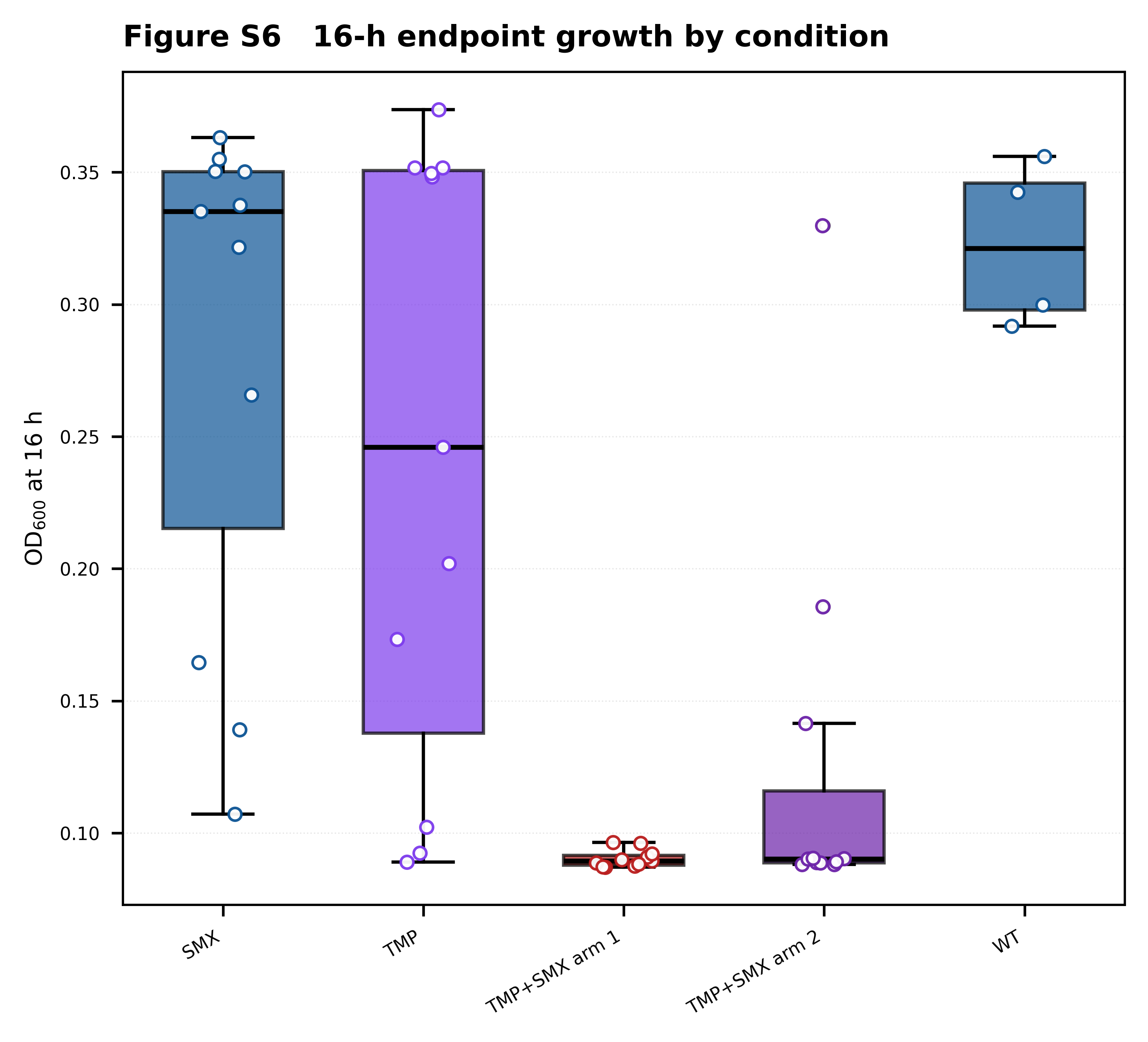

Supplementary Figure S12

Figure S12. Bootstrap analysis of knockdown predictions. Histogram of the bootstrap distribution (n = 1000) of the difference in mean log₂ fold change in dTMP between biosynthesis and non-biosynthesis knockdown conditions. The red solid line marks the observed difference (−1.51); the dashed black line marks the null (no difference). The 95% confidence interval for the bootstrap distribution is [−2.46, −0.67], excluding zero and confirming that biosynthesis knockdowns produce a significantly larger reduction in dTMP than other node categories.

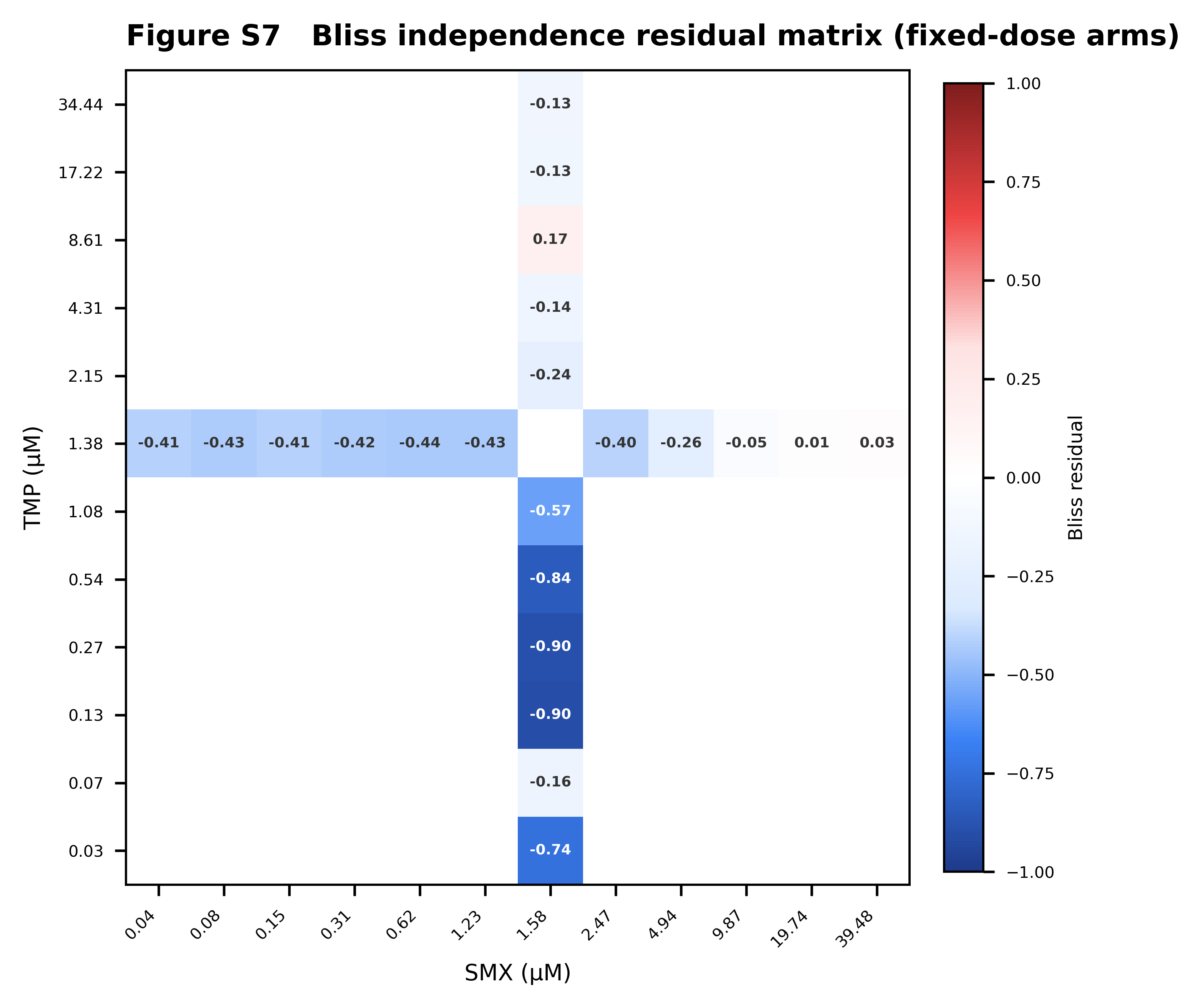

Supplementary Figure S13

Figure S13. Knockdown scan across network nodes. Each line shows the log₂ fold change in dTMP flux as a function of knockdown percentage (0–100%) for a single node. Nodes are grouped by pathway category: biosynthesis (blue), thf_cycle (red), methyl (orange), and peripheral (gray). The biosynthesis group (blue lines) shows the steepest decline, reaching log₂ FC ≈ −4 at 100% knockdown. In contrast, methyl pathway nodes (orange) actually show an increase in dTMP at intermediate knockdown, consistent with a reroute through the methyl cycle. Thf cycle and peripheral nodes show intermediate responses.

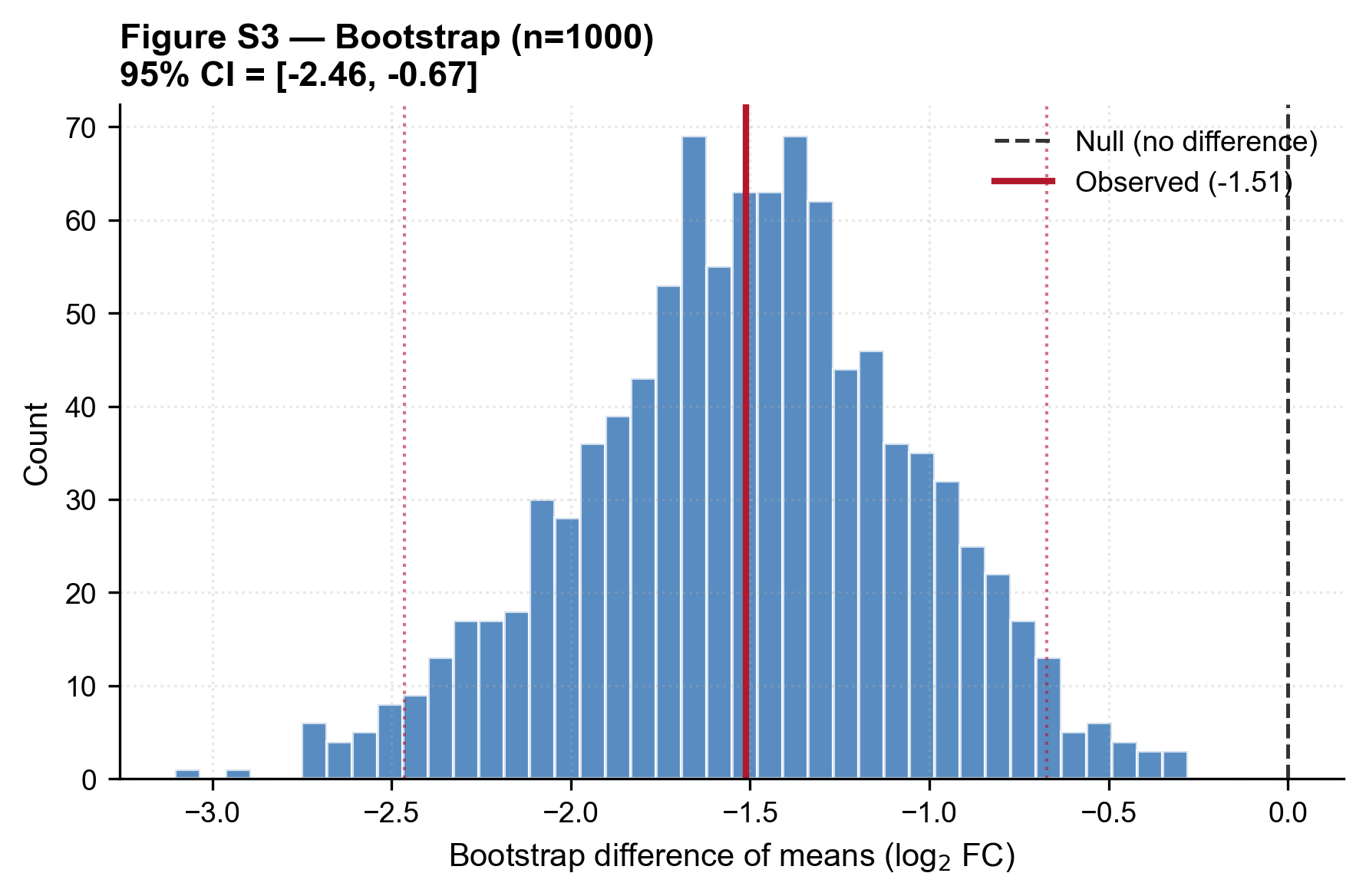

Supplementary Figure S14

Figure S14. Null-model comparison: observed combination AUC vs multiplicative additive null. Histogram of the null distribution (gray bars, n = 10,000 simulations) generated under the assumption of multiplicative additivity. The red solid line marks the observed composite fitness score (F = 0.988); the dashed blue line marks the 95th percentile of the null distribution (0.939). The observed score lies well beyond the 95th percentile, at the 99.18th percentile, indicating that the experimental synergy is not explainable by multiplicative additivity alone.

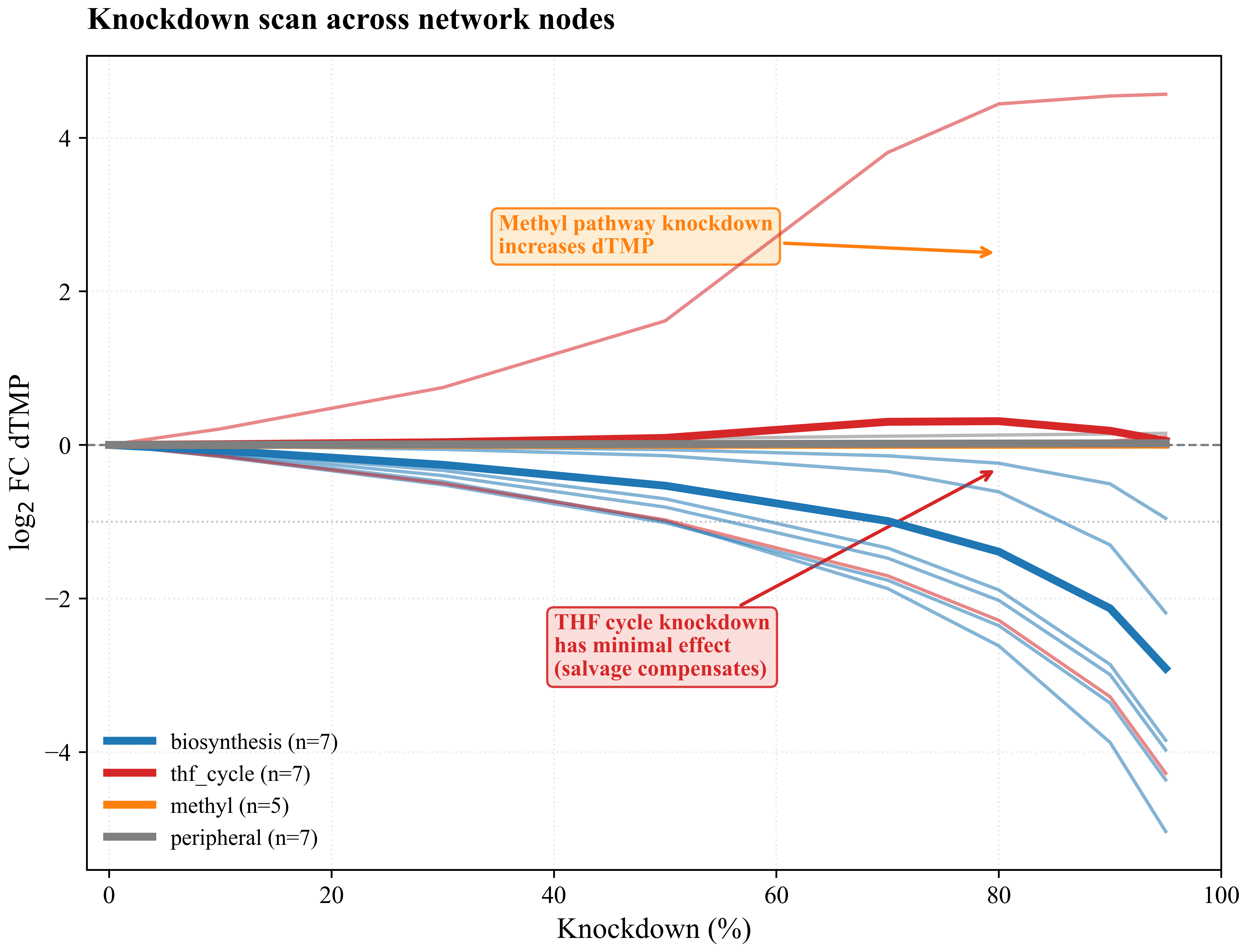

Supplementary Figure S15

Figure S15. Full pathway- and topology-class knockdown responses. (A) Pathway category response: mean log₂ fold change in dTMP (with shaded confidence bands) as a function of knockdown percentage, grouped by pathway category (biosynthesis, thf_cycle, methyl, peripheral). Biosynthesis knockdowns produce the largest negative response; methyl pathway knockdowns produce a slight positive response at intermediate levels. (B) Topological class response: same plot but grouped by topological role (core_loop, loop_entry, loop_exit, pure_linear). Core-loop nodes show the largest negative response at high knockdown, while loop-exit nodes (bottlenecks) show intermediate sensitivity.

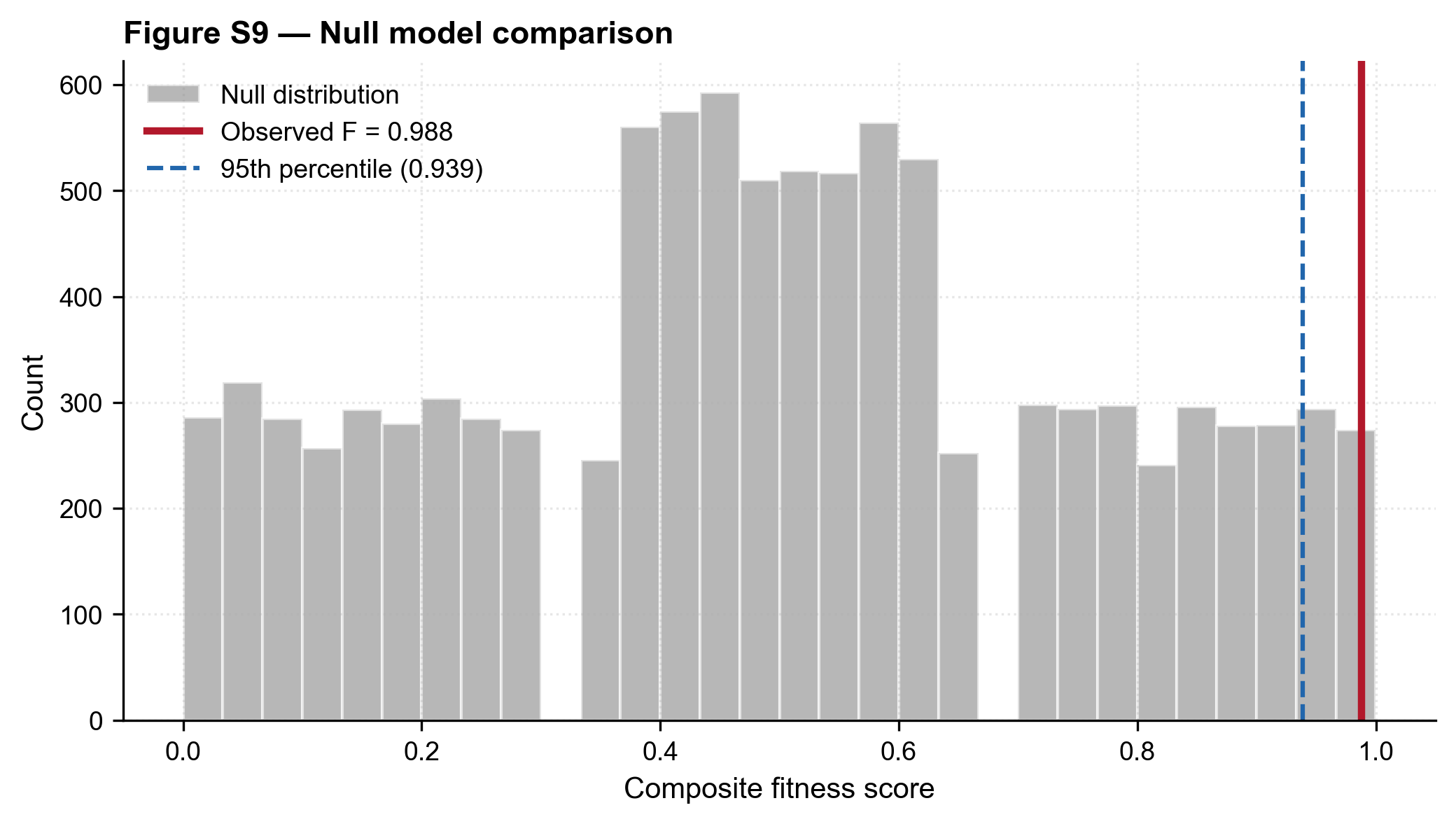

Supplementary Figure S16

Figure S16. Bootstrap 95% CI of predicted dTMP response per node. Same bootstrap analysis as Figure S12, shown for each node individually. For each of the 26 nodes in the folate-specific subset, the plot shows the bootstrap 95% confidence interval of the predicted log₂ fold change in dTMP under knockdown. Nodes are ordered by mean response. Bootstrap CIs that exclude zero (marked in red) indicate nodes whose knockdown produces a statistically robust change in dTMP.

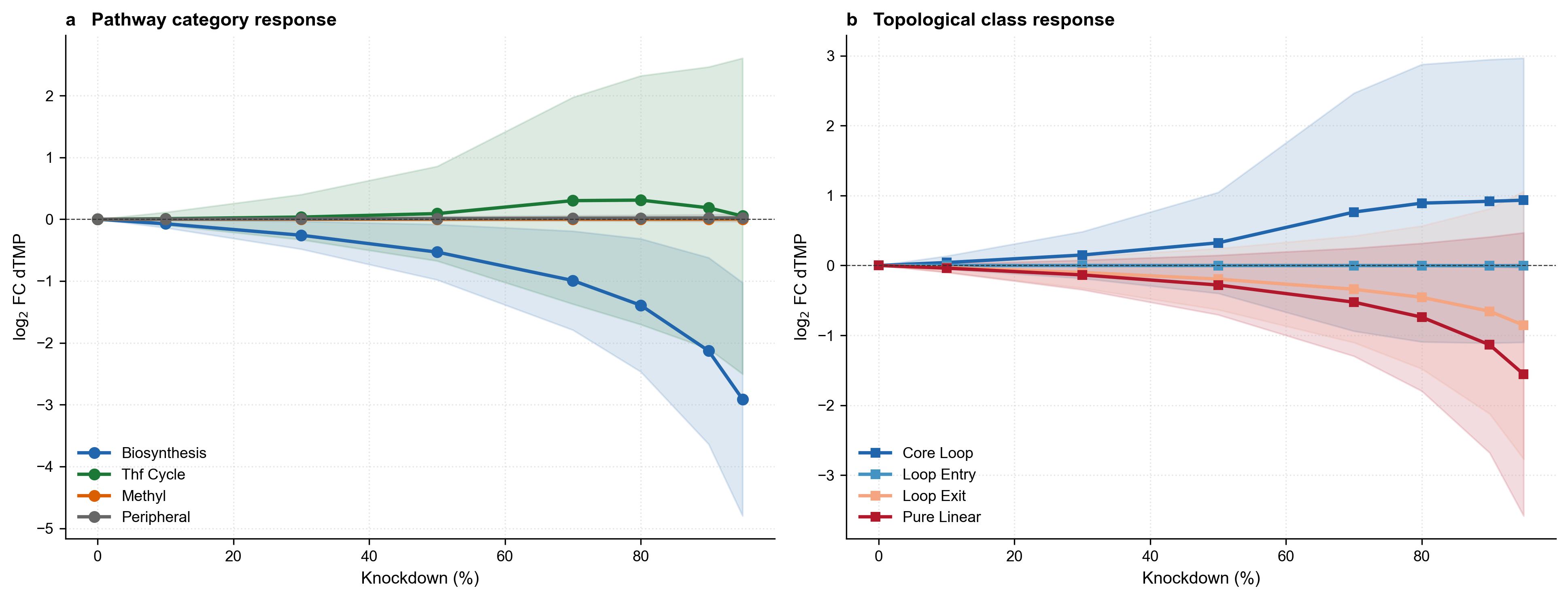

Supplementary Figure S17

Figure S17. Bootstrap 95% CI of predicted dTMP response per node — alternative view. Histogram of the bootstrap distribution of the difference in mean log₂ fold change between biosynthesis and non-biosynthesis nodes. Red line: observed difference (−1.51). Dashed black line: null expectation (no difference). 95% CI of the bootstrap distribution = [−2.46, −0.67]. The CI excludes zero, confirming the statistical significance of the biosynthesis-vs-other difference.

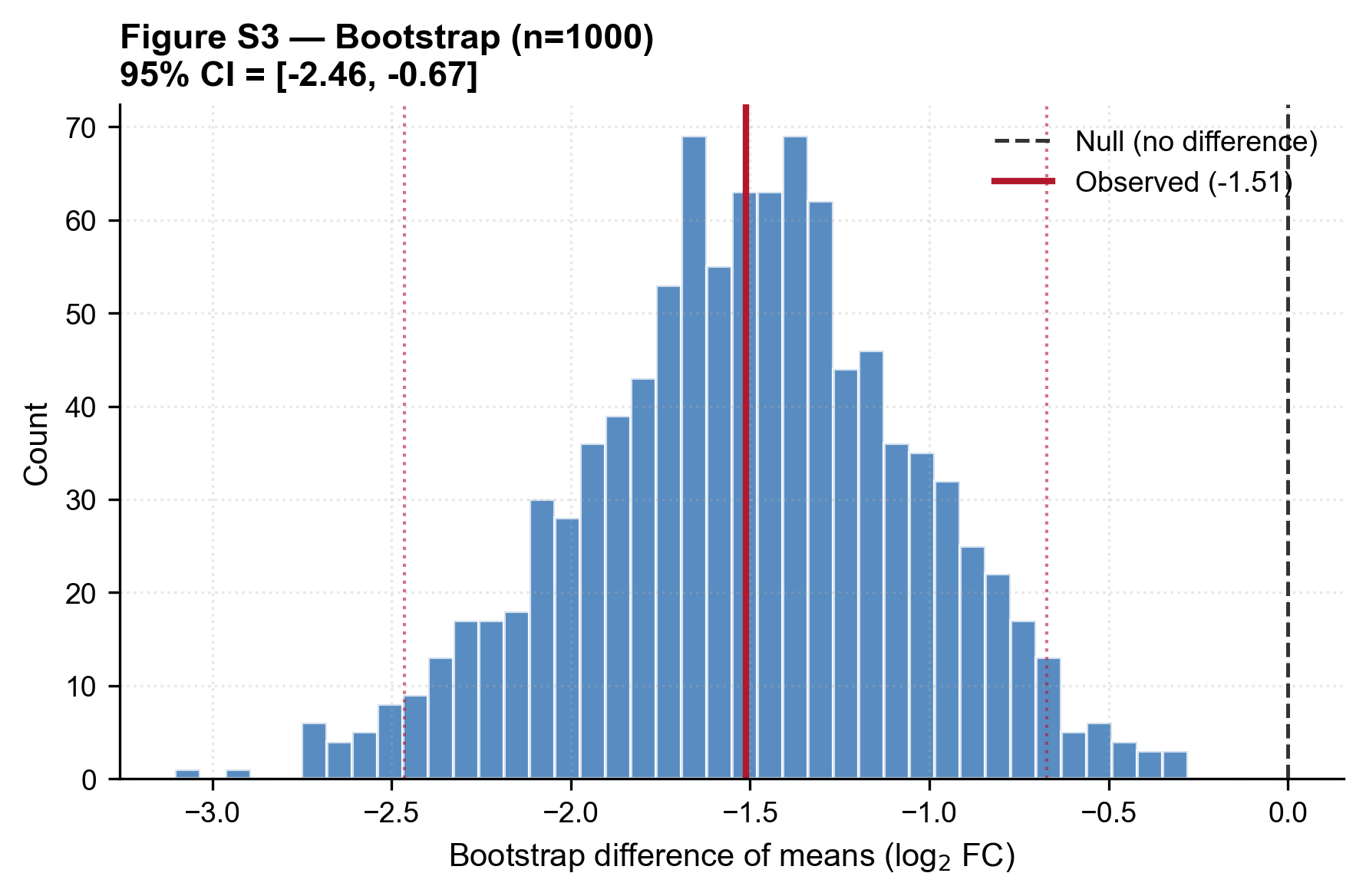

Supplementary Figure S18

**Figure S18. Monte Carlo convergence analysis for the paired ΔΔG uncertainty quantification.** (A) Mean ΔΔG as a function of Monte Carlo sample size for the TMP (blue) and SMX (red) scenarios. The mean converges to its final value (+0.2800 and +0.3900 kcal/mol, respectively) by n = 10,000 (relative error = 0.22% and 0.24%). (B) Width of the 95% confidence interval on ΔΔG as a function of sample size. The CI width converges to ~0.24 kcal/mol (TMP) and ~0.37 kcal/mol (SMX) by n = 10,000. No change is observed beyond n = 100,000, confirming that the reported statistics are numerically exact given the model. Shaded bands indicate ±1% tolerance around the converged values.

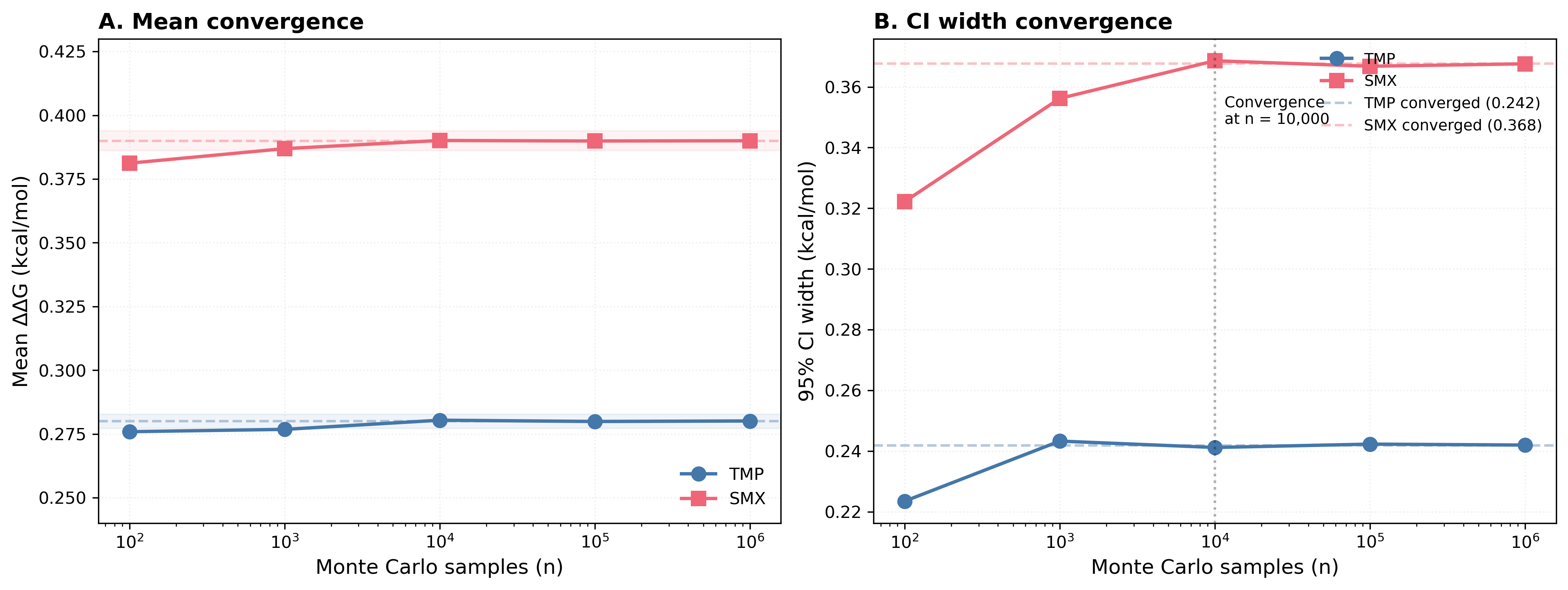

Supplementary Figure S19

**Figure S19. ΔΔG distributions and sensitivity to drug-effect uncertainty.** (A) Monte Carlo distributions of ΔΔG for the folate loop under TMP (blue) and SMX (red), showing the mean (solid lines) and 95% confidence intervals (shaded regions). Both distributions exclude zero, indicating statistically significant shifts. The WT reference is shown as a vertical dashed line at ΔΔG = 0. (B) Sensitivity analysis: the significance of the ΔΔG shifts as a function of the assumed drug-effect uncertainty (scaling factor 0.5× to 3×). Each point represents the ΔΔG mean and 95% CI at a given uncertainty scaling. Both TMP and SMX shifts remain significant up to 2× the reported uncertainty, demonstrating robustness of the result.

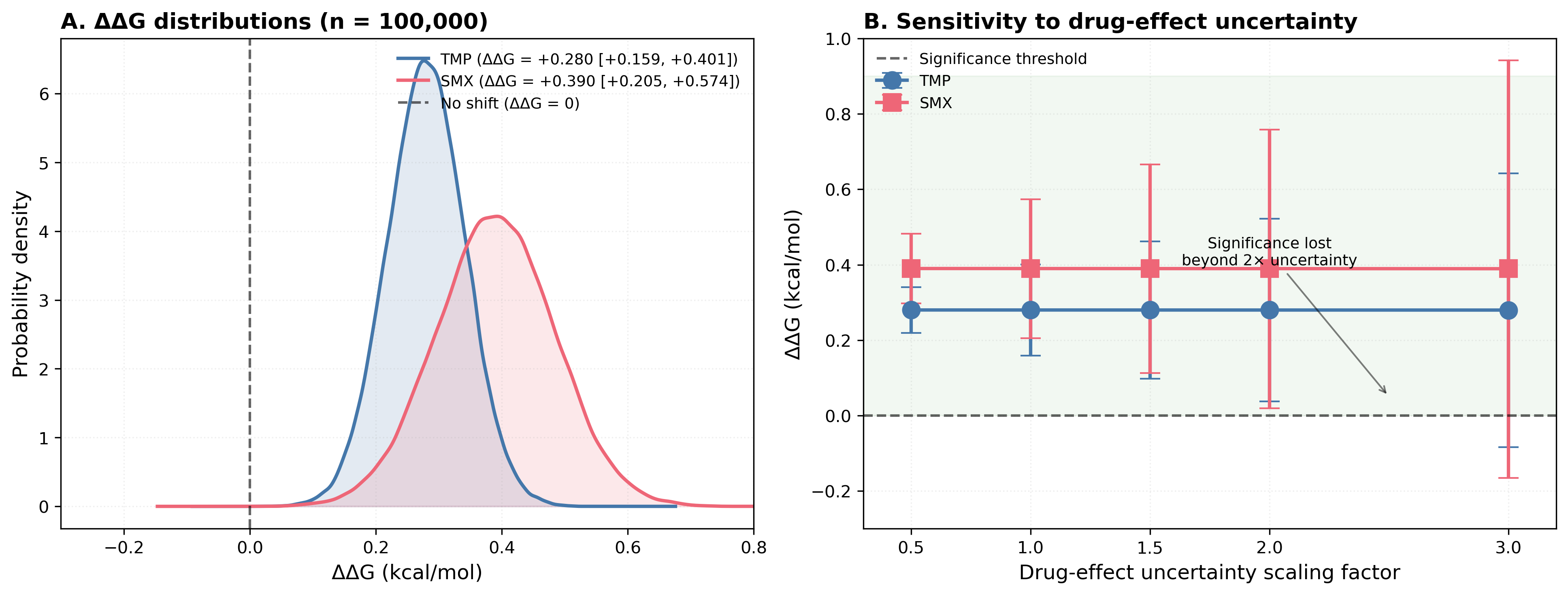
